# *Bonsai*: Tree representations for distortion-free visualization and exploratory analysis of single-cell omics data

**DOI:** 10.1101/2025.05.08.652944

**Authors:** Daan H. de Groot, Sarah X. Morillo Leonardo, Mikhail Pachkov, Erik van Nimwegen

## Abstract

Single-cell omics methods promise to revolutionize our understanding of gene regulatory processes during cell differentiation, but analysis of such data continues to pose a major challenge. Apart from technical challenges such as the sparsity and heterogeneous noise properties of these data, the crucial problem is that we know little about the potentially very complex high-dimensional structures that the data represent. Consequently, there is an urgent need for exploratory analysis methods that allow rigorous representation and visualization of the structure in the data. However, currently popular methods such as UMAP and t-SNE are unsatisfactory because they are *ad hoc*, stochastic, uninterpretable, and known to severely distort the structure in the data.

Here we show that these challenges can be overcome by representing the data on tree structures and present *Bonsai* : a novel method that reconstructs the most likely tree relating any set of high-dimensional objects while rigorously accounting for heterogeneous measurement noise.

We show that, in contrast to other visualization methods, distances along the *Bonsai* trees accurately represent true distances between the objects in high-dimensional space across many types of datasets. Moreover, *Bonsai* automatically regularizes measurement noise, outperforming even methods specifically designed for that purpose on tasks such as nearest-neighbor identification.

By analyzing a blood cell dataset, we show that *Bonsai* trees not only capture known lineage relationships but also provide novel biological insights. For example, *Bonsai* uncovers that different subsets of NK cells derive from the myeloid and lymphoid lineage, and pinpoints genes that distinguish myeloid-NK from lymphoid-NK cells.

*Bonsai* is free from tunable parameters and scales to datasets of hundreds of thousands of cells. The accompanying tool, *Bonsai-scout*, provides visualizations of the *Bonsai* trees and allows for interactive data exploration such as identifying subclades and their markers, visualizing features along the tree, changing the tree layout, and zooming in on substructures. Finally, application to a dataset of football statistics shows the generality of *Bonsai* in successfully capturing complex structures in high-dimensional data.

## Introduction

By providing detailed quantitative and genome-wide information on the states of single cells, single-cell omics methods promise to transform our understanding of the gene regulatory processes underlying development, aging, and tissue regeneration. Indeed, major efforts are underway in which the gene regulatory states of single cells are measured by counting the mRNA copy-numbers of each gene using scRNA-seq [1], or probing the accessibility of the chromatin using single-cell ATAC-seq (scATAC-seq [2, 3]). These efforts have already led to the creation of single-cell *atlases* of model organisms [4–6] and these resources likely contain invaluable information on what kinds of cell states exist, the relationships between them, the genes that distinguish them, and the regulatory circuits that ultimately drive these differences.

However, the analysis of single-cell omics data continues to remain a major challenge toward fulfilling the promise of these experimental methods. Indeed, that a consensus on how to analyze such data is yet to emerge is demonstrated by the astounding proliferation of analysis tools: more than 1750 scRNA-seq analysis tools were published in the last eight years [7], i.e. almost as many as the total number of scRNA-seq papers (Fig S1 and [7, 8]). Apparently, almost every new dataset needs to be accompanied by its own new analysis method to properly present its conclusions.

This raises the question of why it is so hard to analyze these types of data. Part of the answer lies in a number of technical challenges. In particular, single-cell omics data are very high-dimensional, very sparse, and very noisy. Most challengingly, the noise is highly heterogenous, with noise levels varying over several orders of magnitude across measurements. However, a more fundamental problem is that we do not know what type of structure to expect in the data. Distributions of points in high-dimensional spaces can have extremely complex structures for which we lack intuition. We do not even have answers to simple questions such as: Do we expect to find cells in discrete ‘cell types’ corresponding to well-separated accessible subspaces, or do we expect a continuum of accessible states with only small differences in density? Are cells truly constrained to low-dimensional manifolds, as is often assumed, or does the intrinsic stochastic nature of gene regulation cause variability in essentially all dimensions?

Because we do not know what structure to expect or look for, there is an urgent need for methods that can help explore the complex high-dimensional structure of the data by representing both local and global relationships accurately. However, to do this meaningfully, such methods need to visualize the structure in the data in two dimensions without distorting it and without hallucinating structure that does not exist. Unfortunately, currently popular methods such as t-SNE [9] and UMAP [10] are known to fail severely in this regard [11–13].

Because it is well-appreciated in the field that the structures evident in the visualizations produced by tools like t-SNE and UMAP cannot be relied upon, the current practice in the field is to employ these methods almost in a trial-and-error manner [14–16]. In particular, since these visualization methods come with many tunable parameters, they can be tweaked until the visualizations match prior biological knowledge or preconceived expectations, but this of course inherently hinders making truly novel observations or falsifying strongly held beliefs. What is needed are methods for exploratory analyses that can visualize the true structure in the data without distortion and without arbitrarily tunable parameters.

We here show that all these challenges can be overcome by representing single-cell omics data using *tree structures*. First, there are many historical examples where attempts to describe the relationships between high-dimensional objects led to the use of hierarchical representations.

For example, long before the theory of evolution proposed all organisms are related by common descent [17], taxonomists proposed to describe the variety of organisms in hierarchical structures [18]. Trees are used ubiquitously to describe relationships between DNA sequences [19], and hierarchical structures for describing high-dimensional objects have proved effective in machine learning as well [20, 21].

Below we will show that the *n*(*n* − 1)*/*2 true pairwise distances between *n* objects in a high-dimensional space can *generically* be accurately represented by distances along the ∼ 2*n* branches of a tree, meaning that the structure of high-dimensional data can be well-represented by a tree. In addition, since cells from a single organism are related through the lineage tree of cell divisions, we know that gene expression patterns of single cells have in fact diverged along the branches of a tree structure. This implies that a tree representation is also the most natural way of representing lineage relationships between cell states. Finally, since trees can always be displayed in two dimensions, this allows accurate visualization for exploratory analysis without any distortion of the true distances between the cells.

We thus set out to develop a method for representing scRNA-seq data on tree structures. Although many methods have been developed for reconstructing phylogenies, there were several new challenges to overcome. First, we had to develop a probability model for movement through the high-dimensional gene expression space that assigns likelihoods to any possible tree structure (i.e. a topology and branch lengths) given the expression states of cells at the leaves of the tree. Second, we had to develop methods for rigorously accounting for the complex heterogeneous noise properties of scRNA-seq data into this likelihood framework. Third, we had to develop methods to efficiently identify the maximum likelihood tree for a given scRNA-seq dataset with many thousands of cells.

We here present *Bonsai*, a Bayesian method that rigorously solves these challenges from first principles without any tunable parameters. *Bonsai* takes any set of objects with estimated coordinates in a high-dimensional continuous space, together with individual error-bars on each estimated coordinate of each object, and reconstructs the most likely tree with each object at one of the leaves, so that the true high-dimensional distances between all pairs of objects are well approximated by the distances along the branches of the tree. In particular, for scRNA-seq data, the distances between cells along the branches represent the distances between the gene expression states of the cells. In addition, *Bonsai* estimates the most probable gene expression states not only for the cells at the leaves, but also for all internal nodes. Consequently, by following expression changes along the branches, *Bonsai* also automatically infers gene expression trajectories along all lineages in the tree.

*Bonsai* opens up a novel way of exploring cell-to-cell relations which vastly improves on conventional methods. First, we show that in contrast to other visualization methods, *Bonsai* accurately captures cell-to-cell distances in a large variety of realistic simulated datasets, including datasets that do not have an underlying tree structure. In particular, we show that as the dimensionality of the data increases, the accuracy of the tree representation improves generically. Second, we show that by forcing the cells onto a tree structure, in combination with rigorously taking the heterogeneous noise properties of scRNA-seq data into account, *Bonsai* also leads to significantly more accurate estimates of the true positions of the cells in gene expression space. In particular, *Bonsai* dramatically improves nearest-neighbor identification over standard methods.

Application to real data shows *Bonsai*’s power to uncover new biological insights. Rather than just clustering cells into different ‘celltypes’, the reconstructed tree fully resolves which cells are more or less similar on all scales, identifying global lineage relationships. Indeed, using a blood cell dataset as an example [22], we show that *Bonsai*’s tree not only recovers known lineage relationships between cell types, but identifies novel lineage relationships as well.

Finally, in recent years, experimental progress has led to the collection of high-dimensional datasets across a number of fields in science including cytometry [23], metagenomics [24], population genetics [25], neurobiology [26], and imaging data [27]. We propose that *Bonsai* is applicable to data from all these domains and expect it to be a powerful tool for exploratory analysis across all these types of data. To illustrate *Bonsai*’s general ability to rigorously capture the structure of any high-dimensional dataset with continuous features, we apply it to a dataset with statistics of professional football players, showing that it not only accurately identifies groups of players with different roles in the game, but also automatically identifies exceptional players.

Apart from providing *Bonsai* as a stand-alone tool, we have also implemented an automated pipeline for scRNA-seq analysis as a webserver at `bonsai.unibas.ch` that starts from an mRNA-count matrix, normalizes the data using Sanity [28], identifies clusters of statistically indistinguishable cells using cellstates [29], and then runs *Bonsai* to reconstruct a tree relating the cellstates. Finally, to facilitate exploring *Bonsai*’s results, we also provide *Bonsai-scout*, an interactive app that can be used to view the tree using different layouts, to zoom in on parts of the tree, to define clusters of cells by finding maximally-separated clades, to overlay gene expression on the reconstructed tree, and even to find marker genes that most distinguish different clades of cells. Example results of our integrated scRNA-seq pipeline for two cell atlases of human and mouse [4, 5] are provided as community resources.

## Results

The key problem faced by methods for visualization and exploratory analysis of high-dimensional data is that there is no way of placing points in a two-dimensional image such that the distances between these points accurately reflect the distances between the points in the original high-dimensional space. A major advantage of tree representations is that representing distances along the branches of the tree provides much more flexibility in representing distances while maintaining the ability to visualize the tree in two dimensions. However, while there are *n*(*n* − 1)*/*2 distances between a set of *n* points, a binary tree has only 2*n* − 1 tunable branch lengths, so that in general a tree should not be able to to accurately capture all pairwise distances.

An obvious exception is when the process by which the objects have diverged from each other can itself be described by a tree. Indeed, the ubiquitous use of trees to represent relationships between DNA sequences is justified by the fact that these have diverged along evolutionary lineages that can generally be described by trees. For collections of cells from a single organism we similarly know that the transcriptomes (or epigenomes) have diverged along the lineage tree of cell divisions. Therefore, it is natural to use trees to represent the lineage structure of single-cell omics data.

### The blessing of dimensionality

But what if the data do not derive from a tree structure? To what extent can the 2*n* − 1 branches of a tree capture the *n*(*n* − 1)*/*2 distances for a set of objects that did not diverge along a tree? To test this, we created datasets in which 100 points were placed at random distances from the origin and at random directions in either 2, 10, 100, 1000, or 10 000 dimensions. We represented these datasets using PCA [30], UMAP [10], and *Bonsai*, and then calculated the correlations between the true distances and those in the visualizations (Fig S2). Strikingly, we find that while increasing dimensionality makes conventional visualization methods deteriorate quickly in performance, the accuracy of tree structures keeps improving as dimensionality increases. Where PCA deteriorated from a perfect correlation to an *R*^2^-value of 0.20, UMAP went from an already weak correlation to no correlation at all. In contrast, *Bonsai*’s visualization improved from an *R*^2^ of 0.51 to an almost perfect correlation in 10 000 dimensions (*R*^2^ = 0.999). Thus, at least for these random scatters of points, the higher the dimensionality, the better all pairwise distances can be represented by a tree.

We next tested whether this observation extends to real scRNA-seq datasets by processing a dataset of cord blood cells [22] using Sanity [28] and then treating the gene expression states that Sanity estimates as a ground truth dataset. We created datasets with different dimensionality by selecting 10, 100, 100, or 10’000 genes and tested to what extent the distances along the *Bonsai* representations of these datasets matched the true distances between the cells. Strikingly, we again find that as the dimensionality of the datasets increases, the tree representations perform better and better in capturing pairwise distances (Fig S3).

We thus find that, from the point of view of representing the structures of complex datasets, rather than a curse there may be a ‘blessing of dimensionality’: the higher the dimensionality of the data, the better all pairwise distances can be represented by distances along the branches of a tree.

### *Bonsai* reconstructs the maximum likelihood tree structure for single-cell transcriptomics datasets

Since we are going to focus on the analysis of scRNA-seq data, we will in the following assume that the objects of our dataset are cells and their positions in high-dimensional space are their gene expression states, even though the arguments apply much more generally.

For a given dataset *D*, consisting of measurements of the gene expression states of a collection of cells, we need to define a model that assigns likelihoods *P* (*D*|*T*, ***t***) of a tree, as defined by its topology *T* and branch lengths ***t***. The optimal representation of the dataset then corresponds to the tree that maximizes this likelihood. Very generally, the likelihood can be decomposed into factors corresponding to the movement in gene expression space along the branches of the tree, and factors for the probability of the measurements given the assumed positions of the cells at the leaves of the tree:

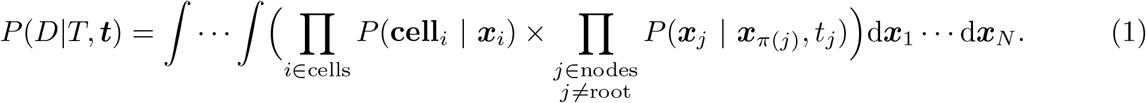

Note that in this definition, we are marginalizing over the true positions in gene expression ***x***_*j*_ of each node *j*. In SI.B.1, we describe the derivation of this likelihood expression in detail. Briefly, the tree has two types of nodes: leaf-nodes corresponding to observed cells in the dataset, and internal nodes that are inferred as intermediates on the trajectories between cells. These can for example be interpreted as likely ancestor cells in developmental settings, or as possible intermediate cell states in a cellular reprogramming context.

The first product in the likelihood only runs over leaf-nodes, and *P* (**cell**_*i*_ | ***x***_*i*_) is the probability of the observed data for cell *i* assuming true gene expression state ***x***_*i*_. *Bonsai* assumes that the likelihoods *P* (**cell**_*i*_ | ***x***_*i*_) can be approximated as a product of independent Gaussians for each component of the feature vector ***x***_*i*_, with independent standard-deviations for each feature (which may vary across cells as well). For example, for scRNA-seq data we need to provide a most likely position in the high-dimensional gene expression space, with independent uncertainty estimates for each gene in each cell (see Fig 1**A**). In our applications these are obtained directly from the normalization method *Sanity* [28].

**Figure 1.**
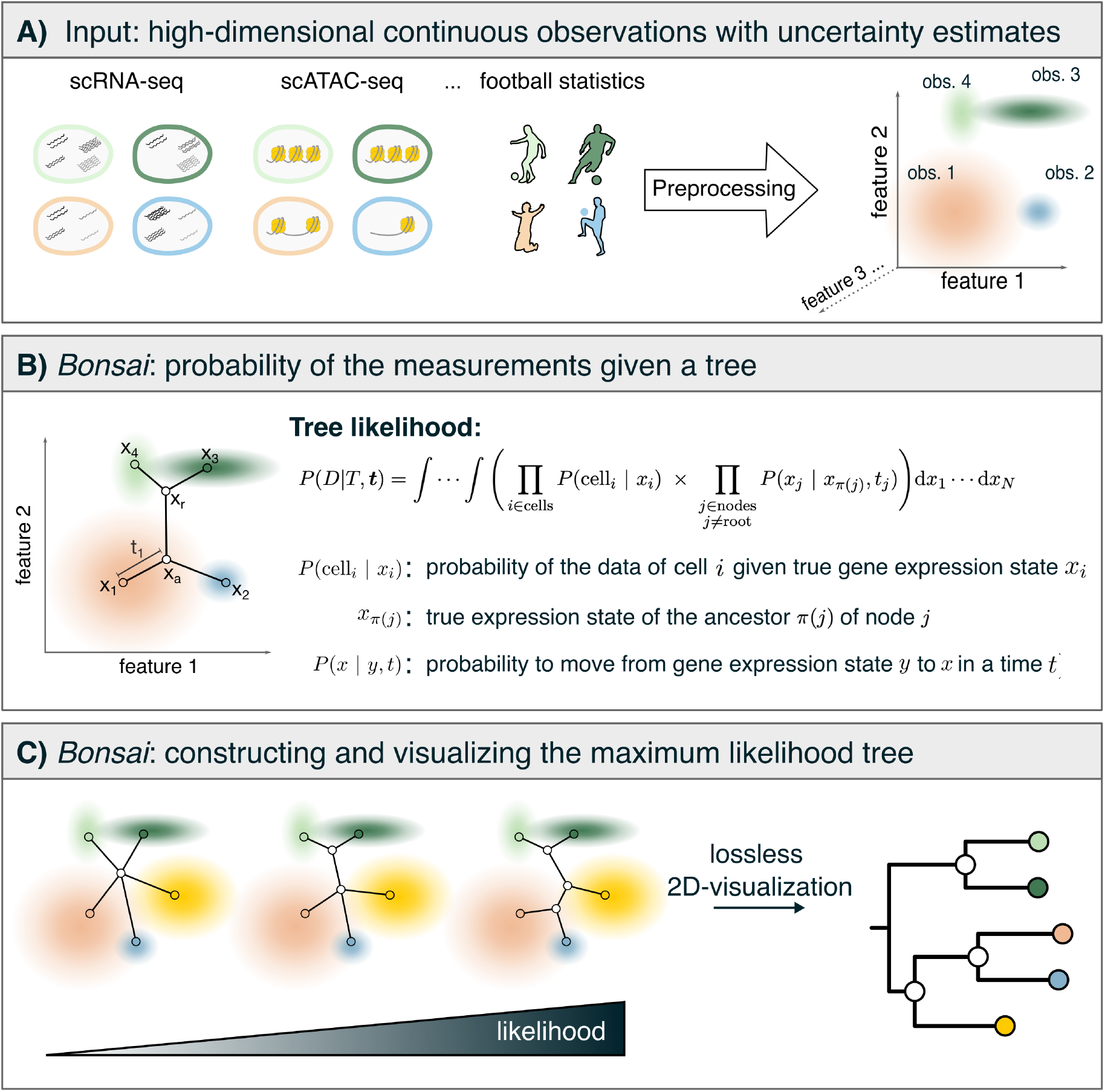
A graphical summary of the *Bonsai* tree representation method. **A)** Raw data of any type should be preprocessed such that, for objects that are near each other, their Euclidean distances meaningfully reflect how dissimilar they are. For example, for scRNA-seq data, we define gene expression states by the output of the *Sanity* algorithm, which corrects for both biological and technical sampling noise [28]. The input to *Bonsai* is a vector of features defining the most likely position of each object, plus independent uncertainty estimates for all feature values. **B)** Structure of the likelihood calculation for a proposed tree. **C)** *Bonsai* searches the space of all trees by iteratively adding internal nodes, to find the tree that maximizes the likelihood. This tree can always be visualized in two dimensions.

The second product corresponds to the probabilities for the movement in gene expression space along each of the branches of the tree. In particular, if node *j* is connected by a branch of length *t*_*j*_ to its ancestral node *π*(*j*), then *P* (***x***_*j*_ | ***x***_*π*(*j*)_, *t*_*j*_) corresponds to the probability to move from the gene expression state ***x***_*π*(*j*)_ of the ancestor, to state ***x***_*j*_ of node *j*.

Determining which movements in gene expression space are more or less likely boils down to making assumptions about the dynamical process that governs gene regulation, which we know relatively little about. In fact, a main reason for gathering scRNA-seq data is to learn about this process in the first place. We will thus take a Bayesian approach where we assume as little as possible about which movements in gene expression space are more or less likely, and let the data guide us to the most likely tree structure. In turn, this tree structure will then provide information on the true gene expression dynamics.

More specifically, the minimal assumption that we make to determine the probabilities *P* (***x***_*j*_ | ***x***_*π*(*j*)_, *t*_*j*_) is that movement in gene expression space can be described by a continuous Markov process, i.e we expect no discontinuous jumps in gene expression, and we expect changes in state to only depend on the current gene expression state and not on how the cell got to this state.^1^ Moreover, we will not impose any preferred directionality for the gene expression dynamics. In SI.B.1.4, we show that based on classical results in Markov process theory [31,32] this constrains our prior to being a homogeneous diffusion process so that the conditional probabilities *P* (***x***_*j*_ | ***x***_*π*(*j*)_, *t*_*j*_) become Gaussians as well. This allows us to perform all the integrals over the positions ***x***_*i*_ analytically, obtaining an analytical expression for the likelihood of any tree (see SI.B.2). Very roughly, as discussed in the Methods, one can think of the log-likelihood of a tree topology under our model as being proportional to the sum of the *logarithms* of the lengths of its branches.

Given this likelihood function, *Bonsai* searches the space of all trees to maximize the likelihood of the data (Fig 1C). Since the number of possible trees grows super-exponentially with the number of cells [33], it is computationally infeasible to perform an exhaustive search. Instead, we developed a search algorithm inspired by phylogenetic methods [34–37] which we describe in detail in the Methods section and in SI.C. Coarsely, we start from a *star-tree* which consists of one root to which all cell nodes are connected. From that tree, we iteratively add an internal node so as to maximally increase the likelihood, and we stop the iteration when the likelihood can no longer be increased. After this, we continue to search the tree space for a better solution by trying local changes to the tree topology using so-called Nearest Neighbor Interchanges.

After finding the optimal tree, *Bonsai* reports its topology *T* and branch lengths ***t*** in the standard Newick format. To visualize and interactively explore the resulting trees, we also provide an application, called *Bonsai-scout*, that allows different visual layouts [19] and a variety of downstream analyses.

### *Bonsai* representations conserve cell-to-cell distances on all scales

To test to what extent *Bonsai* produces representations that maintain the structure in realistic scRNA-seq datasets, we simulated seven scRNA-seq datasets of 1024 cells with known ground truth positions and then sampled raw UMI counts to match statistics from real scRNA-seq datasets including the distributions of total UMI counts, mean expression levels across genes, variances in expression levels across genes, and so on (see SI.E for a detailed description). For six of the seven datasets, we assume that the ground truth gene expression of the cells diverged along the branches of a tree. In the first dataset this is a simple binary tree with all branches of equal length and each gene’s expression diffusing independently along each branch of the tree. In the subsequent five datasets we added increasing complexities including unbalanced trees with some clades containing many more cells than others, random branch lengths from a wide distribution, and using a realistic covariance matrix (mimicking observed correlations in expression across genes so that the diffusion along the tree’s branches effectively occurs in a lower dimensional subspace).

Finally, we also created a non-treelike dataset by simulating seven ‘pseudobulk’ clusters with sizes ranging from 34 to 320 cells. Within each cluster the cells were given almost identical ground truth states, but their raw data differ because their UMI counts were sampled independently from Poisson distributions.

We find that *Bonsai* successfully captures both the local and global structure in all these datasets, presenting a huge improvement over commonly used visualization tools such as Principal Component

Analysis (PCA) and Uniform Manifold Approximation and Projection (UMAP) (Fig 2).

**Figure 2.**
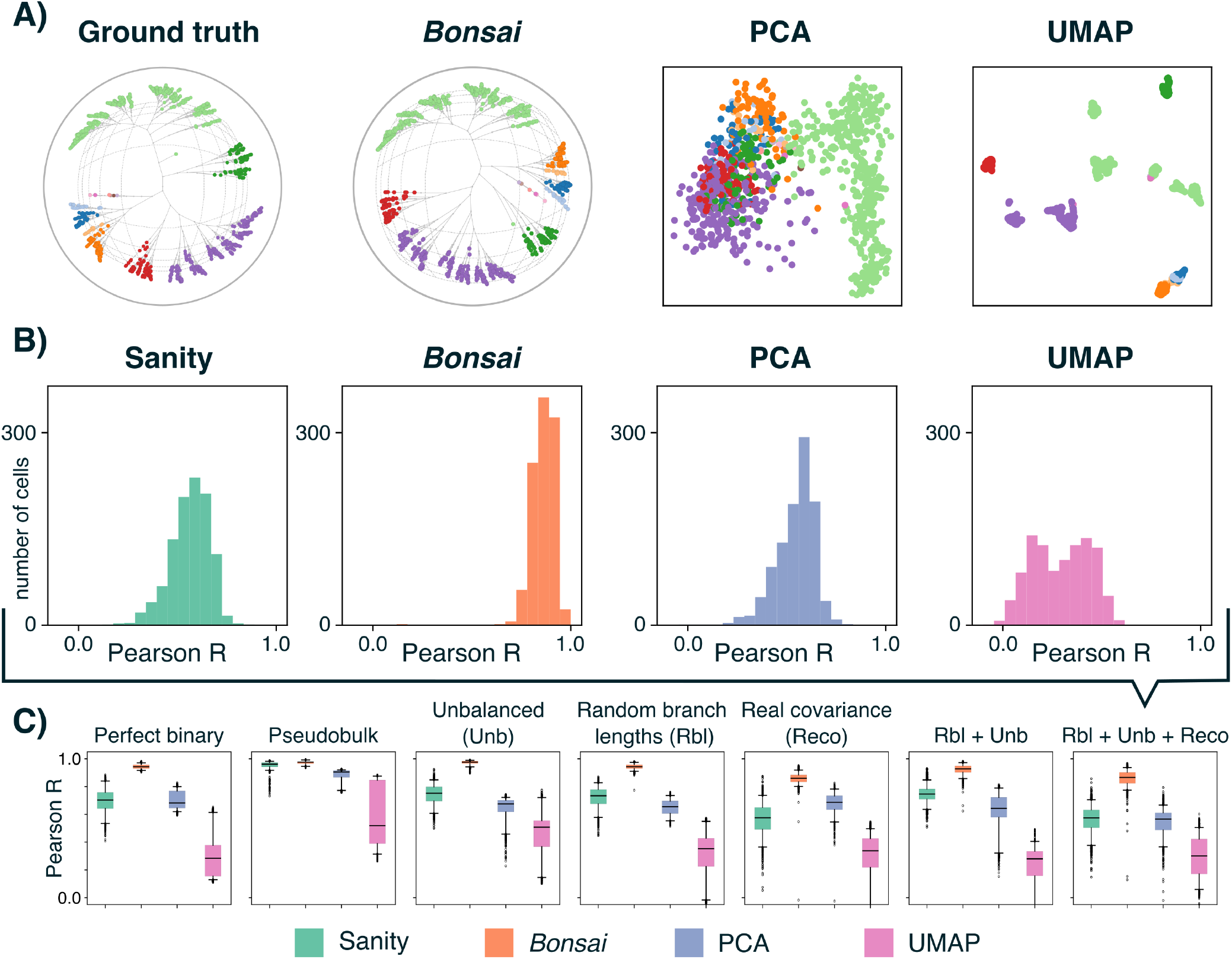
*Bonsai* tree representations accurately capture cell-to-cell distances. **A)** Comparison of the ground truth distances in the most complex synthetic tree-like scRNA-seq dataset (left), with the visualizations of this dataset by *Bonsai* (second from left), 2D-PCA (second from right), and 2D-UMAP (right). The different colors indicate the 16 clades created by the first number of branchings in the ground-truth tree. Both the ground truth tree and the *Bonsai* -reconstruction are plotted on the hyperbolic disk, which naturally shrinks all distances near the edge of the disk. This effect is illustrated by the equal-sized squares on the background that appear to shrink away from the center. **B)** Histograms of the correlations (Pearson-R) between the true distances from each cell to all other cells, and the corresponding distances in each of the visualizations; the closer the correlations are to 1, the better the distances in the visualization match those in the ground truth. **C)** Boxplots of the same distributions of correlations for all seven simulated datasets (see S4-S9 for details). *Bonsai* consistently outcompetes conventional visualization tools independent of whether we 1) take an unbalanced tree (Unb), 2) take random branch lengths (Rbl), and 3) diffuse in a lower-dimensional subspace (Reco). *Bonsai* also gives a superior visualization even if the data did not derive from a tree but rather using 7 pseudobulk-clusters from a real dataset.

Figure 2A compares the visualizations of the most complex dataset, in which the ground truth is an unbalanced tree with significantly varying branch lengths, and in which the true expression levels are correlated across genes in a manner mimicking real data. Whereas *Bonsai* almost perfectly recovers the structure of this dataset, the PCA and UMAP visualization fail almost entirely on this task. For example, although UMAP clusters some groups of cells together correctly, it completely fails to represent the distance structure between or within these groups, and tuning its parameters does not solve this problem. As perhaps could be expected, the PCA visualization only manages to capture the rough global structure along the two dimensions with most variation, i.e. separating the light green clade of cells from the others along the first axis, and separating the purple and red from the blue and orange clades along the second axis.

To quantify the extent to which the true cell-to-cell distances in a dataset are captured by each representation, we calculated, for each cell, the correlation between the true distances to all other cells and the corresponding distances in the representation. Figure 2B shows histograms of these correlations for the dataset of Fig 2A, and Fig 2C shows boxplots of the distributions of correlations for all 7 datasets. Given that these synthetic datasets are all heavily affected by the same Poisson sampling noise as real scRNA-seq datasets, it is remarkable that *Bonsai* manages to accurately capture almost all pairwise distances across these datasets (see Figs S4-S9 for more details). These results suggest that, in contrast to existing methods, *Bonsai* can accurately capture both the local and global structure in scRNA-seq datasets.

### *Bonsai* representations outperform existing methods on nearest-neighbor identification

Many scRNA-seq analysis methods employ so-called *k*-Nearest-Neighbor graphs, where cells are connected to their *k* nearest neighbors in gene expression space. However, we have previously shown that, due to the large measurement noise in scRNA-seq data, it is typically highly challenging to correctly identify the true nearest-neighbors of each cell [28], and we developed a method that explicitly uses the error-bars provided by the Sanity normalization to improve on nearestneighbor identification. It is therefore striking that, as shown in Fig 2B-C, *Bonsai* also substantially outperforms Sanity’s distance estimates.

To investigate this further, we compared *Bonsai*’s performance on identifying nearest-neighbors with those of Sanity, PCA, and UMAP on all 7 synthetic datasets, and find that *Bonsai* dramatically outperforms these methods, including the specialized distance estimation method that comes with *Sanity* [28] (Fig S10). This is remarkable, not only because the inferences of *Bonsai* and *Sanity* are based on the same preprocessed values and uncertainties, but also because we previously showed that *Sanity* vastly outcompetes other methods on this task [28]. This unparalleled accuracy most likely arises because reconstructing a tree forces the estimated distances to be consistent between nearby cells on the tree, so that information about similar cells is automatically taken into account for each distance estimate.

To confirm this, we started from the dataset with 100 randomly placed points in a 100-dimensional space of Fig S2 and then created different datasets by taking either 1, 2, 5, 10 or 20 noisy ‘measurements’ of each of the 100 points. We find that, in contrast to the PCA and UMAP representations, the *Bonsai* representations systematically improve at estimating the true distances between the points as the number of measurements increases (Fig S11). That is, by representing the similarities between the points on a tree, the estimates of the true positions of the points increases.

*Bonsai* thus automatically accomplishes the type of regularization that data-diffusion methods aim at by estimating states of individual cells by smoothing over the states of predicted nearest-neighbors, e.g. [38, 39].

### Visualization and exploratory analysis of large-scale datasets with *Bonsai-scout*

To facilitate the analysis of large datasets, the *Bonsai* -algorithm employs various mathematical and computational speed-ups so that its computation time and memory usage stay within acceptable limits (see Methods, and SI.C and SI.D). In Figure 3A we show that the computation time scales approximately as 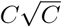 with the number of cells *C*, which is notable since a naive calculation of pairwise distances already scales quadratically. To demonstrate *Bonsai*’s efficiency and provide resources for the community, we reconstructed a *Bonsai* tree for the well-known *Tabula Muris* [4] dataset with almost 70k mouse cells (Fig 3B-D), and for more than 100k human epithelial cells from the *Tabula Sapiens* (Fig S12) [5].

**Figure 3.**
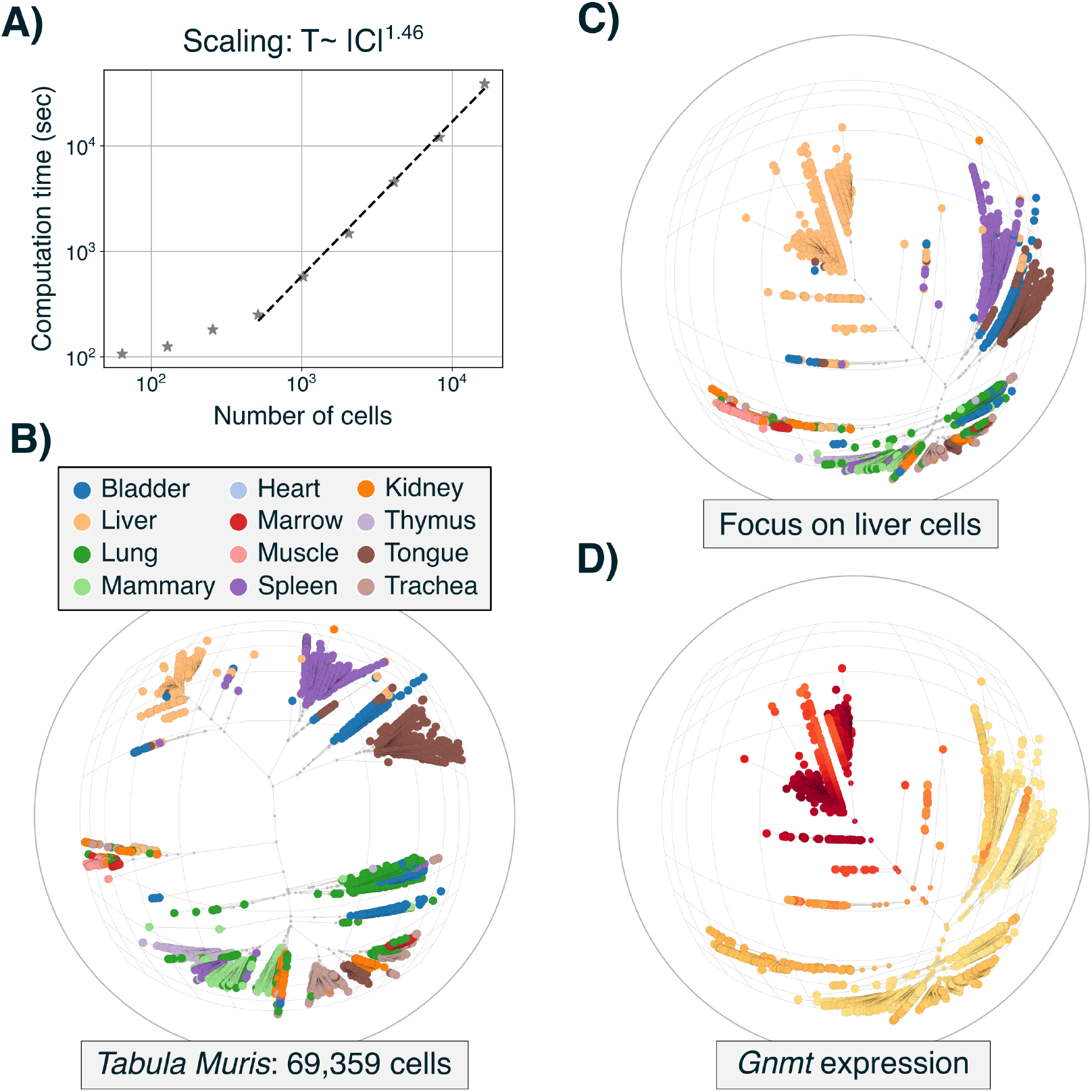
*Bonsai*’s computational efficiency allows analysis of cell atlases that can be explored with *Bonsai-scout*. **A)** *Bonsai*’s computation times using 20 CPUs as a function of the number of cells for simulated datasets with 2^6^ = 64 up to 2^14^ = 16 384 cells. Since *Bonsai*’s computation time scales approximately as 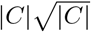, it is possible to reconstruct trees on large-scale atlases such as the Tabula Muris [4]. **B)** To facilitate the analysis of *Bonsai*’s tree representations, we developed *Bonsai-scout*, an interactive visualization tool that allows various exploratory analyses. The panel shows a visualization of the Tabula muris atlas with cells colored by tissue of origin. **C)** The visualization can be recentered to focus on any area of the tree, e.g. to focus on the clade of liver cells. **D)** *Bonsai-scout* allows finding marker genes that distinguish between different clades, and displaying the expression of these markers on the tree. Here the expression of liver marker gene Gnmt is shown.

We further had to overcome the challenge of visualizing trees with tens of thousands of cells. In conventional 2D-embeddings of large datasets, points are inevitably packed together so closely that any visual analysis is influenced not only by the parameters of the visualization method but even by the order in which points are plotted. Fortunately, the hierarchical structure of trees provides a natural way to navigate and zoom in on different subtrees. Although there are tools for DNA sequence analysis that can handle large trees, e.g. [40], these are not suitable for exploring single-cell data. We have therefore built *Bonsai-scout*, a customized application that facilitates exploring *Bonsai* trees, as we showcase in Figure 3B-D and Fig S12 for the Tabula Muris and Tabula Sapiens datasets.

*Bonsai-scout* supports different layouts of the tree, including the equal-daylight layout [19] shown in Fig 3 and the more conventional dendogram (Fig S12). When using the equal-daylight layout, the tree is shown on a hyperbolic disk with tunable curvature, where distances naturally get compressed toward the edges of the disk [41]. The user can interactively control the visualization by re-centering it so as to focus on one area of the tree (Fig 3C), actively zooming in to a particular part, flipping branches, overlaying cell type annotations (Fig 3B-C), and so on. *Bonsai-scout* also implements a number of methods for interactive exploratory analysis such as showing the expression of any gene on the tree (Fig 3D), cutting the tree into any number of major clusters (i.e. clades, see Methods), and even identifying marker genes for any clade of interest (Fig 3D and see Methods). We make *Bonsai-scout* available both as a standalone program that users can run themselves, and through a webserver where the *Bonsai* representations of the Tabula Muris and Tabula Sapiens atlases are already available for exploration (see Code and data availability).

Finally, to facilitate *Bonsai* analysis we have implemented an automated analysis pipeline at `bonsai.unibas.ch` where users can upload the UMI count table for their own dataset. First, *Cellstates* [29] is run to identify groups of cells with statistically indistinguishable gene expression states, and *Sanity* [28] is used to obtain normalized gene expression levels (i.e. log transcription quotients) and their error-bars. Finally, these results are used to reconstruct the *Bonsai* tree, after which the users get the flat file results as well as a web-address where the results can be explored using *Bonsai-scout*.

### *Bonsai* infers known and novel lineage relationships of blood cell types

To test *Bonsai*’s ability to recover known lineage relationships and uncover novel biology, we applied it to a scRNA-seq dataset of 7509 cord blood mononuclear cells [22]. This dataset is ideal for testing *Bonsai* because the extensive study of blood cell formation has resulted in considerable prior knowledge, both on the cell type differentiation hierarchy and on which genes can be used as cell type markers [42–44]. In addition, this dataset includes measurements of the single-cell abundance of 13 cell-surface proteins often used for cell type classification. We used only the scRNA-seq data to reconstruct the *Bonsai* -tree so that we could use the surface protein data as independent validation.

In Figure 4A *Bonsai*’s representation is visualized as a dendogram with cells colored based on the cell type annotations from the original publication [22]. It is broadly accepted that at the coarsest level, the lineages of blood cells divide in a lymphoid branch and a myeloid branch. Within the lymphoid branch, cells further differentiate in T-cells, B-cells, and NK-cells, whereas in the myeloid branch cells differentiate into monocytes, red blood cells, and megakaryocytes. We find that the lineage relationships inferred by *Bonsai* recapitulate the broad differentiation into lymphoid and myeloid branches and this is confirmed by the expression of the myeloid marker protein CD11c (Fig 4B). Furthermore, apart from these major branches, *Bonsai* also identifies distinct clades with known cell types, e.g. we see a distinct clade of T-cells, as confirmed by the expression of the CD3-marker (Fig 4B). We also see that red blood cells form a distinct clade that is very distant from all other cell types, which is in line with their unique transcriptional profile dominated by hemoglobin expression.

**Figure 4.**
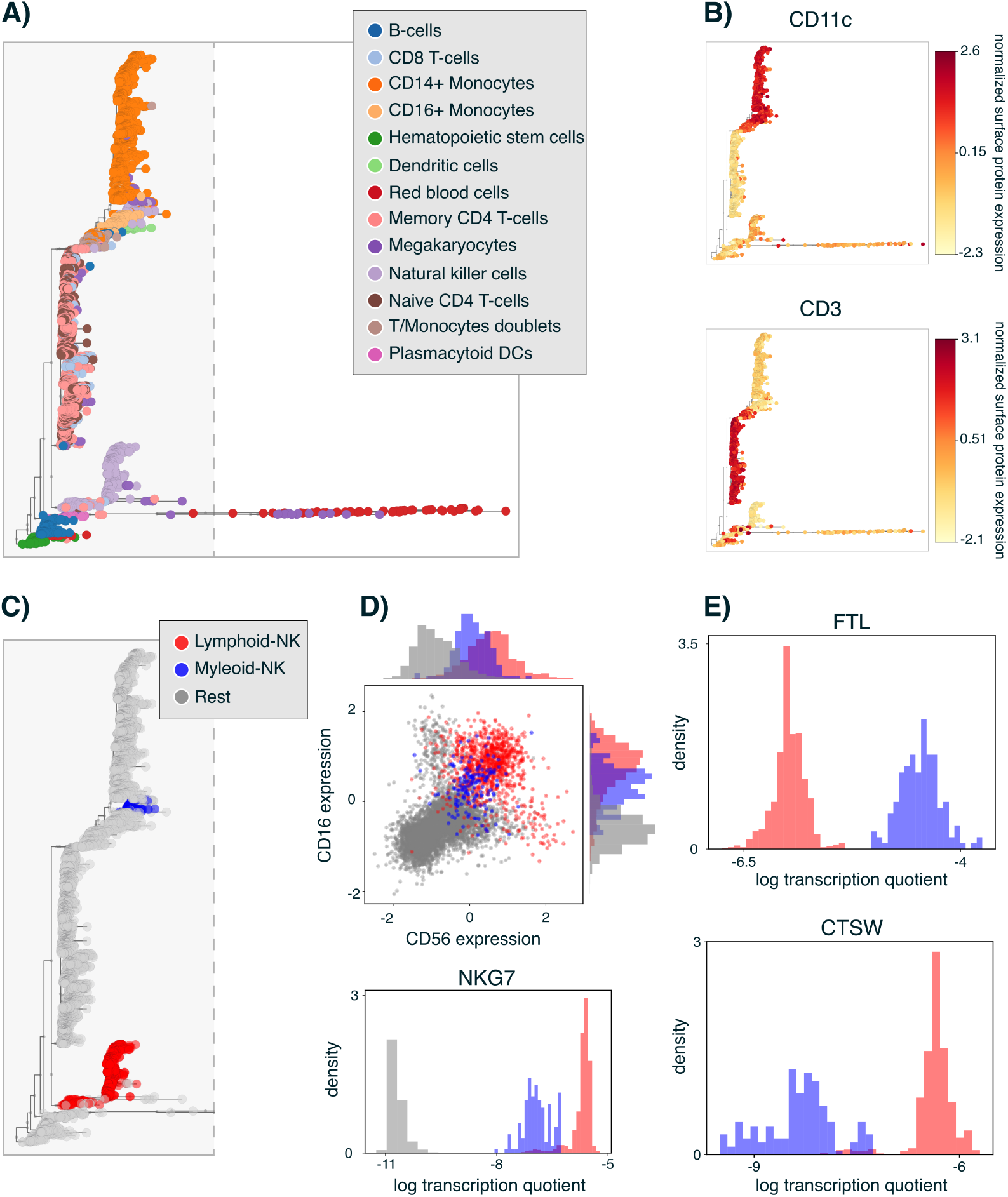
The *Bonsai* representation of a blood cell dataset recovers known lineage relationships and infers that different groups of NK-cells derive from different lineages. **A)** The *Bonsai* -tree representing a dataset of cord blood cells [22], where the leaf-nodes are colored by the celltype annotation provided with the original publication. Note that, in this dendrogram layout, the cell-to-cell distances are captured only by the horizontal distances along the tree (i.e. vertical distances are meaningless). **B)** Overlay of the expression of the surface proteins CD11c and CD3 which are markers for myeloid-cells and T-cells, respectively. **C)** Cells that were annotated as NK-cells are inferred to derive from different lineages, with one occurring within the myeloid branch (highlighted in blue) and one within the lymphoid branch (highlighted in red). **D)** In the top panel, the expression of the NK-cell surface markers CD56 and CD16 confirms the NK-cell annotation for both the lymphoid-NK cells (red) and myeloid-NK cells (blue) as compared to the remaining cells (grey). In the bottom panel, histograms of the expression levels (log transcription quotients) of the NK-marker gene NKG7 further confirm the NK-cell annotation. **E)** Histograms of the expression levels of two genes that were identified as markers of the myeloid-NK (FTL, top panel) and lymphoid-NK (CTSW, bottom panel) subpopulations.

Apart from recovering known lineage relationships among blood cell types, *Bonsai* also makes novel biological predictions. For example, we surprisingly find that cells annotated as natural killer cells (NK-cells) separate into one group located in the lymphoid-part of the tree and another group within the myeloid branch (Fig 4C). While it is commonly assumed that all NK-cells derive from lymphoid progenitors [45–49], *in vitro* studies have shown that NK-cells can be generated from myeloid progenitors [50, 51]. However, whether such myeloid-derived NK-cells also occur in the developmental trajectory of NK-cells *in vivo* is under debate [52].

We thus find that the *Bonsai* representation supports the hypothesis that NK-cells can originate from both lymphoid and myeloid progenitors. We find that both groups indeed express known markers for NK-cells (Fig 4D), but our marker gene analysis (see Methods) also identifies a substantial number of genes whose expression clearly separates the lymphoid-NK from myeloid-NK cells (Fig 4E and Supplementary Figs S13 and S14). Looking at these marker genes, we may even speculate about potential functional differences between myeloid-NK and lymphoid-NK cells. It appears that the genes that are upregulated in the myeloid lineage are more related to the recognition of ‘foreign’ molecular patterns, metal ion sequestration and release of granules with antimicrobial activity, while the genes that are up in the lymphoid lineage are more involved in cytolysis (cell killing) and the destruction of virus-infected and tumor cells. Notably, markers of the lymphoid-NK cells include MHC class I genes, whereas markers of the myeloid-NK cells include MHC class II genes. This is consistent with the proposed functional subdivision, since MHC class I molecules present intra-cellular antigens to CD8+ cytotoxic T cells which can kill infected or cancerous cells, whereas MHC class II molecules primarily present extracellular antigens to CD4+ helper T cells. While these are just speculations at this point, they underscore that *Bonsai*’s analysis immediately suggests specific novel biological hypotheses.

In summary, application of *Bonsai* to a dataset of blood cells not only confirms that it is able to recover the known lineage relationships of blood cell types, but also shows that *Bonsai* uncovers interesting novel biological findings.

### *Bonsai* is applicable to any set of objects with high-dimensional continuous features

We propose that tree structures are generally suited for representing relationships between highdimensional objects, so that *Bonsai* should be applicable to explore structure within data-types far beyond scRNA-seq. To test *Bonsai*’s general applicability, we applied it to a dataset of a completely different nature, but for which we still have some intuition for its structure: a dataset of football player statistics [53]. We preprocessed the data by shifting and normalizing each feature to have zero mean and unit variance, and then projected the data on the first 50 PCA components.

Figure 5A, shows that, despite this primitive pre-processing, *Bonsai*’s tree structure identifies 12 well-separated and interpretable groups of players. Indeed, by visualizing marker features identified by *Bonsai-scout* (see Fig 5C and Suppl. Figs. S15-S17), we show that these groups correspond to players that fulfill particular roles in their teams. Additionally, *Bonsai* identifies some players as clear outliers, indicating that they show exceptional statistics, and, remarkably, we find that these were indeed some of the highest-reputed football players in the season of 2021-2022. In summary, although we are not claiming that *Bonsai*’s analysis identifies new structure in football player statistics that is not already known by experts, the results demonstrate that *Bonsai* automatically identifies subtle and meaningful structures across a wide variety of data types.

**Figure 5.**
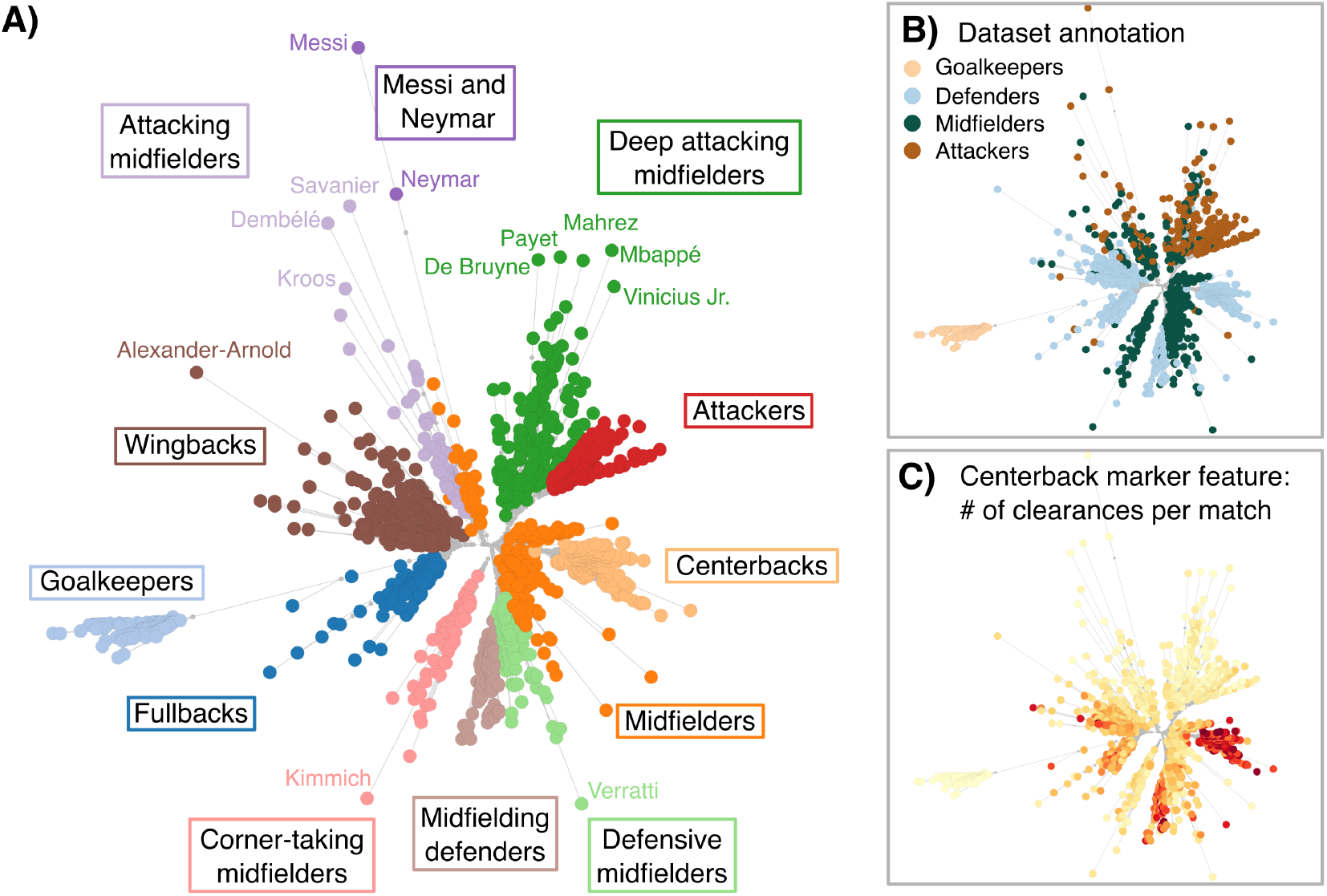
*Bonsai* is applicable to a wide variety of datasets. **A)** The *Bonsai* -tree representing the statistics of 1436 football players from five international competitions in the season 2021-2022 [53]. The colors indicate 12 clusters of players that were found by our tree-based clustering method (see Methods), which we manually annotated using example players and marker features (see Supp. Figs. S15-S17). In addition, names of the 13 most outlying players are shown. The same *Bonsai* tree with players colored by their annotated position on the field shows that these clearly cluster on the tree. **C)** An example of a marker feature identified by *Bonsai-scout* : central defenders can be distinguished from the other players based on the number of clearances per match.

## Discussion

The rapidly increasing number of large, high-dimensional single-cell omics datasets has the potential to provide transformative new insights into fundamental biological questions, such as what drives cell differentiation, what characterizes cell identity, and how ‘cell types’ might be rigorously defined. However, to fulfill this potential, since we do not *a priori* know what complex high-dimensional structures may be hidden within these large noisy datasets, there is an urgent need for exploratory analysis and visualization tools that do not compromise on accuracy. In particular, in order to not have to rely on prior biological knowledge, it is crucial that the structures that such visualization methods display can be trusted not to distort the true structure in the high-dimensional data, so that their predictions can be relied upon. Unfortunately, for lack of a better alternative, most researchers in the field continue to turn to visualization tools [9, 10] that are well known to severely distort the true structure in the data and hallucinate structure that does not exist [11–13].

Here we have shown that by representing scRNA-seq datasets on a tree with the cells at its leaves, it is possible to accurately capture all structure in these high-dimensional data, e.g. such that all pairwise distances between cells closely match the distances along the branches of the tree (Fig. 2). Moreover, by applying our *Bonsai* method to scRNA-seq data with known lineage relationships, we not only showed that *Bonsai* recovers these lineage relationships, i.e. the lineage structure of major blood cell types, but also provides intriguing novel predictions, i.e that natural killer cells divide into lymphoid- and myeloid-derived groups with clearly distinct expression profiles that reflect their origin (Fig. 4).

Representing structures in high-dimensional data by a hierarchical representation is of course not novel in itself. Indeed, hierarchical clustering of data has been a popular approach from the earliest days of transcriptomics [54] and this raises the question as to why such approaches have so little popularity in the analysis of single-cell omics data. The answer is that in order to meaningfully capture the relationships between large numbers of cells from these highly noisy and sparse data, a substantial number of difficult technical challenges needs to be overcome. First, we needed to develop methods that appropriately take into account both the biological and measurement noise in the data because scRNA-seq data is not only very noisy and sparse, the noise is also highly heterogeneous, with noise levels varying over several orders of magnitude across genes and cells. Second, we had to develop a rigorous probabilistic model for the movement of cells through gene expression space. Third, given the large numbers of cells at the leaves of the tree representation, the space of possible tree structures is enormous and we needed to develop methods that allow effective search of this space. For example, we showed that the high-dimensional positions of all nodes of the tree can be analytically integrated out of the likelihood function, so that the likelihood depends only on the topology and branch lengths of the tree. Further, we developed efficient numerical methods not only to optimize branch lengths given a tree topology, but also adapted methods from phylogenetics to effectively search the space of tree topologies, allowing us to reconstruct trees for datasets with over a hundred thousand cells.

Because we did not want to make any strong prior assumptions about the structure of the data or the movement of cells through gene expression space, our likelihood model assumed only that, on short time scales, gene expression changes are continuous, Markovian, and without any prior bias in direction. We showed that this leads to prior probabilities matching homogeneous diffusion through the high-dimensional gene expression space. Although this implies that *a priori* there are no expected correlations in the expression changes of different genes, if such correlations are evident in the *data*, then *Bonsai* will readily infer such expression correlations (e.g., Suppl. Fig S19).

Apart from providing reliable visualizations, one of the key advantages of *Bonsai* is that, by inferring the gene expression states at all internal nodes of the tree, it allows us to quantify the expression changes that occurred along the branches of the tree. Indeed, it is well accepted within the field of evolutionary theory that in order to understand evolutionary dynamics, one should focus on the changes that occur along the branches of the phylogeny, e.g. [55]. Similarly, in order to understand the gene regulatory circuitry underlying cellular differentiation and cell identity, it is far more informative to study the correlation structure of gene expression changes along the branches of the tree, than comparing gene expression correlations directly across the cells, because the movements along the branches reflect the differentiation trajectories of the cells. Although many methods have been proposed for trajectory inference in scRNA-seq data, these are typically based on either a nearest-neighbor graph or on a low-dimensional embedding of the data [56–58], but both of these pre-processing steps are heavily affected by noise and have proven to be shaky foundations for further analysis [11–13, 59, 60]. In contrast, *Bonsai* reconstructs the tree that connects all cells in a way that maximizes the probability of the inferred gene expression changes along the branches of the tree, while rigorously taking into account the heterogeneous measurement noise properties of the cells at the leaves. Indeed, we showed that even on the basic task of nearest-neighbor identification, *Bonsai* strongly outperforms other methods (Fig. S10).

*Bonsai*’s tree representations of scRNA-seq data enable many different downstream analyses that would otherwise be very challenging to implement. For example, by measuring the correlation structure of gene expression changes along the branches of the tree, one can ask to what extent this correlation structure differs in different parts of the tree, which would help understand the role of regulatory ‘context’ for driving expression changes. As another example, every branch of the tree corresponds to a partition of the cells into two subsets, and we can systematically study how the set of markers of such bi-partitions varies along the tree, e.g. whether there are branches that are particularly enriched for clear marker genes, which would indicate the occurrence of major differentiation events. The structure of clades near the leaves can also be used to systematically estimate densities of cells in gene expression space. All these examples of possible downstream analyses illustrate how *Bonsai*’s tree representations can be used to help unravel the underlying gene regulatory circuitry.

There are many avenues for future extensions of the *Bonsai* analysis. First, although *Bonsai* was computationally optimized to run on datasets of more than a hundred thousand cells, and the absence of tunable parameters means *Bonsai* only needs to be run once on a dataset, it would still take a long time to process the largest single-cell datasets that contain tens of millions of cells. One approach to analyzing such larger datasets would be to use prior information to provide *Bonsai* groups of cells that are known to form clades on the tree. For example, for scRNA-seq we recommend using *cellstates* to group cells that are statistically indistinguishable, since *cellstates* also correctly models the Poisson noise in the data [29]. For very large datasets, *Bonsai* can probably be sped up by using heuristics that have recently scaled phylogenetic reconstructions to millions of genomes [37, 61, 62]. For example, we could first reconstruct a backbone of the tree based on a subset of the data, after which the remaining cells can be iteratively placed on the backbone. For now, however, we focused on contributing a robust method that reconstructs the most likely single-cell tree on datasets of most sizes, rather than trading off rigor for scalability.

We foresee that *Bonsai* can be of great use in the analysis of multimodal datasets. For example, it is currently possible to obtain both transcriptomic (scRNA-seq) and epigenomic (scATAC-seq) read-outs from the same cells [63], but the scATAC-seq data is even more sparse and noisy than scRNA-seq. Often, scATAC-seq analysis proceeds by first grouping cells into so-called pseudo-bulk samples, after which downstream analyses can be performed that would be insufficiently robust at the single-cell level. *Bonsai* provides a principled way of doing this because the pseudo-bulk samples can be defined at different scales by the clades of the tree reconstructed on the scRNA-seq modality. Additionally, since after suitable pre-processing, we can also build a tree on the scATAC-seq data, this provides the opportunity to study the differences between the transcriptomic and epigenomic cell-to-cell relations by comparing the *Bonsai* -trees of both modalities. Finally, we could combine the information of all modalities by using the combined data to reconstruct a single tree representation.

It is increasingly appreciated that having lineage information at the single-cell level is crucial for understanding gene expression dynamics [64, 65] and there are now various experimental techniques that simultaneously measure single-cell transcriptomes while recording lineage histories by creating random mutations on genetically encoded barcodes [66–71]. However, it has become clear that the lineages cannot be fully reconstructed based on the lineage tracing information alone [72]. Two methods have been proposed to combine the scRNA-seq and lineage tracing data for lineage reconstruction [73, 74], but these methods are unsatisfactory for a number of reasons, i.e. the two modalities are used sequentially rather than on an equal footing, the transcriptomic information is discretized into preconceived cell types, and one of the methods does not scale to large datasets. A very exciting prospect is to extend *Bonsai* to combine the gene expression information with lineage tracing data by extending the likelihood function to include the observed mutations in the lineage-tracing barcodes. We propose that this would not only allow more accurate lineage reconstructions, it would also allow systematic study of the extent to which gene expression and cell division histories align.

Finally, our successful application of *Bonsai* to a non-scientific dataset of a fundamentally different nature, i.e. statistics on professional football players (Fig 5), suggests that its tree representations are able to accurately capture structure in high-dimensional data of almost any type. Intriguingly, as high-throughput approaches have become common across different scientific fields, there are many areas where researchers have turned to t-SNE and UMAP for visualizing structure in their datasets. This includes data from cytometry [23], metagenomics [24], population genetics [25], neuronal spiking behavior [26], and even image and text categorization [27]. Notably, like scRNA-seq data that we focused on here, many of these other types of data likely suffer from the same heterogeneous measurement noise properties that *Bonsai* is uniquely equipped to deal with. We thus expect that for data from all these areas, *Bonsai*’s tree representations will be able to much more accurately capture the structure in these data, thereby providing a powerful method of exploratory data analysis across a wide variety of scientific fields.

## Methods

In the following sections we will give an overview of the *Bonsai* method, its implementation, and how specific datasets were analyzed. More detailed mathematical derivations underlying *Bonsai* and details of its computational implementation are provided in the Supporting Information.

### Recursive calculation of the tree likelihood

The foundation of *Bonsai* is a likelihood model for the observed data, **cell**_*i*_, of each cell *i* given a tree topology *T* and a set of branch lengths ***t***, which takes the form

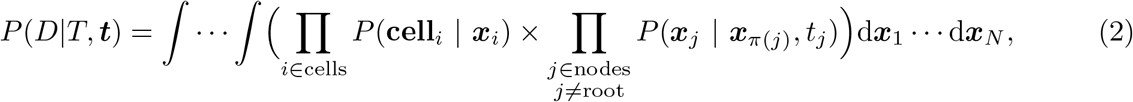

where the integrals are over the true positions ***x***_*j*_ of each node *j* in gene expression space. As described in SI.B, a crucial ingredient to the tractability of this model is that all factors in the likelihood have a Gaussian form which also factorizes over the genes. First, the likelihood of the observed data **cell**_*i*_ given the associated node’s true position ***x***_*i*_ is given by:

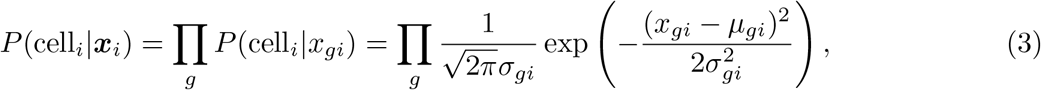

where *µ*_*gi*_ is the estimated cell-position from the data, and *σ*_*gi*_ is the standard-deviation on this estimate for each gene *g*. Second, the likelihood of the gene expression change along the tree-branch leading from node *π*(*i*) to *i* is given by:

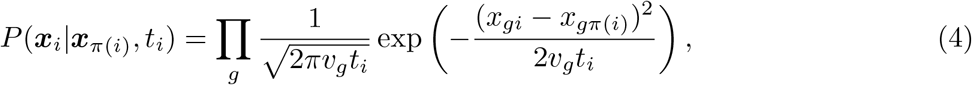

where *x*_*gi*_ and *x*_*gπ*(*i*)_ give the coordinates of the two nodes, *v*_*g*_ is the estimated variance in the expression of gene *g*, and *t*_*i*_ is the length of the branch.

As described in SI.B.2, we found that, for any tree-topology, this likelihood can be efficiently calculated recursively, giving a continuous version of Felsenstein’s pruning algorithm [75]. For each node, say *j*, we can express the likelihood of all the data downstream of this node in the tree as a Gaussian function with effective positions 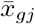 and associated standard-deviations 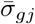 for each gene *g*. We can thus replace the likelihood contribution of the entire subtree downstream of node *j* by the likelihood of an *effective leaf*. This means that we can recursively simplify the tree by *pruning* off sub-trees and summarizing their likelihoods as effective leaf contributions. Eventually, we will get the likelihood of all data as a function of the root’s position *P* (*D*|*T*, ***x***_root_), where we have thus marginalized over all other node positions. Importantly, because we assume a uniform prior on the root’s position, this expression is also directly proportional to the posterior distribution on the root’s position *P* (***x***_root_ |*D, T*), which we will exploit below. To get the likelihood of the data conditioned only on the tree *T*, however, we additionally marginalize over the root’s position. Importantly, thanks to this pruning procedure, all integrals in equation (2) can be performed analytically, allowing the likelihood *P* (*D*|*T*, ***t***) to be calculated efficiently.

### Intuitive approximation of the likelihood function

It is helpful to develop an intuition regarding how the likelihood of a tree topology depends on the distances along the branches of the tree. To do this, we calculate a log-likelihood *L*(*T*) of a tree topology *T* by considering the simplified situation where there is no measurement noise, i.e. the position ***x***_*i*_ of each cell *i* at the leaves is known precisely. In addition, instead of marginalizing over all the positions of the internal nodes and optimizing the branch lengths, we optimize both with respect to all internal node positions and all branch lengths.

We then find that the log-likelihood of a tree topology *T* is simply given (up to an additive constant) by a sum over the logarithms of the squared-distances along the branches of the tree, i.e.

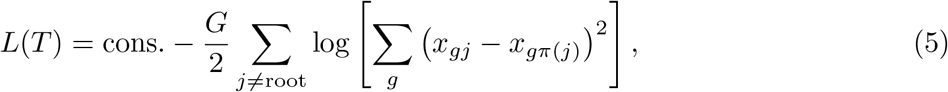

where *G* is the number of genes, *π*(*j*) is the ancestor node of node *j*, and the positions ***x***_*j*_ at the internal nodes are set so as to maximize this likelihood. This occurs when the position ***x***_*j*_ of each internal node *j* equals the weighted average of the nodes that it is connected to, with weights given by the inverse squared distances to each of the nodes. That is, the optimal internal node positions ***x***_*j*_ obey

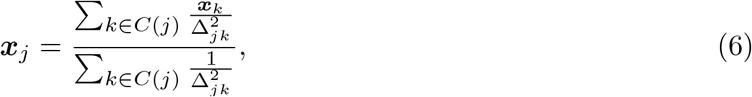

with the squared-distances along the branches

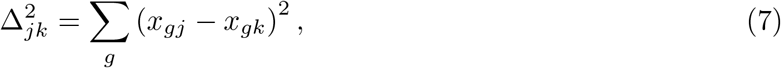

and *C*(*j*) the set of nodes that are connected to node *j*.

Although this simple expression for the likelihood of a topology only holds when there is no measurement noise on the leaves and when we optimize not only the branch lengths but also the positions of the internal nodes (rather than marginalizing over them), it does illustrate that under our model the log-likelihood does not correspond to a sum of distances or squared-distances along the branches, but rather approximately equals the sum of the *logarithms* of the distances along the branches.

### Obtaining posteriors over the positions of the internal nodes

In the Supplementary Information, we show that the tree likelihood is independent of which node we choose as the root of the tree. Therefore, by picking a node of interest *i* as the root, and marginalizing over all other nodes’ positions as outlined above, we can get a posterior distribution *P* (***x***_*i*_|*D, T*, ***t***) for the position ***x***_*i*_ of each node *i*.

This posterior gives the probability of the internal node being in a certain gene expression state, conditioned on the data and the tree topology with corresponding branch lengths. These posteriors all have Gaussian form, and *Bonsai* reports their means and standard deviations which can then be used for downstream analysis, such as in the *Bonsai* -scout tool for data visualization and marker gene detection.

### An outline of Bonsai’s tree-search algorithm

We here shortly outline how we search the large space of possible trees for the tree topology and branch lengths that jointly maximize the likelihood *P* (*D*|*T*, ***t***) of the data; we refer to SI.C for details. The main steps in *Bonsai*’s tree-search are:

1. Start from a star-tree with optimized branch lengths.
2. Iteratively add internal nodes to maximally increase the likelihood.
3. Resolve polytomies.
4. Re-optimize the branch lengths.
5. Further increase the likelihood by interchanging nearest neighbors.
6. Perform a final re-optimization of the branch lengths.

To start, we create a tree where each data-point has an associated leaf-node, and these leaf-nodes are all connected to a root-node, i.e. a star-tree. For this star-tree, we can efficiently optimize the branch lengths, after which we iteratively add internal nodes to maximally increase the likelihood at each step.

Importantly, it is also possible to start with a different initial tree, as described in the Discussion section. For example, we recommend for scRNA-seq data to first calculate *cellstates* [29] to group cells that are statistically indistinguishable. One can then start from a tree in which, for each *cellstate*, the cells in this *cellstate* are connected to an ancestor node, and each of these ancestors is in turn connected to the root node, i.e. forcing the cells within each *cellstate* to form a clade in the tree.

From the initial tree, we iteratively pick a pair of leaves and add an internal node upstream of these two leaves so as to maximally increase the likelihood of the tree. To do this, we calculate, for each candidate node-pair, how much the likelihood would increase if we would add an internal node upstream of that pair with optimal branch lengths from the internal node to the two leaves and to the root. After adding the internal node that increases the likelihood the most, we use the pruning procedure described above: we summarize all the likelihood-information of this node-pair by replacing the newly added internal node by an effective leaf with a corresponding effective position and effective uncertainty (see S20 for an explanatory illustration). As such, we can again describe the tree as a star-tree, where the two leaves and their newly added ancestor have been replaced by an ‘effective leaf’. We iterate this procedure until we either have a fully resolved tree, or we can no longer add any internal node directly downstream of the root that increases the likelihood.

There are two reasons that this procedure can lead to polytomies (nodes with more than three connecting branches). First, in optimizing the branch lengths we can get zero branch lengths. For example, let node *i* be the ancestor of two leaves *l*_1_, *l*_2_, and say that we now add an ancestor *a* upstream of this node *i* and another leaf *l*_3_. It may be that the optimal branch length from *a* to *i* is zero, such that, effectively, node *a* and *i* are merged. As a result, the resulting tree would have one ancestor connected to the three leaves *l*_1_, *l*_2_, and *l*_3_. Second, we could have picked an initial tree that already contains polytomies (such as the suggested one using *cellstates*). In both cases, we try to increase the likelihood further by resolving such polytomies. We do this by effectively treating the internal node with the polytomy as a new root, and then going over the above iterative procedure of adding internal nodes that increase the likelihood most.

We have found that the above greedy procedure generally leads to good approximations of the maximum likelihood tree. However, in most cases, local rearrangements to the tree can still increase the likelihood. We search for such local rearrangements by performing so-called Nearest Neighbor Interchanges (NNI). Every NNI-move starts by picking an edge, say from node *a* to *b*, and listing all the nearest-neighbor nodes, i.e., nodes that are either connected to *a* or *b*. An NNI-move is nothing more than changing the way in which this set of nearest neighbors is connected in the tree. In the simple case where we have no polytomies, this always comes down to interchanging a neighbor from *a* with one of *b. Bonsai* first performs a random NNI-phase, followed by a greedy NNI-phase. In the random phase, different NNI-moves are sampled weighted by their corresponding change in likelihood, while in the greedy phase, we deterministically perform the NNI-move that most increases the likelihood.

### Optional parameters

Bonsai does not require the user to specify any parameters but, if desired, there are a few optional variants of its default behavior that the user can specify. First, for typical scRNA-seq datasets there are many genes that are sampled so sparsely that the error-bars on their expression measurements are larger than the true variation in their expression values, and we have found that including these genes in Bonsai’s analysis is more likely to decrease than increase the performance. Thus, Bonsai calculates an average signal-to-noise ratio *S*_*g*_ for each gene as

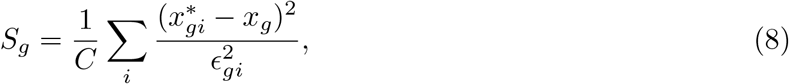

where *C* is the number of cells, 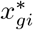 is Sanity’s estimate of the LTQ of gene *g* in cell *i, x*_*g*_ is the estimated average LTQ of gene *g*, and *ϵ*_*gi*_ the standard-deviation of the posterior of the LTQ as given by Sanity. For data other than scRNA-seq data processed by *Sanity*, Bonsai estimates the signal-to-noise of each feature from the input data as described in section SI.B.1.3.

By default only genes with *S*_*g*_ ≥ 1 are included in the analysis but the user can change this threshold value.

Finally, the signal-to-noise estimation also involves estimating the variance *v*_*g*_ of the signal and by default *Bonsai* uses a prior where the amount of diffusion in dimension *g* is assumed proportional to this variance *v*_*g*_ (see section SI.B.1.4). However, if desired the user can specify to not scale the prior by the variances *v*_*g*_.

### Finding clusters in the tree

*Bonsai-scout*, the interactive app that can be used for exploratory analysis and visualization of Bonsai trees, offers a method for finding clusters of cells in the tree. In particular, this method greedily cuts branches to create a set of subtrees which minimize the sum of pairwise distances between all leaves in the created subtrees.

More specifically, for any tree we can calculate the sum of pairwise distances between all leaves along the branches of the tree. Note that this sum differs from the sum of all branch lengths, because branches in the middle of the tree are traversed by many more leaf-to-leaf paths than branches near the leaves. Starting from the original tree, we create two subtrees by cutting the branch that minimizes the combined sum of pairwise distances between leaves in the two resulting subtrees. Given these new trees, we can now cut another branch to again minimize the pairwise distances in the three resulting subtrees, and so on. To create *n* clusters, one repeats this cutting procedure *n* − 1 times.

### Choosing a root node

The final tree that *Bonsai* reports (in Newick format) is rooted. By default, we choose to position the root on the branch that would first be cut in the clustering procedure we described above.

Since the likelihood of the data is independent of the choice of a root-node, this choice is somewhat arbitrary. Notably, the choice of root affects none of the results, but it of course influences the visualization of the tree: when using our Bonsai-scout visualization tool, the root determines the initial centerpoint for circular layouts, and it determines the leftmost point in a dendrogram layout. However, for the circular layout users can double-click to focus the visualization on any arbitrary point in the tree, and for the dendrogram layout the root can be re-positioned interactively.

### The detection of marker genes

*Bonsai* can additionally be used for the detection of marker genes/features that best distinguish the cells/objects from two clades of the tree. This was for example used to highlight the differences between the two groups of NK cells in the blood cell dataset (see S13-S14), and to annotate the different football player clusters (see S15-S18). Let *C*_1_ and *C*_2_ denote two clades of cells on the tree. We define marker genes as genes that either maximize or minimize the probability that, when picking a random pair of cells, *c*_1_ ∈ *C*_1_ and *c*_2_ ∈ *C*_2_, the gene expression 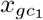 is higher than 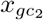. Dropping the gene-indices, we thus want to calculate for each gene:

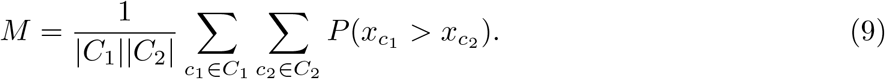

To calculate this, we use the posteriors over the positions of the cell-associated leaf-nodes that *Bonsai* provides. Specifically, for cell *c*_1_, the posterior 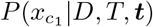 is a Gaussian distribution with mean 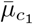 and variance 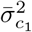, and similarly for cell *c*_2_. To obtain the probability 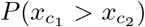, we note that the distribution of the difference, 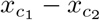, will again be Gaussian with mean 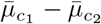 and variance 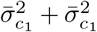. Therefore, we can express the desired probability in terms of the cumulative distribution of this Gaussian, which can be efficiently calculated using the error-function. This gives

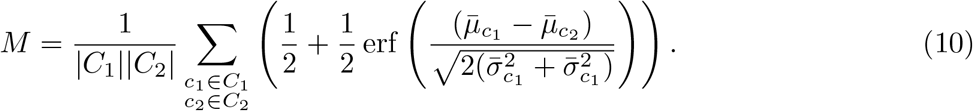

Marker genes are those for which *M* is close to 1 or 0. A value close to 1 means that a high value for this gene is indicative of the corresponding cell being in group *C*_1_, and vice versa for *M* close to 0. We provide a supplementary tool with the Bonsai code that takes a file that specifies the identifiers of the cells in the two groups and calculates the marker scores *M* for all genes. This supplementary tool can be found in the Bonsai code repository under `downstream_analyses/calc_marker_genes.py`.

Although the above is the most accurate way of calculating marker scores, the calculation involves a sum over all pairs of cells from the two clades, which for large clades can be too slow to calculate ‘on the fly’ within the interactive Bonsai-scout. Therefore, to facilitate detecting marker genes in an exploratory data analysis within Bonsai-scout, when the number of cells in the two clades is large, we approximate the marker gene calculation by ignoring the uncertainties on the cell posteriors. In that case, the marker score reduces to

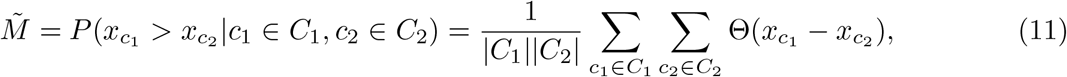

where Θ(•) is the Heaviside-function that is 1 for values larger than zero, and zero otherwise. Now let 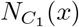 be the number of cells in *C*_1_ that have an expression value lower than *x*, 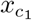 the expression level of cell *c*_1_, and rk(*c*_1_) the rank of cell *c*_1_ in the list of all cells from *C*_1_ and *C*_2_ sorted in increasing order of expression. We then get

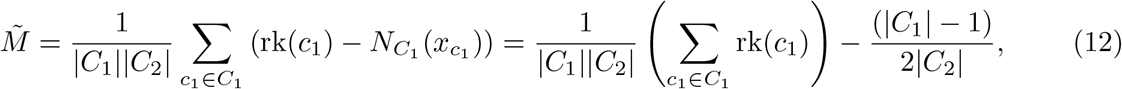

where we used in the second step that we can reorder the sum over *c*_1_ such that we get a simple arithmetic sequence. We recommend using this approximation of the marker score to interactively analyze the data, after which the expression from Equation (10) can be used to confirm or falsify specific hypotheses. Importantly, once we resort to this approximation of the marker score in *Bonsai-scout*, the user gets notified. *Bonsai-scout* then allows for downloading the file that can directly be used as input for the Python-script that we provide to obtain the full marker scores, and which can be found at (`downstream_analyses/calc_marker_genes.py`).

### Detailed descriptions of the shown *Bonsai* reconstructions

#### Processing of simulated datasets

We first process the raw UMI counts of the simulated datasets using *Sanity* [28] and then run *Bonsai* with the default signal-to-noise threshold of 1.0.

UMAP and PCA-visualizations follow the most commonly used preprocessing steps. The rawcounts are log-transformed with a pseudocount: 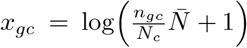, where *N*_*c*_ is the total UMI-count for cell *c*, and 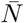 is the average total UMI-counts over all cells. These log-transformed data are projected on the first 2 PCA-components for creating the PCA-visualization, and to the first 10 PCA-components as preprocessing for the UMAP-embedding. Finally, we run UMAP with the parameters *n neighbors=15, min dist=0*.*1, n components=2, metric=‘euclidean’*.

#### Processing of the cord-blood dataset [22]

We downloaded the FASTQ files stored under the GEO accession GSE100866, after which we pseudoaligned the reads with `Kallisto` using the parameter -x “0,0,16:0,16,25:1,0,0”. With `bustools`, we collapsed the UMIs and error-corrected the barcodes using a whitelist. Using the `R`-package `emptyDrops`, we removed empty cells and only retained cells with total UMI count of at least 500. From the remaining cells, we decided to keep only the cells that were also selected in the original publication such that celltype-annotation was available. Finally, for each promoter, we counted the number of observed transcripts (i.e. UMIs) that were initiated from it. After obtaining the UMI count table (with promoters as rows and cells as columns), we ran *Sanity* [28] and *cellstates* [29] on these raw counts.

Bonsai was then run on the log-fold changes inferred by Sanity (stored in `delta_vmax.txt`) and the corresponding standard deviations of their posteriors (stored in `d_delta_vmax.txt`). We start *Bonsai* from an initial tree that is informed by the *cellstates*-clusters, i.e. we connect all cells from each *cellstates*-cluster to a single ancestor, which is in turn connected to the root. In the initial tree, we thus have as many ancestors as the number of cellstates.

For visualizing the surface protein expression, we used the normalized values as described and provided in the original publication [22].

#### Processing of *Tabula Muris*

Raw FASTQ-files were downloaded from GEO accession GSE109774. Further processing was done in the same way as described above for the cord blood cell data.

#### Processing of *Tabula Sapiens*

For the Tabula Sapiens dataset, we downloaded the matrix of UMI-counts from the CZI-database (https://cellxgene.cziscience.com/collections/e5f58829-1a66-40b5-a624-9046778e74f5). After that, we ran *cellstates* [29] for two weeks. Although at that point, the *cellstates*-algorithm could not guarantee that there was not an even more likely clustering to be found, it already reduced the dataset of more than 104 thousand cells to 4000 clusters of cells that were determined to be statistically indistinguishable. Even though *cellstates* could probably have grouped even more cells or clusters together, this would have taken more time. Therefore, we have used the clusters that *cellstates* found up to that point to inform the initial tree for *Bonsai*. Notably, it is unlikely that the final *Bonsai* -results are strongly affected by using cellstates from this incomplete run of *cellstates*, since clusters that would eventually have been merged by *cellstates* are likely to end up close together in the *Bonsai* -tree in any case.

#### Processing of the football statistics dataset

We downloaded the dataset with football player statistics from https://www.kaggle.com/datasets/vivovinco/20212022-football-player-stats. To preprocess this data, we first dropped players with unreliable statistics by: 1) removing 1 player with NaN-values, 2) keeping only the 50% players that played the most minutes to avoid outliers, and 3) removing 12 players that switched teams halfway through the season (unfortunately also dropping Christiano Ronaldo). Then, since the measured features are on vastly different scales, we normalized this by first subtracting the mean for every feature and then normaling the variance of each feature to 1. After this, we projected the data on the first 50 PCA components. In this way, we reduce the effect that (partially) redundant statistics have on the results. The number of principal components used did not strongly affect the results. Then, because *Bonsai* expects uncertainty estimates on the input data but this information is not available for this dataset, we used an artificial small standard deviation of 10^−3^ for all features and players. The exact size of this standard deviation does not affect the *Bonsai* -results either.

For the analysis of the *Bonsai* -tree, we created clusters by iteratively cutting the branch that maximized the score |*L*_*us*_|| *L*_*ds*_ |*t*, where |*L*_*us*_|, |*L*_*ds*_| are the numbers of leaves upstream and downstream of the branch and *t* is the branch length. We found that 12 clusters still gave interpretable groups, although we do not exclude the possibility that there is useful information in a more finegrained clustering. Outlier players were detected by first ‘out-group rooting’ the tree, using the clade of goalkeepers. The intuition behind this is that the statistics for goalkeepers are so different from all other players, that the goalkeeper-clade must attach close to the ‘root of all field players’.

From this root, we defined the 13 outliers by checking which players were furthest removed from the root. These outliers are highlighted in Figure 5.

## Code and data availability

All code needed to run *Bonsai* and *Bonsai-scout* can be obtained from https://github.com/dhdegroot/Bonsai-data-representation. This repository contains an extensive read-me that explains how to install and run *Bonsai*. It also contains the scripts to generate the various simulated datasets, as well as Jupyter notebooks that can be used to reproduce all figures presented here.

The data used for the cord blood cell analysis, the *Tabula Muris*- and *Tabula Sapiens*-visualizations, and the football statistics analysis are all publicly available in the corresponding publications, but we have also made the data and the *Bonsai* results available at http://doi.org/10.5281/zenodo.15350325.

In addition, we already provide 5 example datasets that can be explored interactively in *Bonsai-scout* using the following webpages:

- The cord blood cells: https://bonsai.unibas.ch/bonsai-scout/?dir=Cord_blood_cells_CITE-seq.
- All epithelial cells from the *Tabula Sapiens*: https://bonsai.unibas.ch/bonsai-scout/?dir=Epithelial_cells_from_Tabula_Sapiens.
- The *Tabula Muris*: https://bonsai.unibas.ch/bonsai-scout/?dir=Tabula_Muris,
- The football player statistics: https://bonsai.unibas.ch/bonsai-scout/?dir=Football_statistics
- The simulated binary tree: https://bonsai.unibas.ch/bonsai-scout/?dir=Simulated_dataset_perfect_binary_tree,
- The simulated dataset presented in Figure 2: https://bonsai.unibas.ch/bonsai-scout/?dir=Simulated_dataset_unbalanced_tree_with_random_branch_lengths_and_realist ic_covariance,

### Obtaining a *Bonsai* -representation of scRNA-seq data using our automated pipeline

For scRNA-seq data, we have implemented an automated pipeline for scRNA-seq processing as a webserver at http://bonsai.unibas.ch. Here, users can upload a matrix with the raw mRNA counts per gene (rows) and cell (columns). In addition, users can upload annotation that they may already have for the cells, and that can be visualized on the tree. All analysis will then be performed automatically, which includes running *cellstates, Sanity*, and *Bonsai*. After all computations have finished, users will get an email with a link to the *Bonsai-scout* visualization of their data, and a link to a download page that contains the results of the three methods.

### Input and output of the *Bonsai* code

When using the *Bonsai* code independent of the automated pipeline, users should take note that *Bonsai* requires that the data is preprocessed such that the likelihood of the measurements is reasonably approximated by a multi-variate Gaussian with means *µ*_*gc*_, standard-deviations *σ*_*gc*_ and negligible covariances (as discussed in SI.B.1). *Bonsai* requires these means and standard-deviations as simple feature-by-object matrices. For scRNA-seq data, the data can best be preprocessed using *Sanity* [28], and in that case *Bonsai* should just be pointed towards the full *Sanity* output folder.

The output that *Bonsai* provides will contain:

- A Newick string that describes the tree topology and its branch lengths.
- Two files that describe the tree in a more human-readable format.
- Two *Numpy* binary files that describe the means and variances of the posteriors that *Bonsai* infers for the position of each node, i.e., for all cells at the leaves *and* for the internal nodes.
- A `.json`-file with metadata on the dataset, for example containing the cell-IDs, gene-IDs, and the inferred gene variances, but also paths to where the original data was read from.

This output from *Bonsai* can be used as input for *Bonsai-scout*, for which we first run a preprocessing script that produces a `.hdf`-file containing all necessary data and a `.json`-file containing the initial settings for the tree visualization. Since this `.json`-file is human-readable and -editable, it is possible to change these initial settings by hand, for example for customizing a colormap. These two files are the only necessary files for running *Bonsai-scout*.

## Acknowledgements

This work was supported by SNF grants 310030 184937 and CRSII5 189910 to EvN. We thank Björn Kscheschinski for feedback and inspiration during weekly group meetings on the *Bonsai* project. Anurag Ranjak has aided in the development of several downstream analysis methods, such as the tree-based clustering and the marker gene detection. Georg Angehrn has both been involved in the early conceptualization of the tree-based method, and in rigorously checking the derivations in the Supplementary Information.

## Contributions

## Supplementary Information

## Table of Contents of the Supplementary Information

## SI.A Supplementary Figures

**Figure S1.**
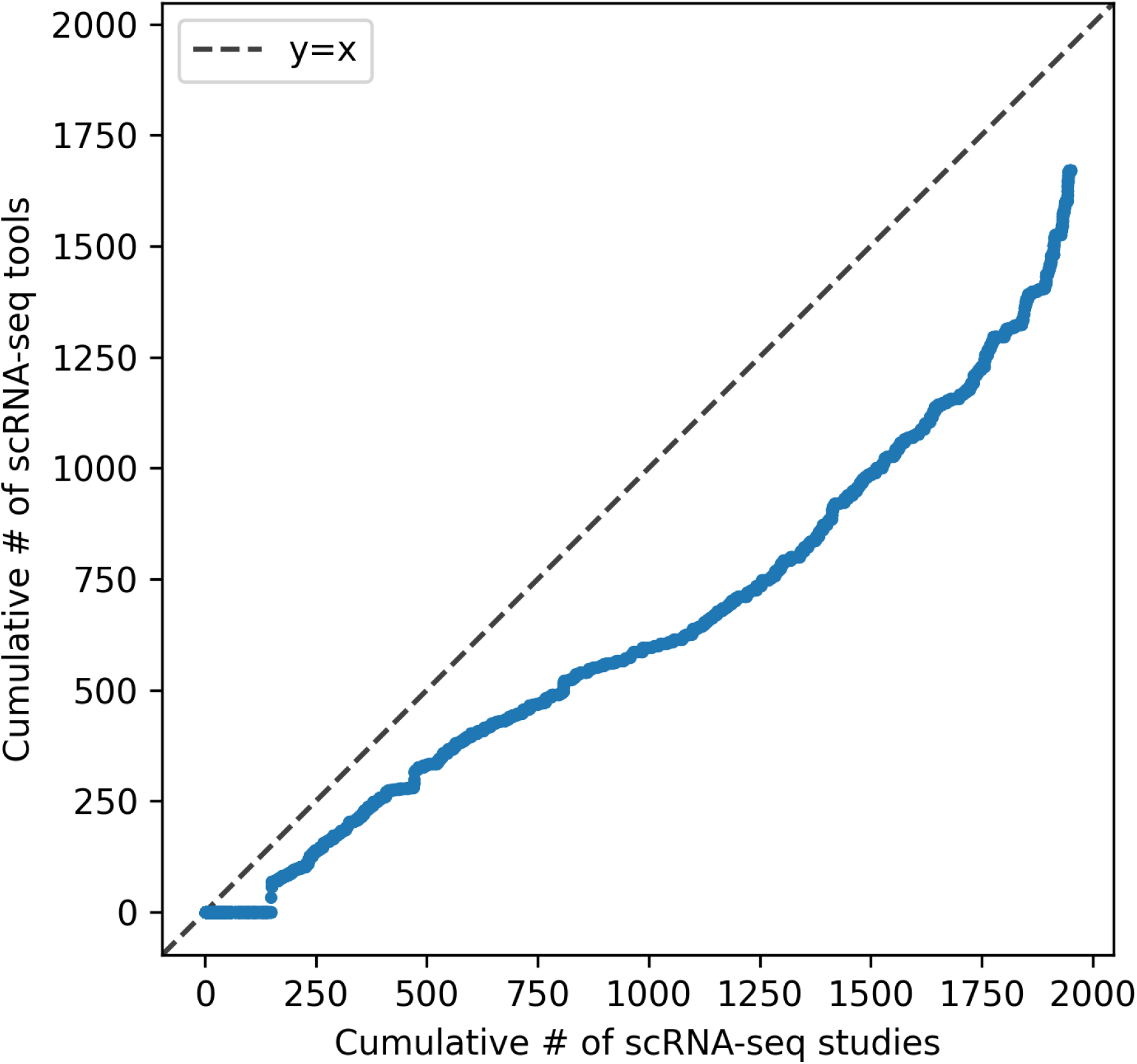
The increase in the number of published scRNA-seq tools almost keeps up with the number of published scRNA-seq datasets. This plot was created using code adapted from https://pachterlab.github.io/kallistobustools/tutorials/scRNA-seq_intro/python/scRNA-seq_intro/ and using data from [7] and [8]. The red line is a linear fit to the data, the black diagonal indicates *y* = *x*.

**Figure S2.**
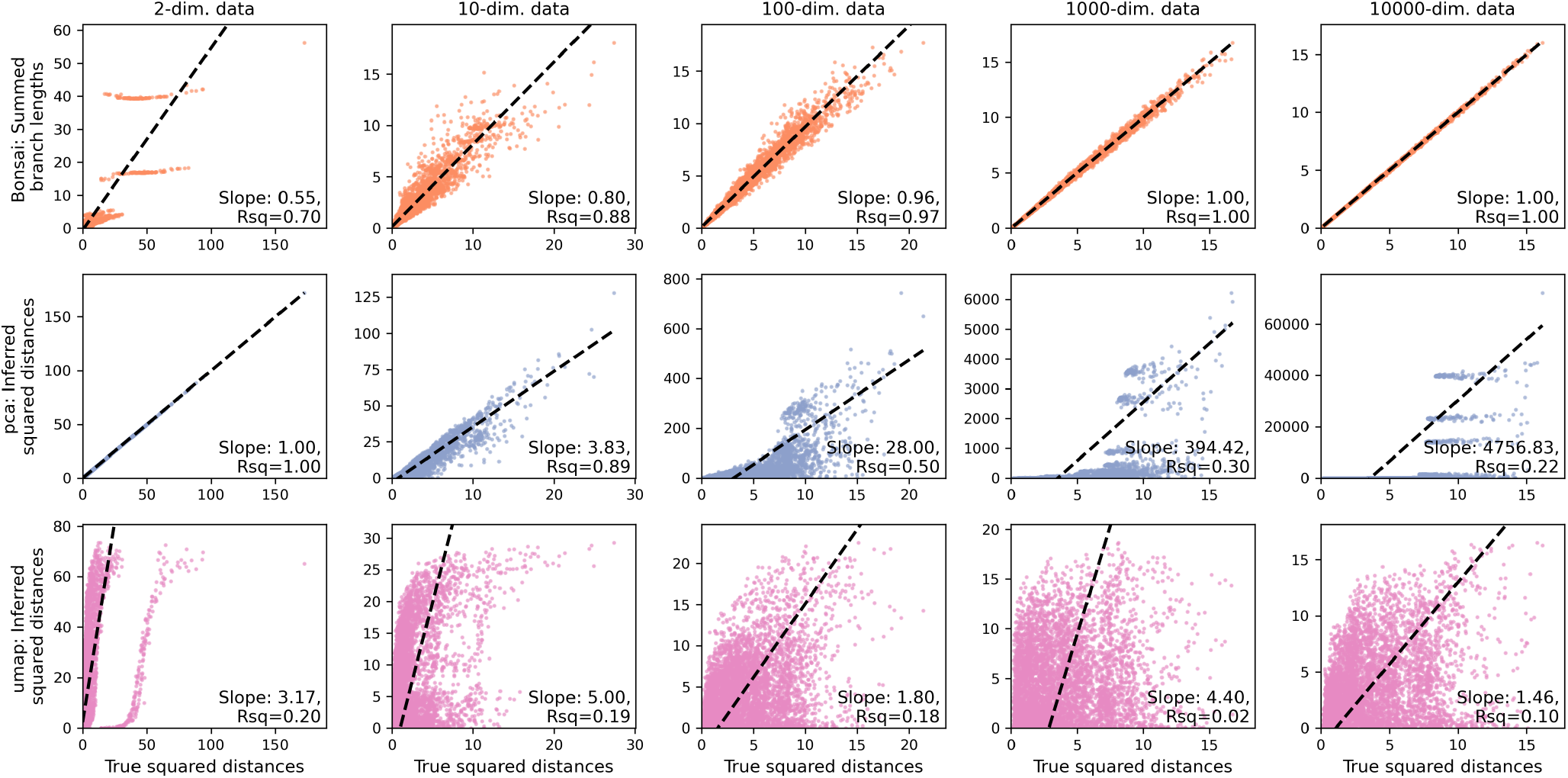
With increasing dimensionality, tree-structures get better at representing distances while other visualizations get worse. We created a simulated dataset by picking 100 cells from a normal distribution for which the dimensionality is varied in the different columns. For a gene-cell pair, (*g, c*), we draw the datapoint from a spherical Gaussian with variance *v*_*gc*_ = *v*_*g*_*v*_*c*_. Here, the gene variance *v*_*g*_ is drawn from an exponential distribution with mean 2. The cell variances, *v*_*c*_, are drawn by first sampling uniformly from a log-scale between 0.1 and 10 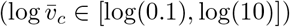, and then normalizing such that the average *v*_*c*_ is one: 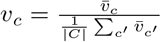 In the first row, we show that a tree-reconstruction can approximate the distances better and better at higher dimensionality, while the second and third rows show that PCA-projection and UMAP-embedding become worse at representing the distances.

**Figure S3.**
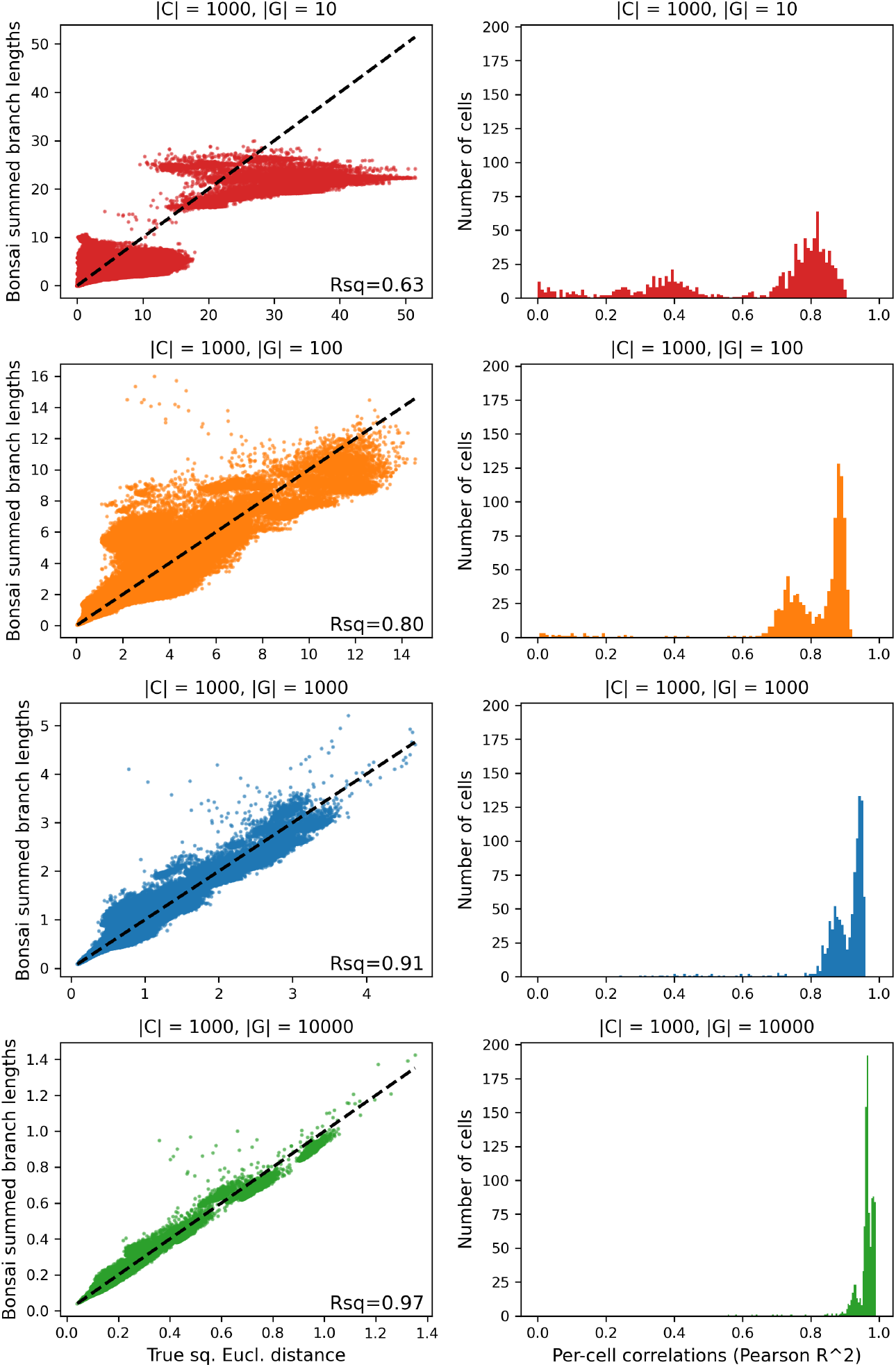
Distances between gene expression values from a real dataset can be accurately captured using a tree structure. To test whether cell-to-cell distances in real datasets can be captured accurately in a tree structure, we mimicked the analysis from Figure S2 using the gene expression values from a dataset of cord blood cells [22]. We normalized the scRNA-seq data using *Sanity*, and randomly selected 1000 cells. We then created several datasets by selecting the 10, 100, 1000 or 10 000 genes with the highest signal-to-noise ratio, as reported by *Sanity*. Finally, *Bonsai* was used to reconstruct a tree that connects the 1000 cells. In the left column, we show the true squared Euclidean distances between all pairs of cells, versus the sum of the branch lengths that connect these cells on the *Bonsai* tree. The dashed line is the *y* = *x*-line, and the Pearson R-squared value is reported in the plots. For the right column, we calculate for each cell the Pearson *R*^2^-value between true and inferred distances to the other 999 cells. The histograms show the distribution of these *R*^2^-values.

**Figure S4.**
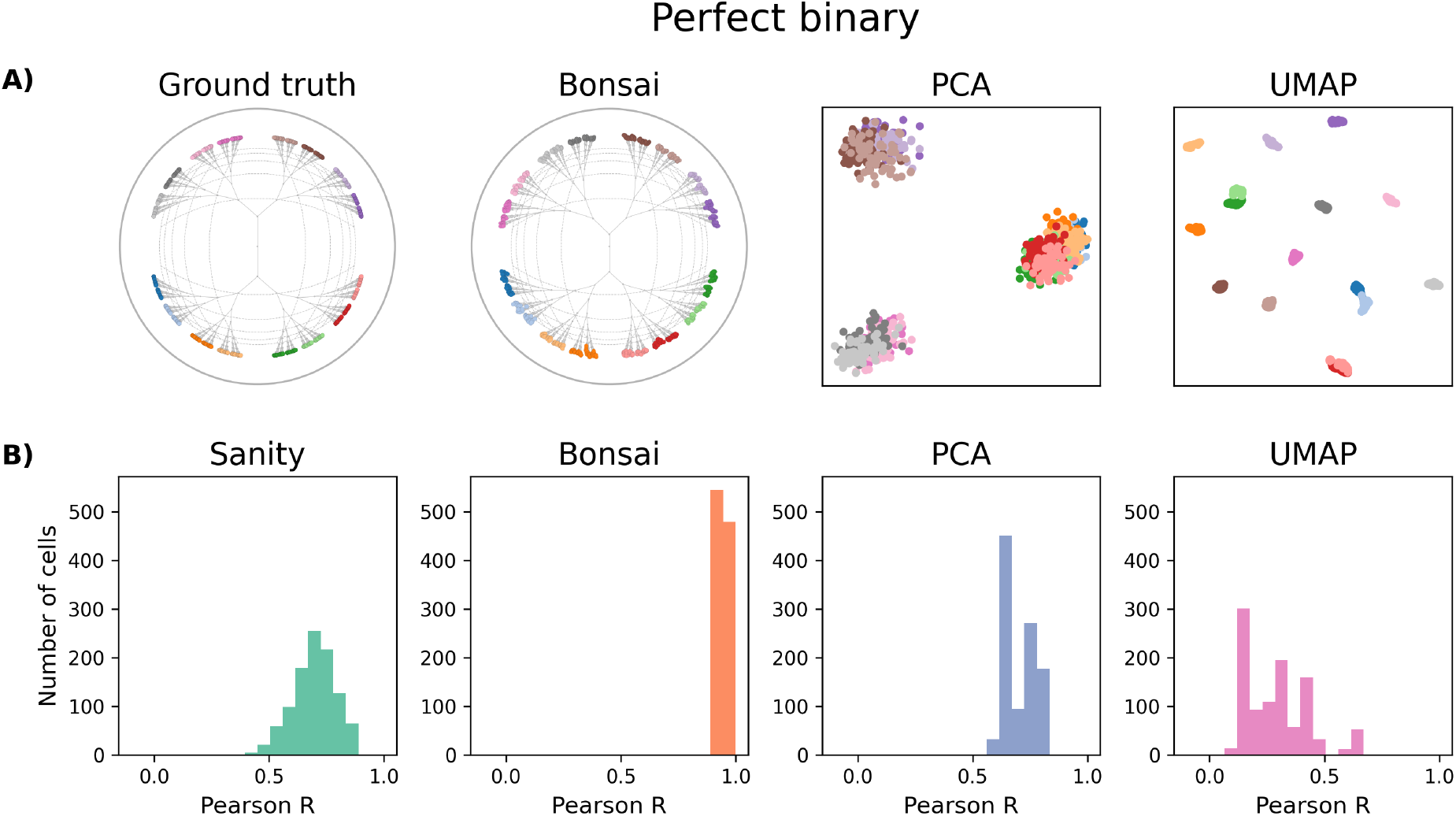
Visualization and pairwise distance recovery on a simulated dataset based on a perfectly binary tree. The ground truth tree is shown in the top-left panel, see Section SI.E for information on the simulation. The *Bonsai* visualization is shown in the equal-daylight layout on the hyperbolic disk. For PCA and UMAP, a standard logp1-normalization was followed by PCA-projection to 2 and 10 dimensions. UMAP-embedding into two dimensions was then run with parameters: `n neighbors=15`, `min dist=0.1`, `n components=2`, `metric=‘euclidean’`. Since *Sanity* is not a visualization tool, we cannot show a visualization. In the second row, we capture in a histogram how well cell-to-cell distances are represented across cells. Specifically, for each cell we compute the Pearson-R value for the correlation between the true and inferred distances between the cell and all other cells. For *Sanity* we used the specialized distance-calculation program, see [28], while taking only genes with a signal-to-noise ratio larger than 1 into account.

**Figure S5.**
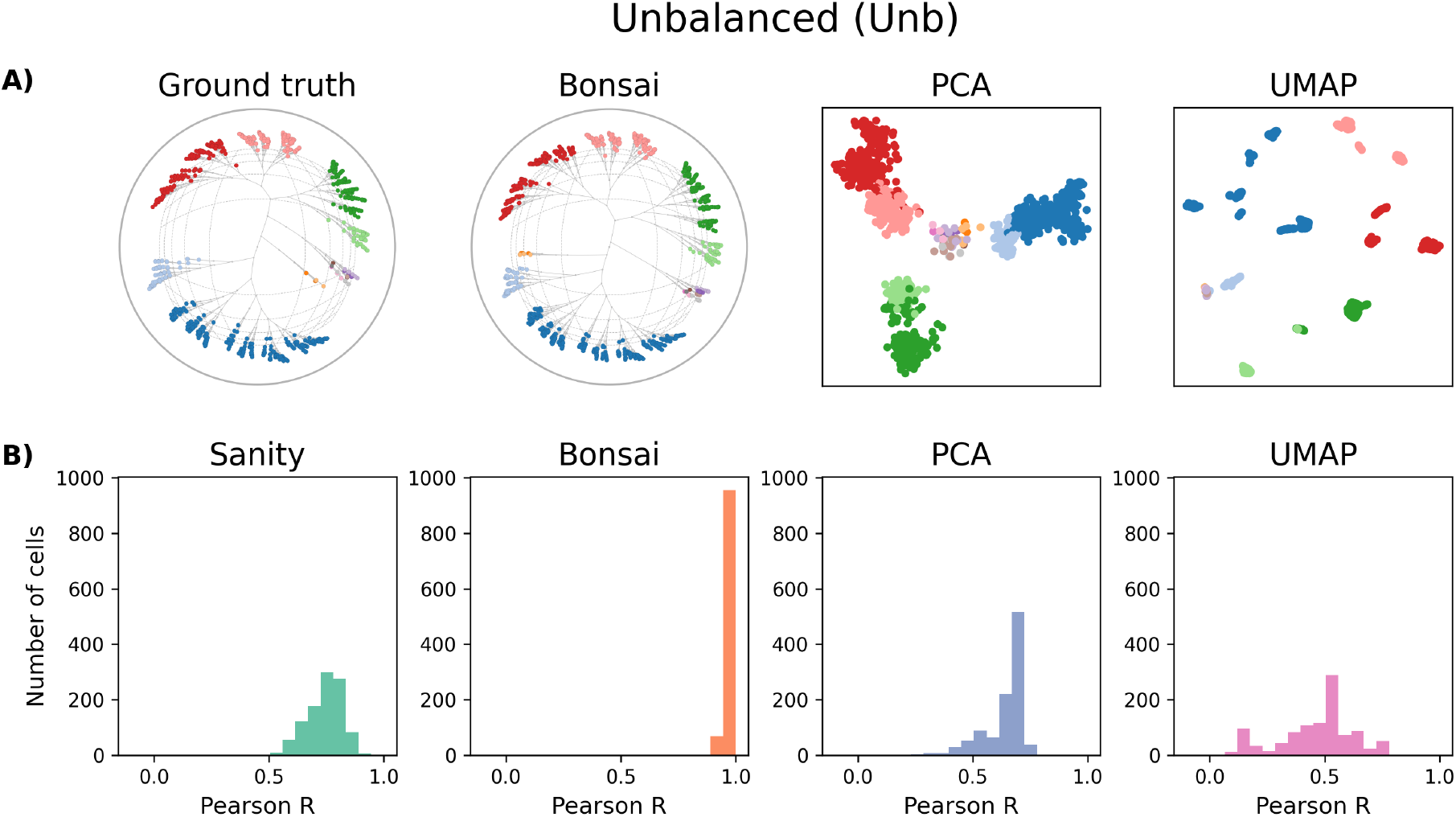
Visualization and pairwise distance recovery on a simulated dataset based on an unbalanced tree. The ground truth tree is shown in the top-left panel, see Section SI.E.2 for information on the simulation. For further information on this figure, see the caption of Figure S4.

**Figure S6.**
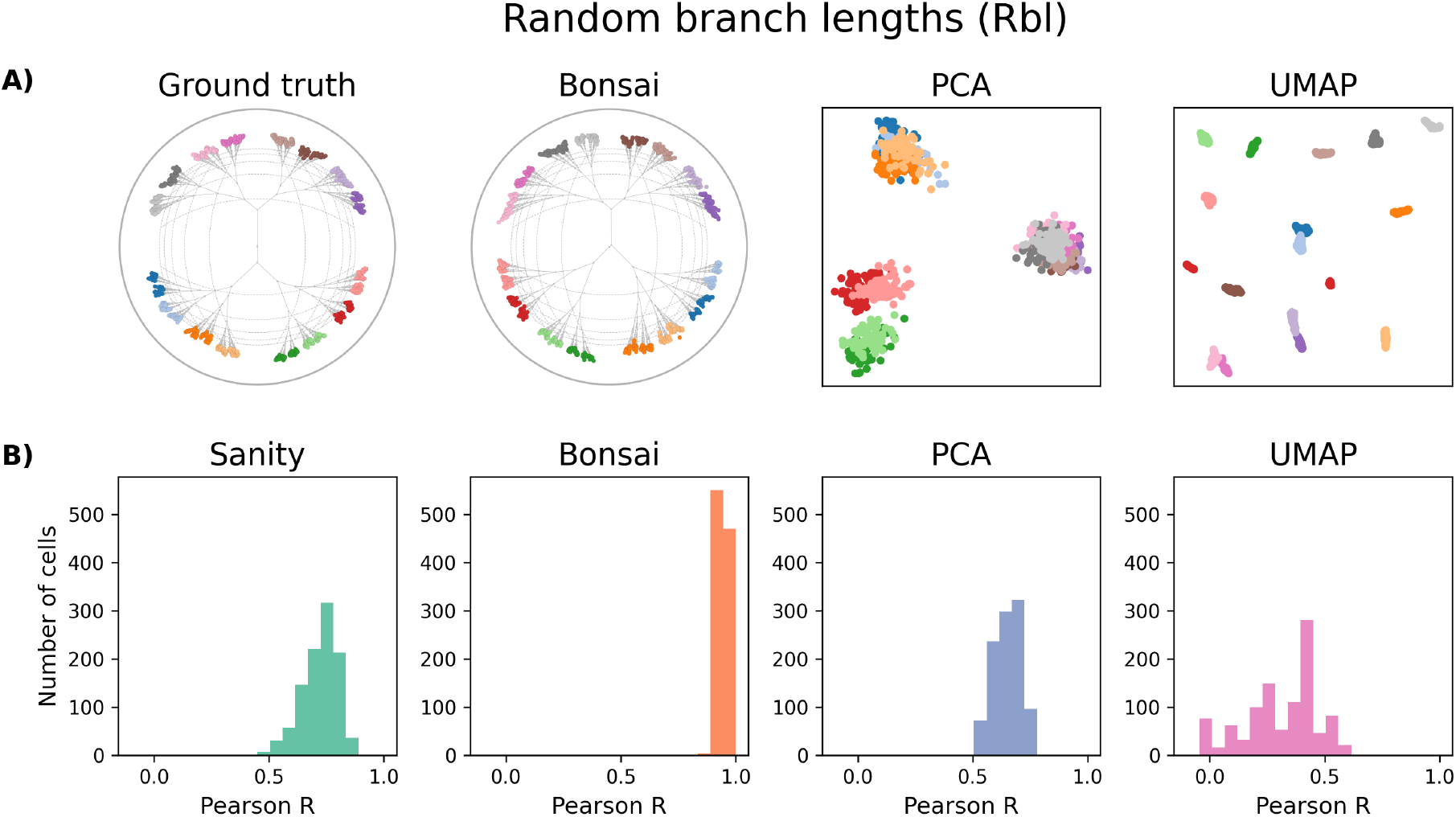
Visualization and pairwise distance recovery on a simulated dataset based on a binary tree with random branch lengths. The ground truth tree is shown in the top-left panel, see Section SI.E.2 for information on the simulation. For further information on this figure, see the caption of Figure S4.

**Figure S7.**
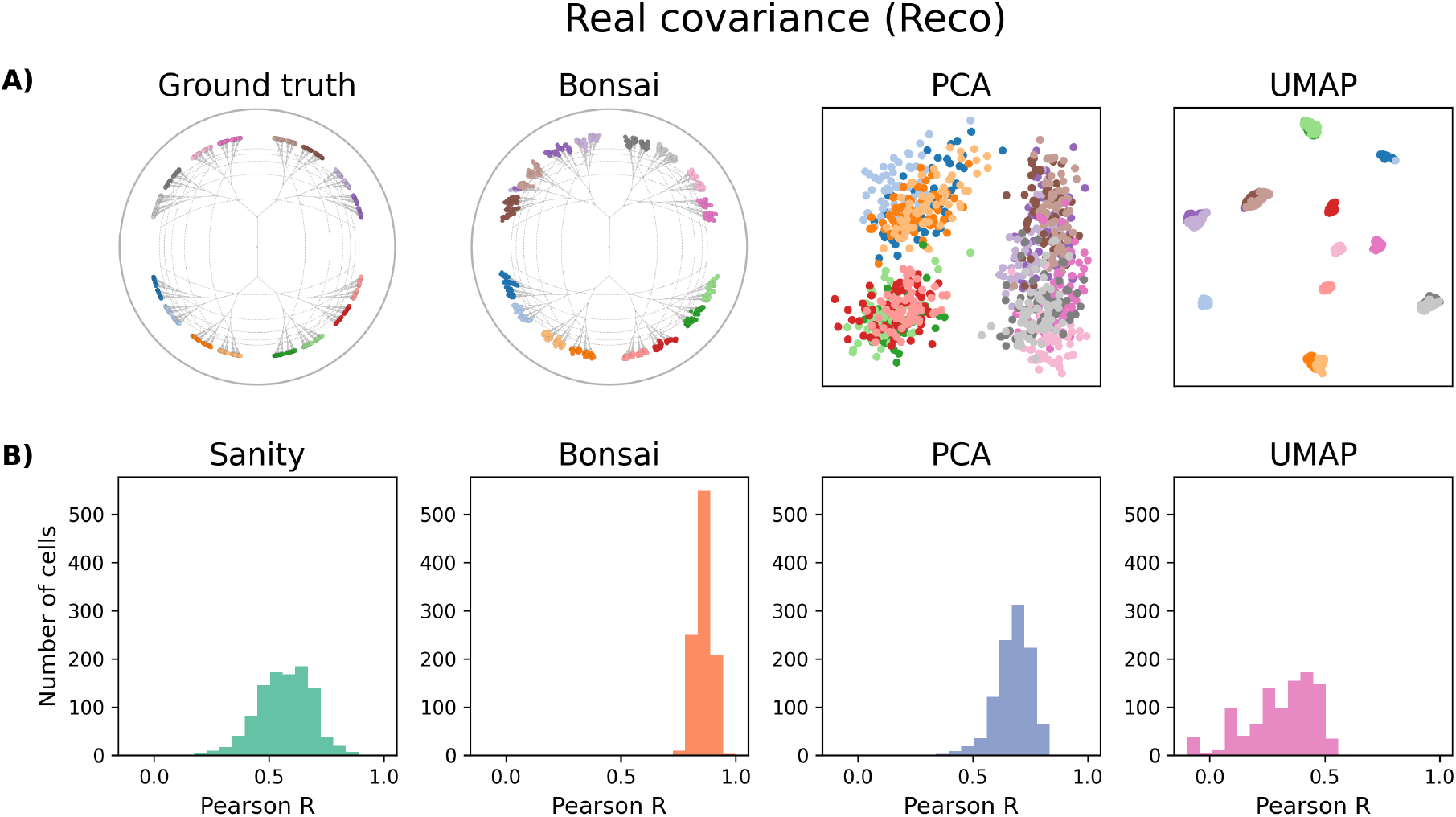
Visualization and pairwise distance recovery on a simulated dataset based on a binary tree using diffusion according to a covariance matrix estimated from a real dataset. The ground truth tree is shown in the top-left panel, see Section SI.E.2 for information on the simulation. For further information on this figure, see the caption of Figure S4.

**Figure S8.**
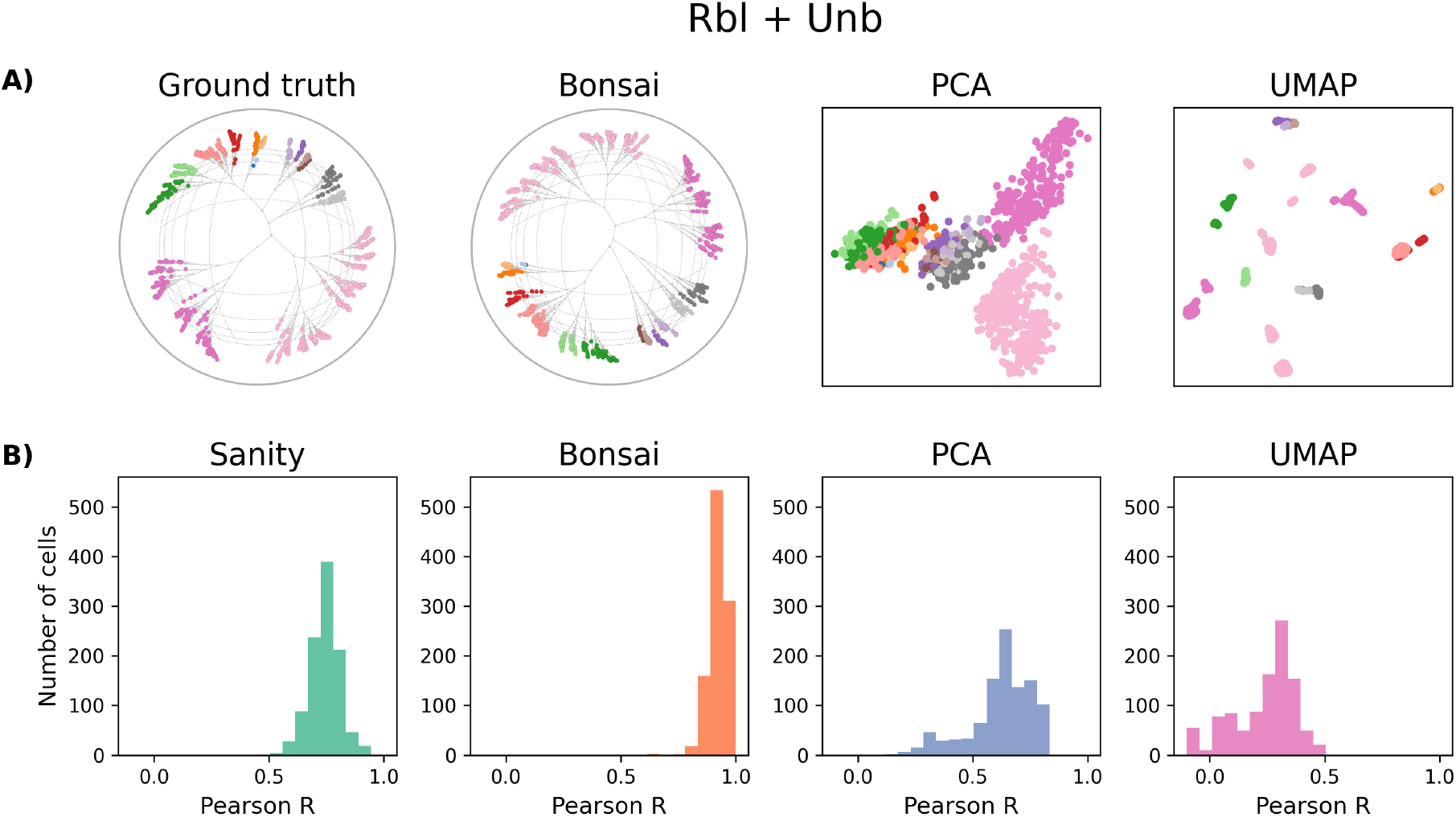
Visualization and pairwise distance recovery on a simulated dataset based on an unbalanced tree with random branch lengths. The ground truth tree is shown in the top-left panel, see Section SI.E.2 for information on the simulation. For further information on this figure, see the caption of Figure S4.

**Figure S9.**
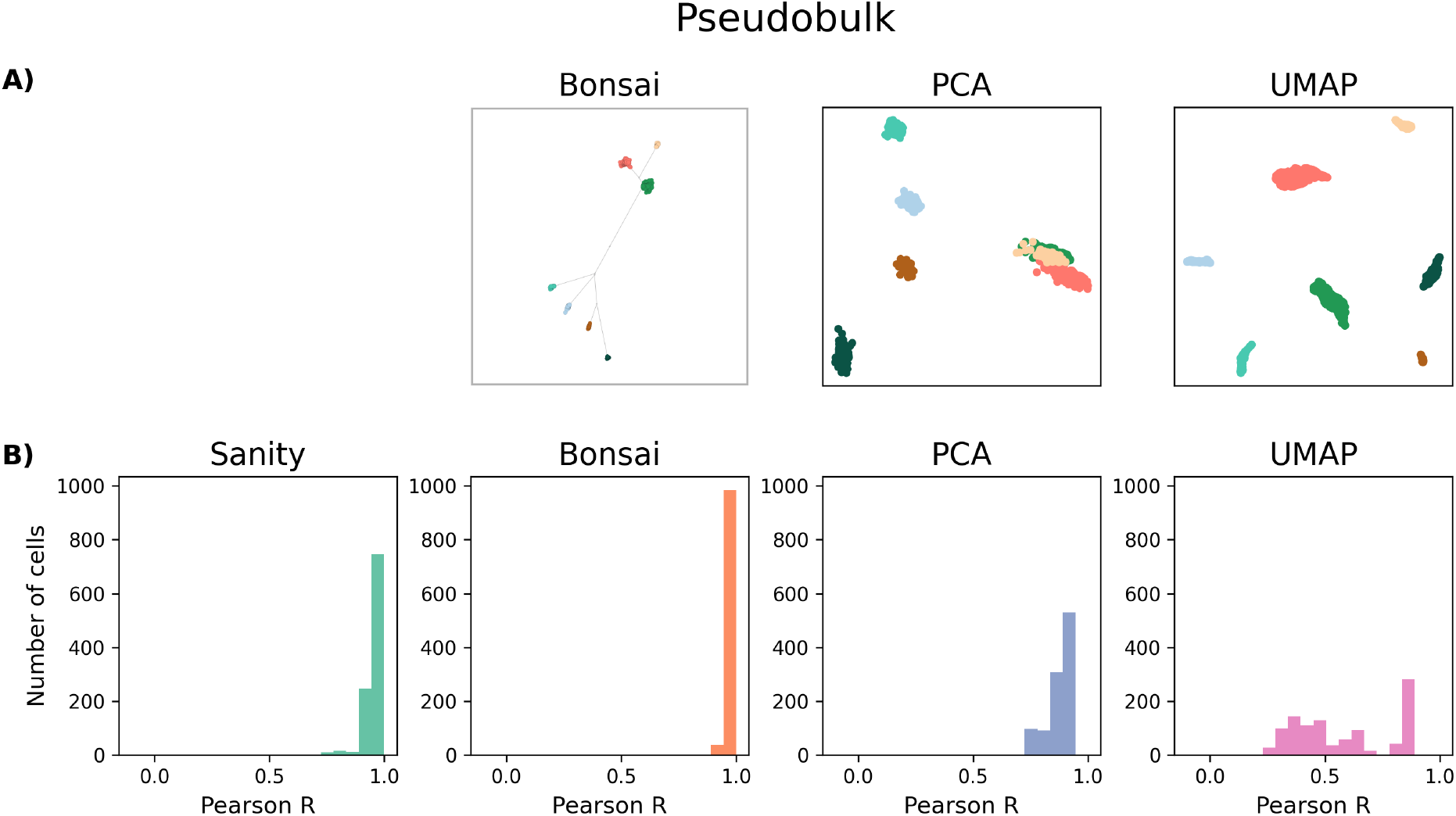
Visualization and pairwise distance recovery on a simulated dataset based on a pseudobulk dataset. Note that there is no ground truth tree for this simulation. See Section SI.E.2 for information on the simulation. For further information on this figure, see the caption of Figure S4.

**Figure S10.**
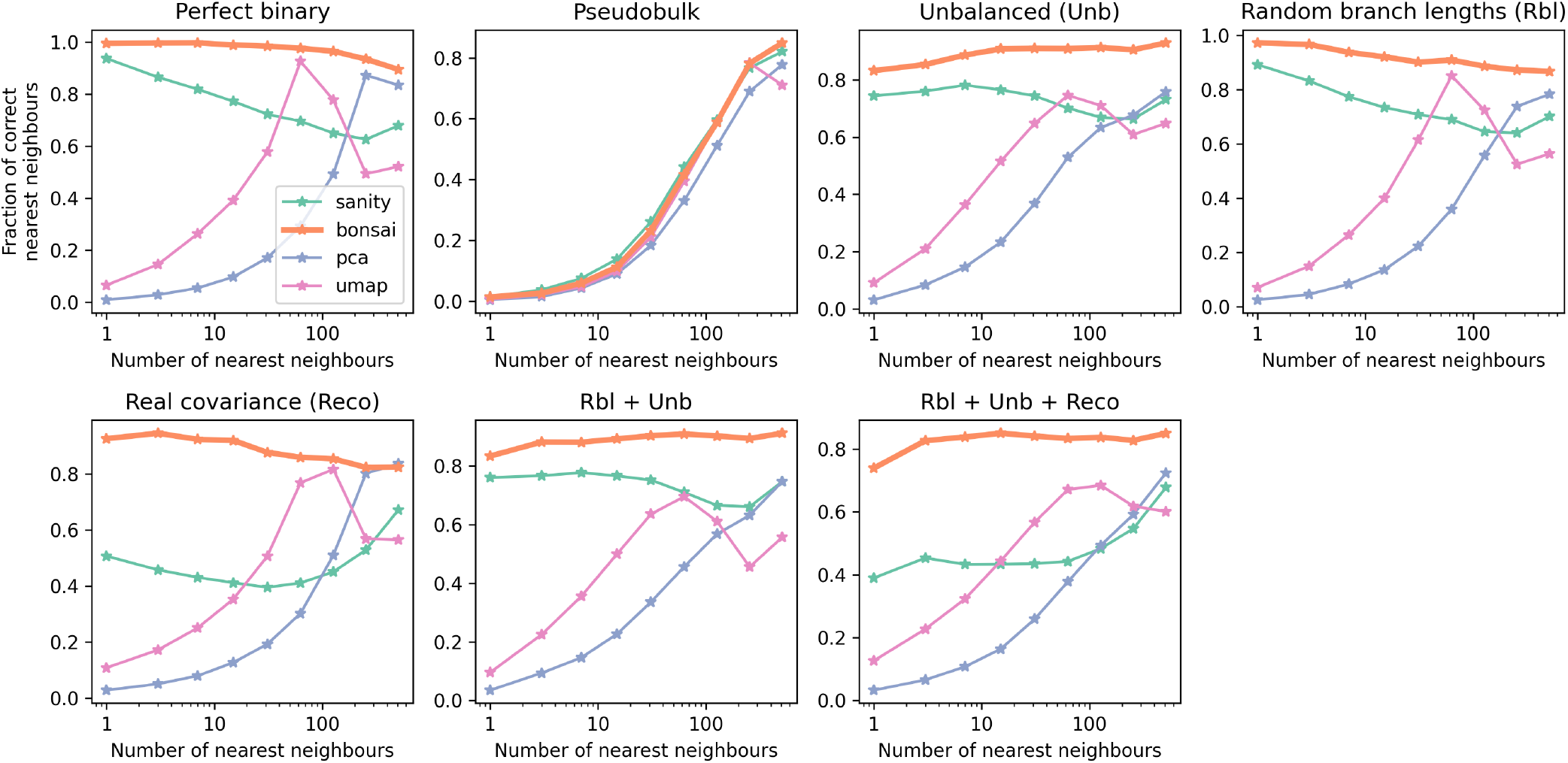
*Bonsai* outperforms other methods on identifying nearest-neighbor cells in scRNA-seq datasets. For the 7 simulated datasets described in SI.E, we show how well various methods identify the nearest neighbors of each cell. Specifically, for different number of nearest-neighbors *k*, we calculate what fraction of the inferred *k* nearest-neighbors are indeed among the *k* nearest-neighbors in the ground truth and show these fractions as a function of *k*. Each panel corresponds to one synthetic dataset (described at the top). For PCA and UMAP, a standard logp1-normalization was run followed by PCA-projection to 2 and 10 dimensions. UMAP-embedding into two dimensions was then run with parameters: `n neighbors=15`, `min dist=0.1`, `n components=2`, `metric=‘euclidean’`. For *Sanity* we used the specialized distance-calculation program (see [28]), using genes with a signal-to-noise ratio larger than 1. For the Pseudobulk-dataset, all methods perform equally bad because no clear nearest-neighbor structure is present in this dataset. For all other datasets, *Bonsai* strongly outperforms both PCA and UMAP for virtually all values of *k*. Notably, PCA tends to perform well only for large *k*, whereas UMAP typically has a local peak in performance for a value of *k* somewhere between 30 and 100, reflecting that this method is optimized for identifying nearest-neighbors at this particular scale only. More surprisingly, *Bonsai* also outperforms *Sanity* on all datasets, even though both methods are based on the same inferred gene expression values. This suggest that *Bonsai*’s increased performance results from implicitly using the tree-structure as a prior for estimating distances.

**Figure S11.**
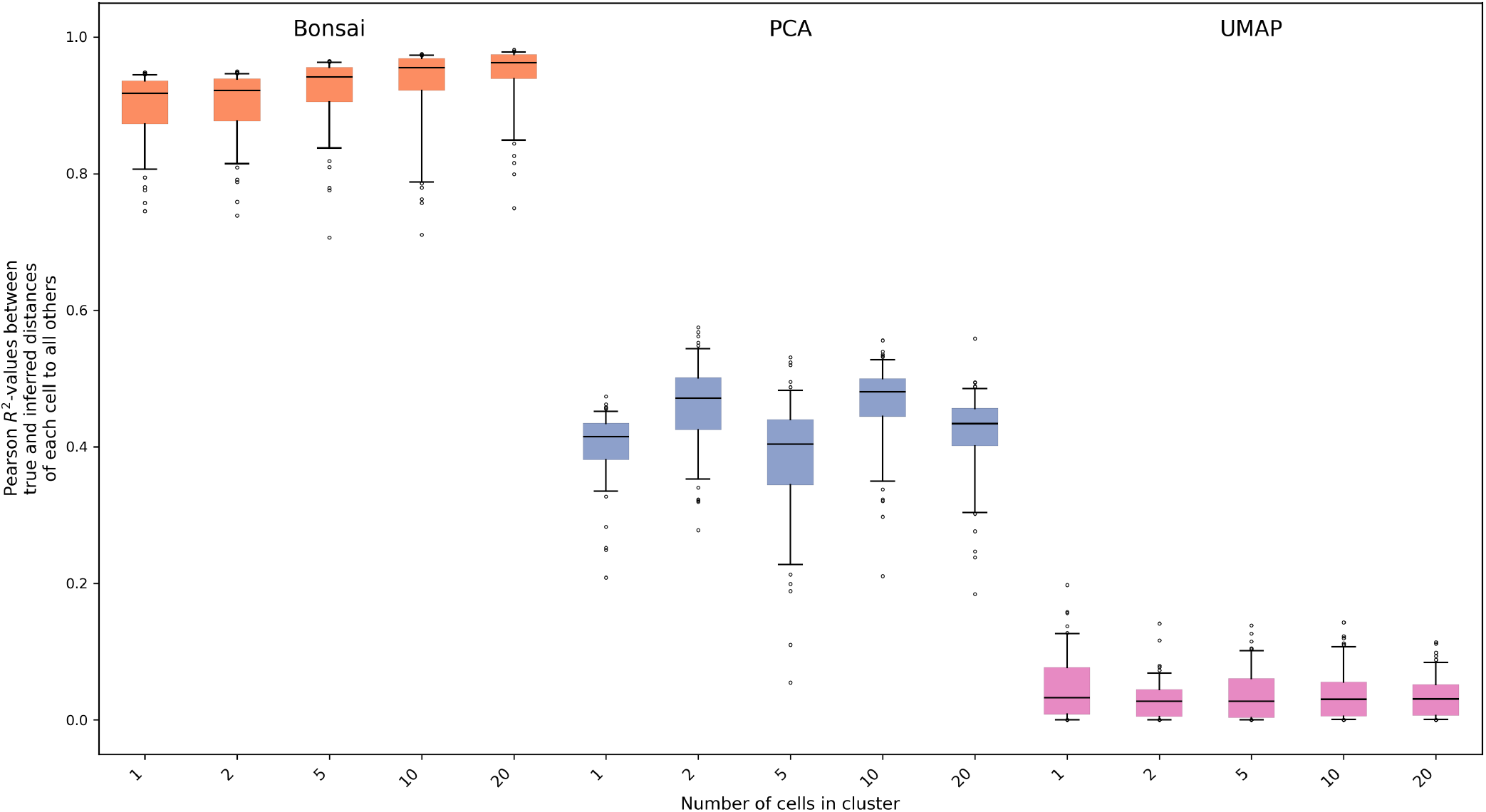
Tree-structures automatically use nearby cells to get more accurate distance estimates. We created synthetic datasets by first taking the 100 points in the dimensional simulated dataset from Fig S2 and using these as cluster-centers. We then sampled either 1, 2, 5, 10, or 20 cells around each cluster-center by adding a noise term drawn from a normal distribution in 100 dimensions with variance 2.5 in each dimension, see Section SI.F.2 for details. We thus created datasets in which each cell has both a *measured position* (i.e. with the noise added) and a *true position* (i.e. equal to its corresponding cluster-center). We used *Bonsai*, PCA and UMAP to construct a representation on the measured positions, and then checked how well these representations capture the pairwise distances between the *true positions*. In particular, we selected only the first cell around each cluster-center from each representation and calculated the Pearson *R*^2^-value between true and inferred distances to the other 99 cells for each of the 100 cells. The boxplots summarize the distribution of these squared correlation coefficients. We see that, apart from generally dramatically outperforming both PCA and UMAP, *Bonsai*’s tree reconstruction approximates the distances from each cell to its neighbors better and better as the number of cells per cluster increases. In contrast, for PCA and UMAP this effect is completely absent. This further supports that the tree structure acts as a regularizer to improve the estimates of each cell’s true position.

**Figure S12.**
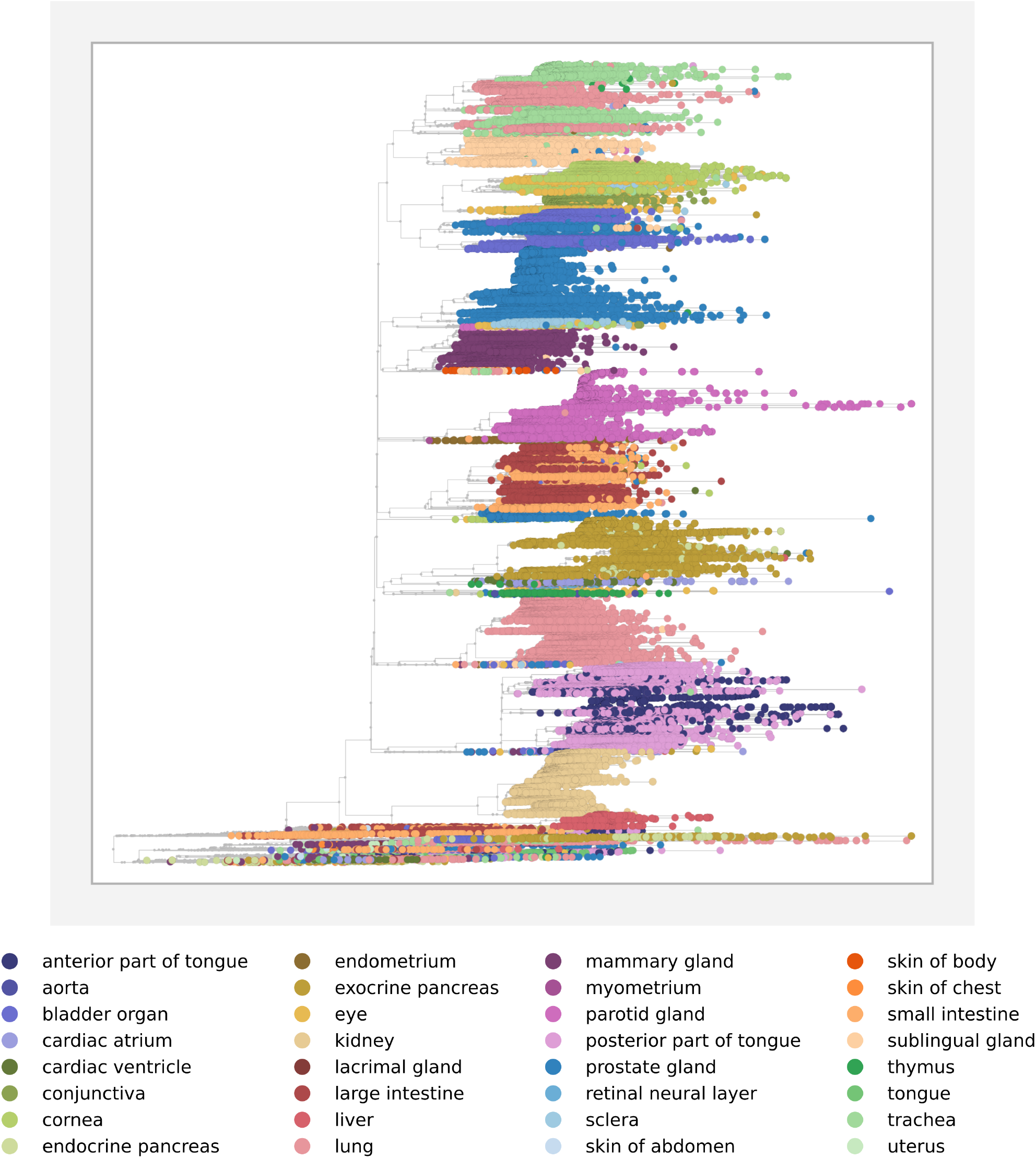
Bonsai reconstruction of 104 148 epithelial cells from Tabula Sapiens [5]. We show the *Bonsai* tree reconstructed on the subset of all epithelial cells from the *Tabula Sapiens* atlas in the dendrogram layout. Different colors indicate the different tissues that the cells originate from. We have made the *Bonsai* -results available for exploration using *Bonsai-scout* under https://bonsai.unibas.ch/bonsai-scout/?dir=Epithelial_cells_from_Tabula_Sapiens as a community resource.

**Figure S13.**
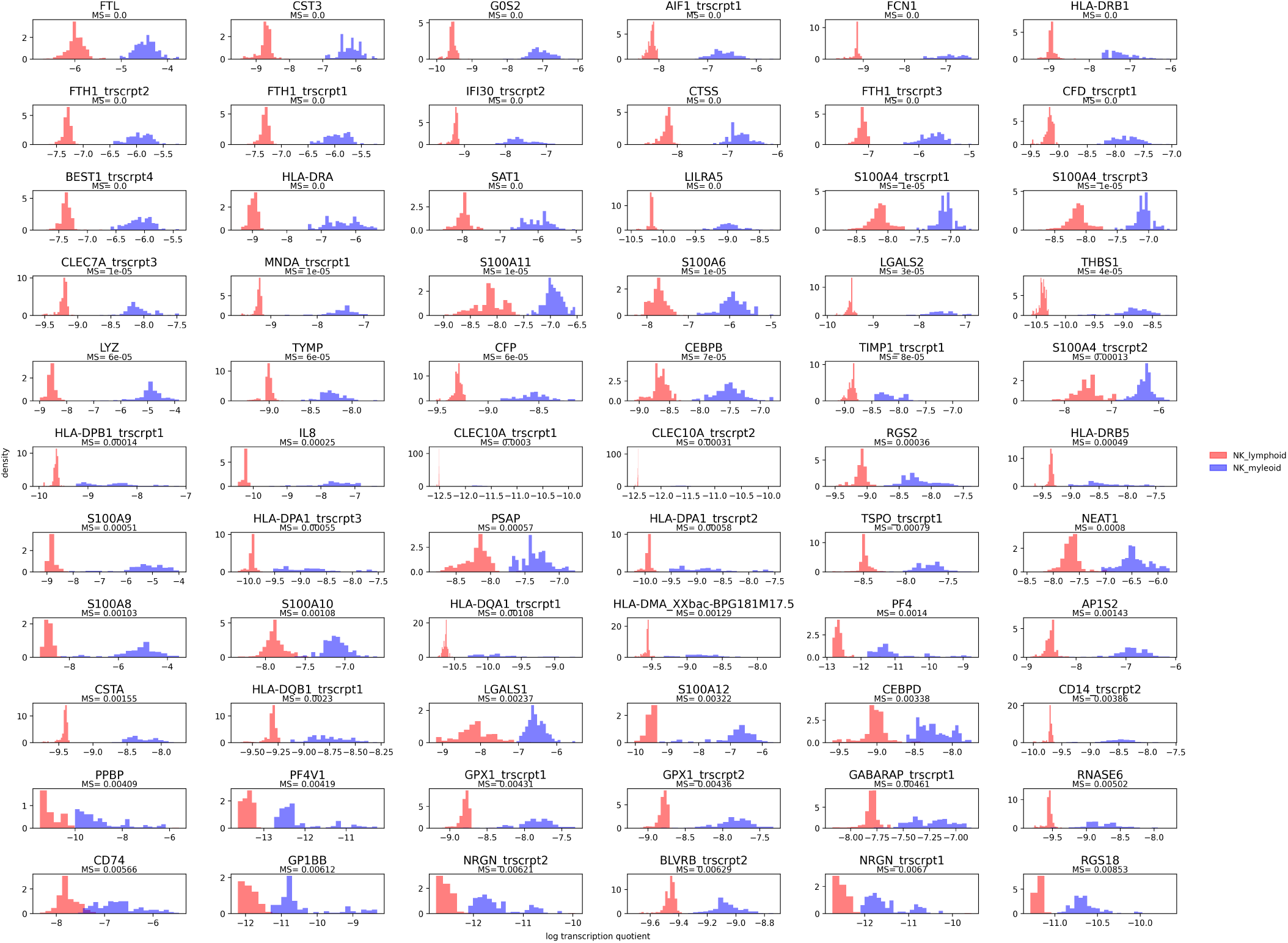
Marker genes that are consistently higher expressed in myeloid-NK cells than lymphoid-NK cells. For the two subpopulations of NK-cells that were identified by the *Bonsai* representation of the blood cell dataset [22] (see Fig 4), we found all marker genes with a marker score MS below 0.01, meaning that the probability that the gene is higher expressed in a randomly chosen lymphoid-NK cell than in a randomly chosen myeloid NK cell is less than 0.01 (see Methods). Each panel corresponds to one marker gene or transcript (shown above its marker score at the top of the panel). The red and blue histograms show the distribution of expression levels (log transcription quotients) in the lymphoid-NK and myeloid-NK cells, respectively. Note that our pre-processing pipeline by default associates expression levels by groups of transcripts that are described from the same promoter so that one gene can give rise to multiple transcript classes, which are indicated by a suffix such as `trscrpt1`.

**Figure S14.**
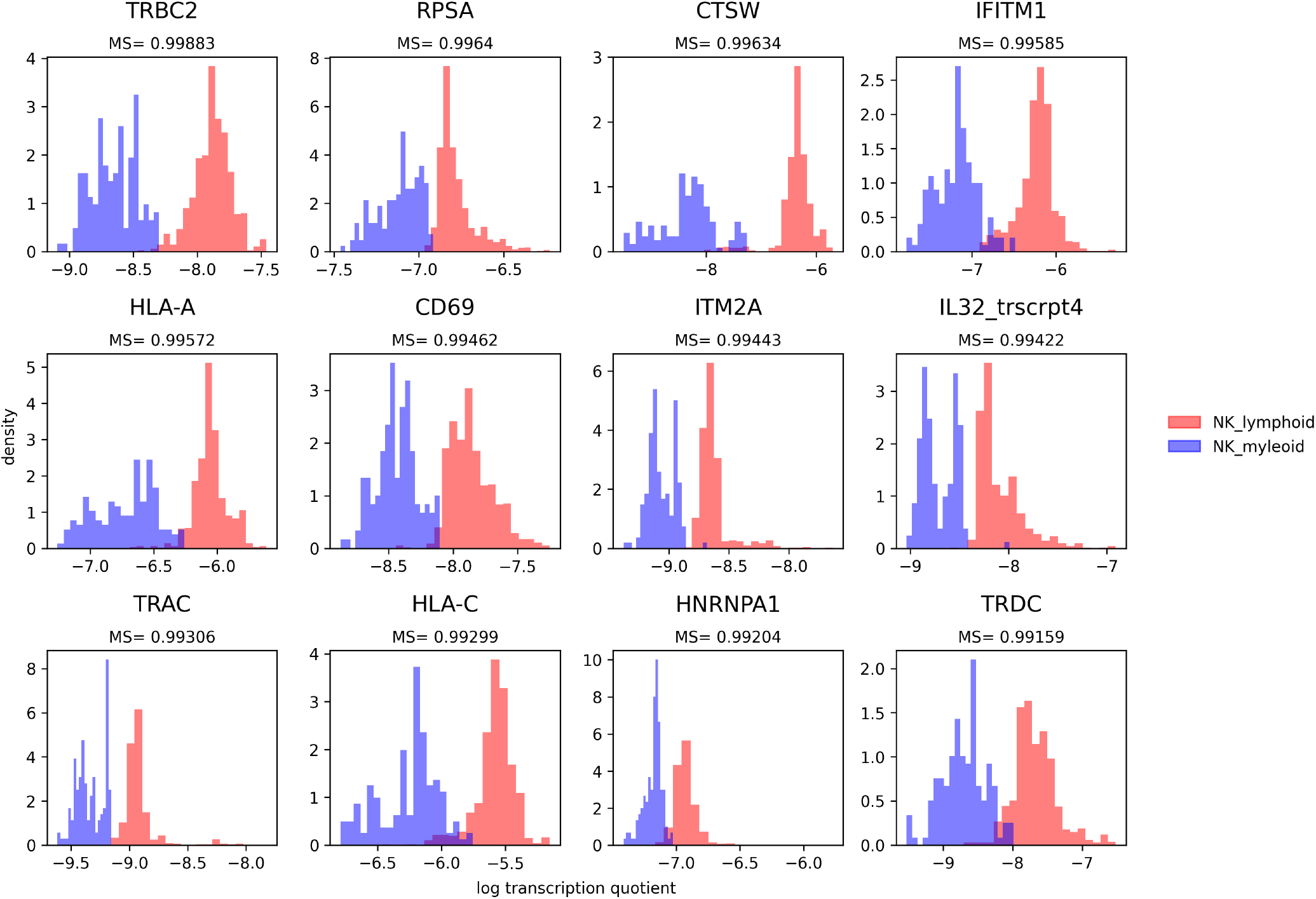
Marker genes that are consistently higher expressed in lymphoid-NK cells than myeloid-NK cells. For the two subpopulations of NK-cells that were identified by the *Bonsai* representation of the blood cell dataset [22] (see Fig 4), we found all marker genes with a marker score MS above 0.99, meaning that the probability that the gene is higher expressed in a randomly chosen lymphoid-NK cell than in a randomly chosen myeloid NK cell is more than 0.99 (see Methods). Each panel corresponds to one marker gene or transcript (shown above its marker score at the top of the panel). The red and blue histograms show the distribution of expression levels (log transcription quotients) in the lymphoid-NK and myeloid-NK cells, respectively. Note that our pre-processing pipeline by default associates expression levels by groups of transcripts that are described from the same promoter so that one gene can give rise to multiple transcript classes, which are indicated by a suffix such as `trscrpt1`.

**Figure S15.**
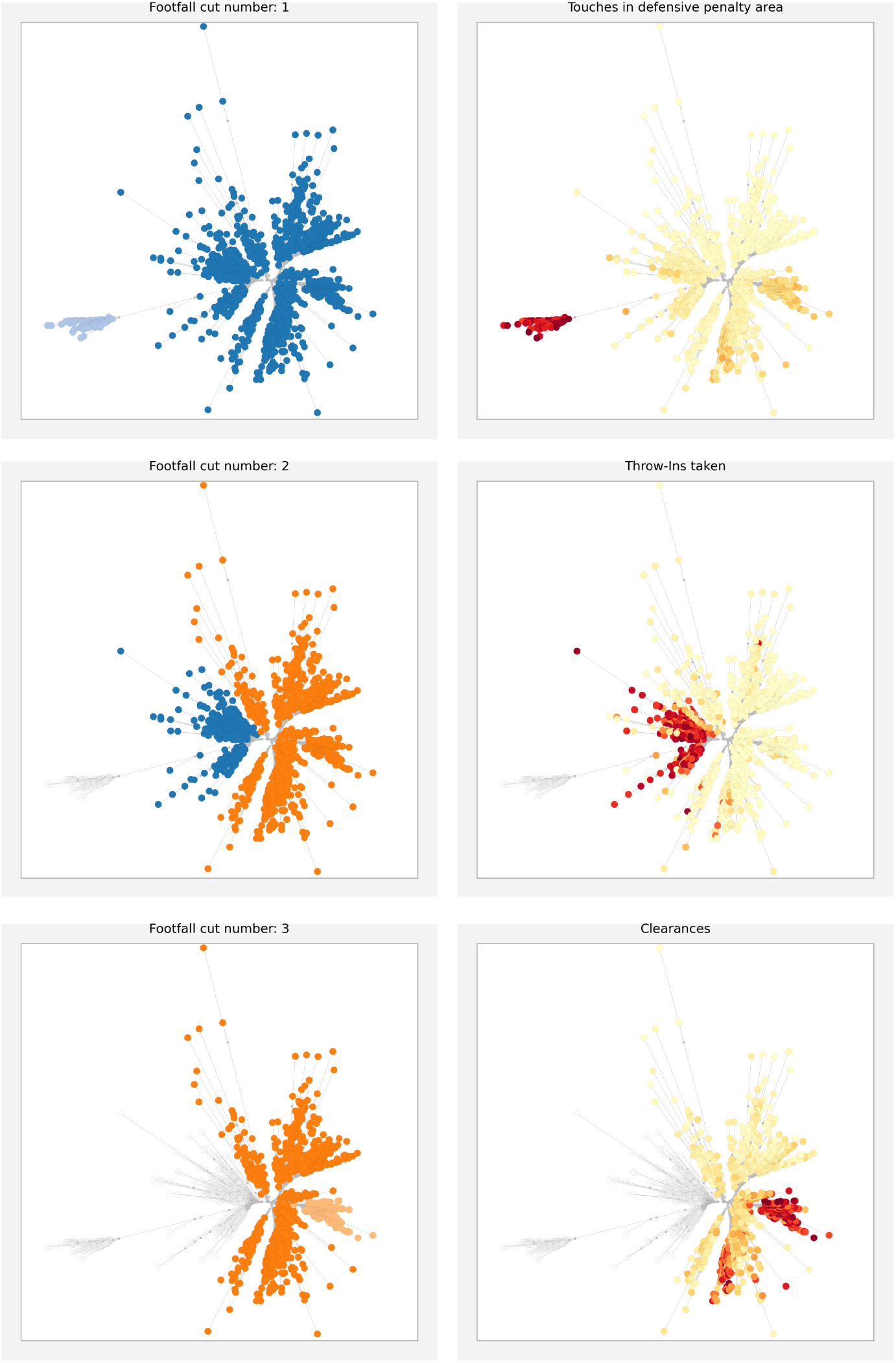
Marker genes distinguishing clusters in the football player dataset. For the dataset with statistics on 1436 football players [53], we found 12 player-clusters using our clustering method that iteratively cuts the tree into smaller subtrees (see Methods). The left panels show, from top to bottom, the first three cuts, and values of example marker features for each split are shown on the right. In the first cut, goalkeepers are split from other players with marker ‘Touches in defensive penalty area’. The second cut splits full backs and wing backs from the others with marker ‘Throw-ins taken’. The third cut splits center backs from the remaining players (i.e. the subtree without the goalkeepers and fullbacks) with marker the number of ‘Clearances’. Thus, after the first three cuts, goal keepers and defenders were split off from the rest.

**Figure S16.**
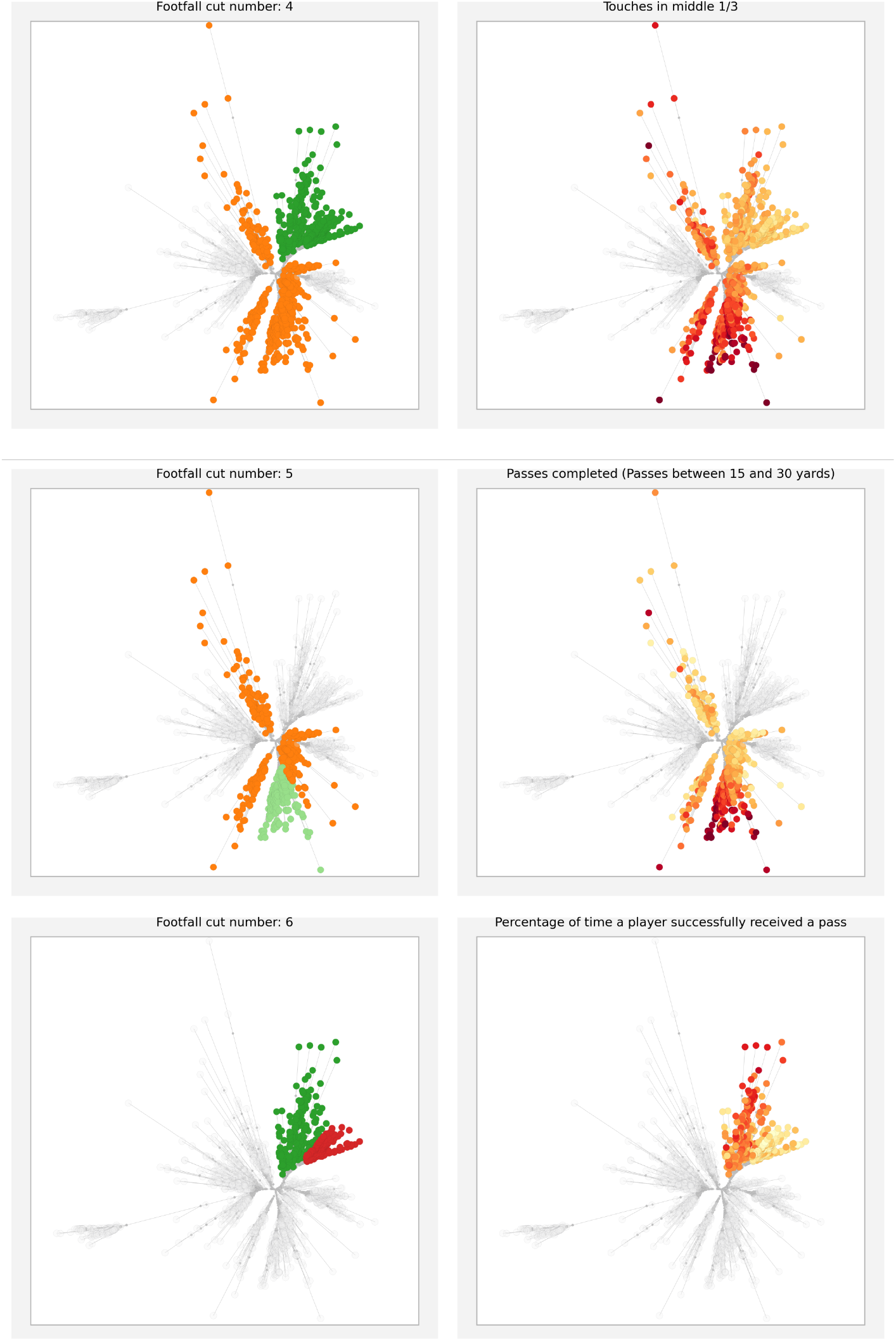
Marker genes distinguishing clusters in the football player dataset. For the dataset with statistics on 1436 football players [53], we found 12 player-clusters using our clustering method that iteratively cuts the tree into smaller subtrees (see Methods). The left panel shows, from top to bottom, cuts 4-6, and values of example marker features for each split are shown on the right. The fourth split cuts attackers from the midfielders with as the marker a lower number of ‘Touches in the middle’. The fifth cut splits a group of defensive midfielders and midfielding defenders from the other midfielders using as the marker feature the number of ‘Passes between 15 and 30 yards completed’. The sixth cut splits the attackers into deep attacking midfielders and pure attackers with marker that pure attackers have a lower percentage of successfully received passes.

**Figure S17.**
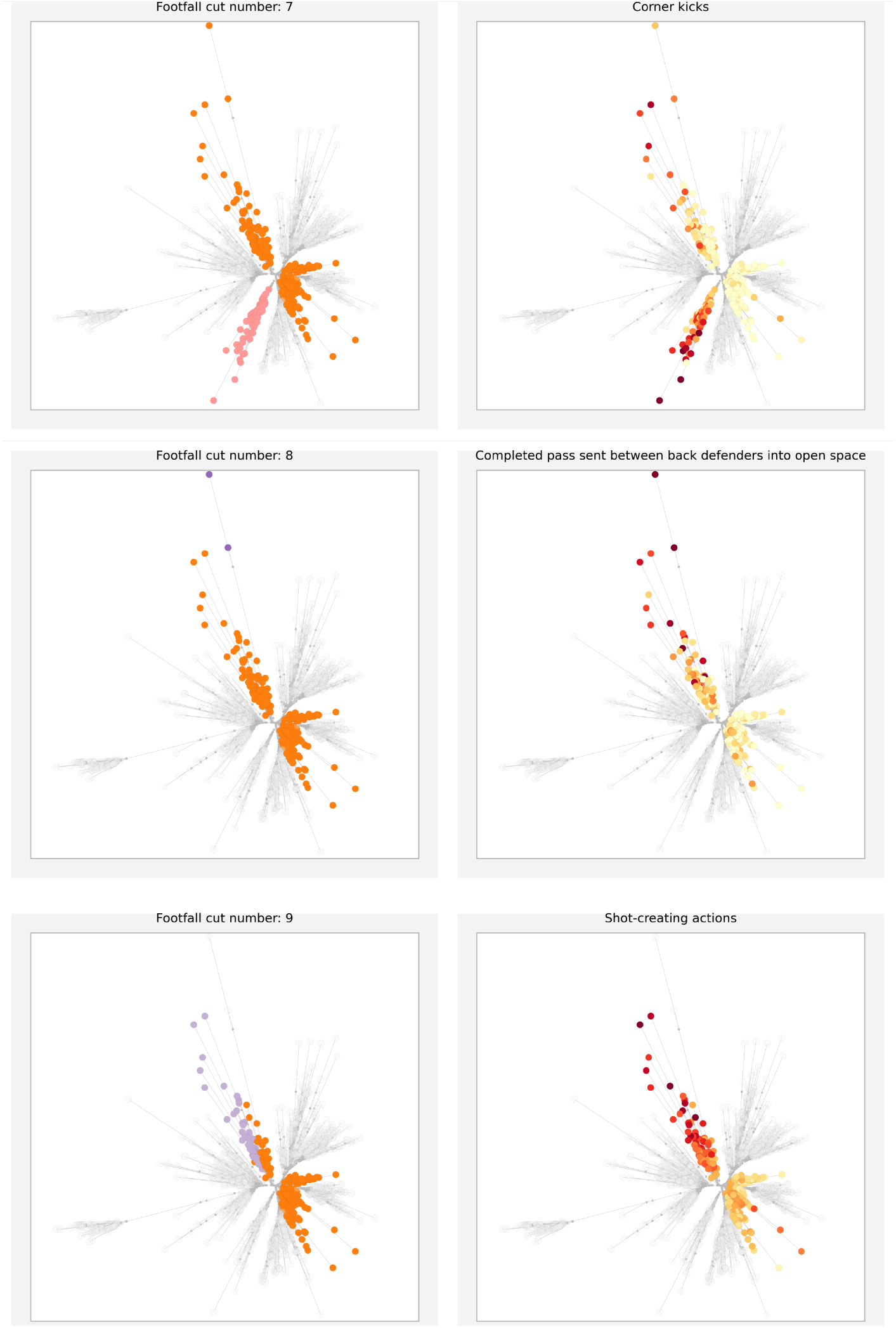
Marker genes distinguishing clusters in the football player dataset. For the dataset with statistics on 1436 football players [53], we found 12 player-clusters using our clustering method that iteratively cuts the tree into smaller subtrees (see the description in the Methods-section). The left panel shows, from top to bottom, cuts 7-9, and values of example marker features for each split are shown on the right. Cut seven splits a group of midfielders from the other midfielders and is characterized by taking corners more than others. The eighth cut splits just two outlier players (Messi and Neymar) from the rest, with marker the number of completed passes that are sent into open space between back defenders of the other team. The ninth cut splits off a group of attacking midfielders that is characterized by having many’shot-creating actions’.

**Figure S18.**
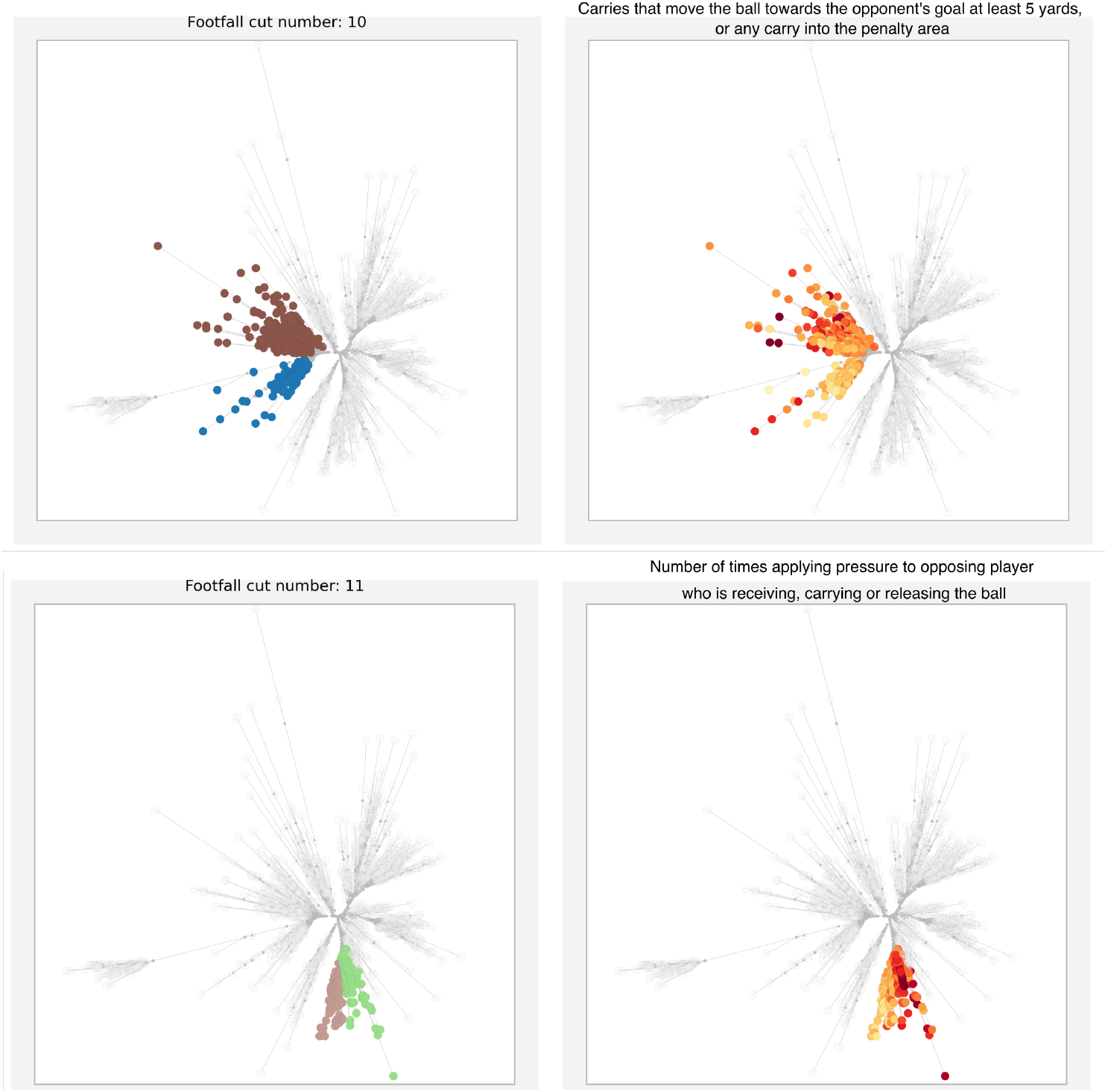
Marker genes distinguishing clusters in the football player dataset. For the dataset with statistics on 1436 football players [53], we found 12 player-clusters using our clustering method that iteratively cuts the tree into smaller subtrees (see the description in the Methods-section). The left panel shows, from top to bottom, cuts 10 and 11, and values of example marker features for each split are shown on the right. The tenth cut splits wing backs from fullbacks, with wing backs being characterized by more often moving the ball forward at least 5 yards. Finally, the eleventh cut splits what we have called midfielding defenders from defensive midfielders, where defensive midfielders are characterized by more often applying pressure to an opposing player that has the ball.

**Figure S19.**
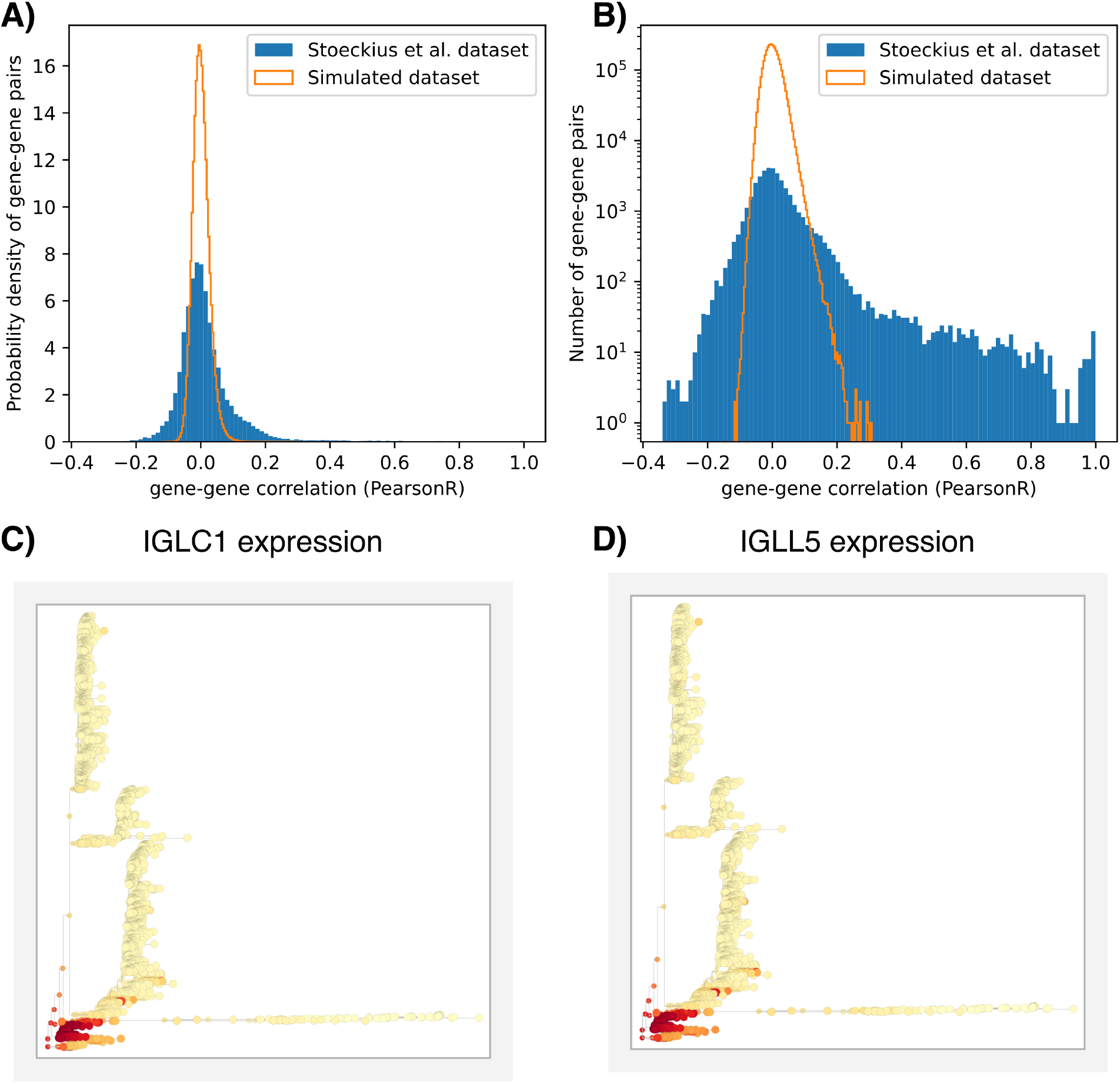
The homogeneous diffusion prior for gene expression changes does not inhibit *Bonsai* from detecting correlations between genes in the data. We can calculate the correlations in the inferred gene expression changes along the branches of the reconstructed *Bonsai* -trees. In **A)-B)**, we show these correlations for all gene-gene pairs for two datasets: 1) for the blood dataset by Stoeckius et al. [22] that was discussed in the main text, and 2) for the simulated dataset based on a perfect binary tree. The data in the simulated dataset are purely based on the homogeneous diffusion prior, and indeed we see only small correlations between genes that arise because we take a finite sample of expression changes. However, the real dataset shows much stronger correlations between genes, proving that the homogeneous diffusion prior is easily overruled by the data if there is evidence for the correlated change of genes. We show the same data in **A)** and **B)**, but in **A)** we show the normalized histogram, while in **B)** we show the y-axis on a log-scale. In **C)-D)**, we show an example of two strongly correlated genes, which indeed have a very similar expression pattern. Since these genes encode for parts of the immunoglobulins, which are produced by B-cells, it indeed makes sense that these are strongly correlated.

**Figure S20.**
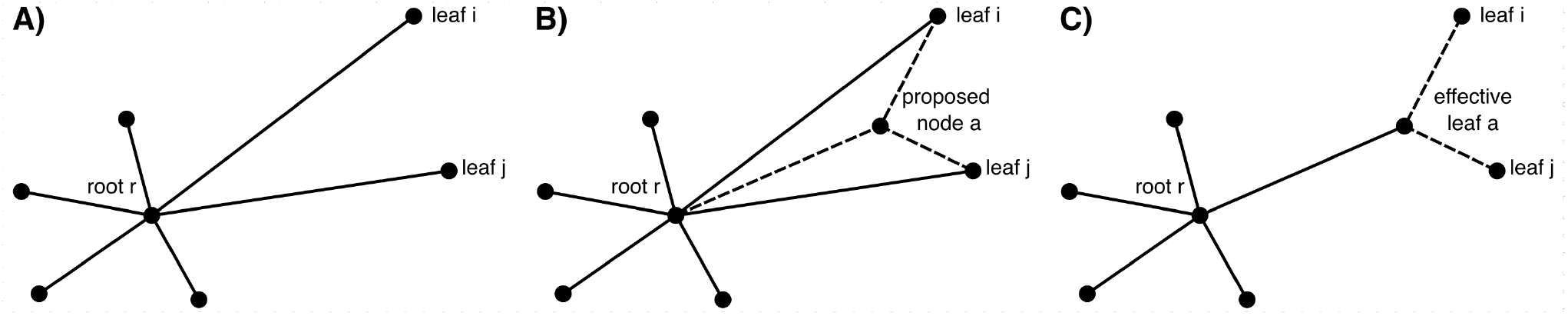
The iterative procedure of adding internal nodes to a star-tree. **A)** The iterative procedure starts from a star-tree in which all leaves are connected to a root-node. **B)** We can calculate the increase in tree likelihood that would result from adding an internal node to any pair of leafs. Here, we would add node *a* upstream of leafs *i* and *j*. Once we pick the optimal internal node to add, we can summarize the likelihood of the subtree downstream of *a* in the properties of an effective leaf *a*. Viewed from the root, we thus have a star-tree again, such that we can start again from **A)**.

## SI.B The likelihood of the data given a tree

In this section we derive the equations for calculating the likelihood of the data given a tree topology, *T*, and its branch lengths, ***t***: *P* (*D*|*T*, ***t***).

### Notational conventions

In the following, we will define a tree by its topology *T* of a set of nodes, connected by branches with branch lengths ***t***. Even though the likelihood that we will derive is independent of which node we designate as the root, we will for notational convenience require that we have defined which node is the root. For a rooted tree, each branch has a unique upstream node, so that we can conveniently indicate the branch upstream of node *i* also as branch *i* with branch length *t*_*i*_.

The leaf-nodes of the tree correspond to the data points that were provided to the algorithm. Bonsai assumes that for each leaf-node *i*, the input data provides an estimated position ***µ***_*i*_, i.e. a vector in a continuous *d*-dimensional ‘feature space’ with a vector of error-bars ***σ***_*i*_ that correspond to the standard-deviations of the ‘noise’ in the measurement of each feature of node *i*. Although Bonsai can take any input data of this type, in the following we will use notation and terminology as if the data points are cells, and the estimated ‘features’ *µ*_*gi*_ correspond to the log-expression of gene *g* in cell *i*, and the error-bars *σ*_*gi*_ are the corresponding measurement noise levels. We generally denote the (unknown) ‘true’ positions of each node *i* as vectors ***x***_*i*_ in the *d*-dimensional feature space (e.g gene expression space). Lastly, we will assume that it is clearly defined which observed cell data is supposed to come from which node on the tree and use the same index *i* for the node corresponding to cell *i*.

### SI.B.1 The likelihood of a tree with given branch lengths

The likelihood of the data *D* given a certain tree *T* with branch lengths ***t*** is the product of two types of terms:

1. The likelihood of observing the mRNA-counts for cell *i* given a true position of that cell in gene expression space. We’ll denote this term by *P* (cell_*i*_|***x***_*i*_).
2. The likelihood that node *i* has moved through gene expression space from the position of its ancestor in the tree *π*(*i*). This term is denoted by *P* (***x***_*i*_|***x***_*π*(*i*)_, *t*_*i*_).

Compiling these terms, we get the likelihood of the data *and* all tree node coordinates conditioned on the tree topology and the branch lengths:

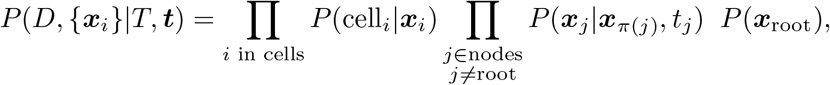

where the *P* (***x***_root_) is the prior probability for the position of the root. We will take a uniform prior for the position of the root, so that this term will drop out of the likelihood. Based on this, we can get the likelihood of the data conditioned on the tree topology and the branch lengths by marginalizing over the positions of all tree nodes:

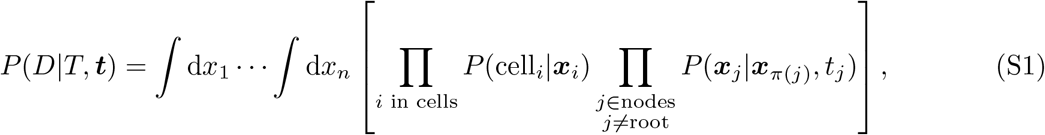

where *n* is the number of nodes in the tree. In the following subsections, we will provide more detail on the individual likelihood terms and solve the integrals.

#### SI.B.1.1 Likelihood of the data at the leaf-nodes given a true position

One part of the likelihood corresponds to factors *P* (cell_*i*_|***x***_*i*_) for the data of object *i* given an assumed position ***x***_*i*_ in the feature space. Bonsai assumes that for each object *i*, the input specifies a vector of estimated position ***µ***_*i*_ with error-bars ***σ***_*i*_, and that this factor in the likelihood is given by a product of Gaussian distributions:

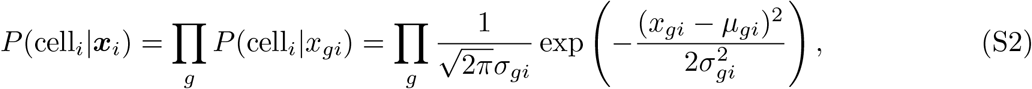

That is, we require that the input data is normalized such that the likelihood of the ‘measured’ features ***µ***_*i*_ is reasonably approximated by a multi-variate Gaussian with standard-deviations ***σ***_*i*_ and negligible covariances. As we discuss for scRNA-seq data below, this might require a careful definition of the variables to be inferred, which will depend on the data type. Apart from the specific application for scRNA-seq, it will be up to the user to provide appropriately normalized data to *Bonsai*.

#### SI.B.1.2 The likelihood of gene expression data given a cell’s true gene expression state

In scRNA-seq data, the experimental output after preprocessing is a so-called count table, which is a matrix of mRNA-counts for each gene (as rows) and for each cell (as columns). However, to normalize this data into estimated positions in gene expression space *µ*_*gi*_ with error-bars *σ*_*gi*_ we need to carefully take both technical and biological noise into account [76–79]. For this, we will use *Sanity* [28], a Bayesian method that provides the best guess for a cell’s position in gene expression space *and* uncertainty estimates on these guesses. We refer to the *Sanity* -publication for a detailed description of this method, but we summarize the key points here.

As described in the *Sanity* publication [28], the mRNA counts in a cell are the result of the processes of transcription and mRNA decay, which are intrinsically stochastic due to the unavoidable thermal noise at the molecular level. Consequently, rather than controlling precisely when how many mRNAs of which genes are made or degraded, the physical state of the cell only determines the *rates* at which each gene is transcribed and its mRNAs are degraded. As the physical state of the cell changes in time, these transcription and degradation rates will vary as well. Consequently, the number of mRNAs *m*_*gc*_ for gene *g* in a given cell *c* is a *stochastic* function of the transcription rate *λ*_*gc*_(*t*) and mRNA degradation rate *µ*_*gc*_(*t*) of this gene in the recent past. In particular, the expected number of mRNAs is given by

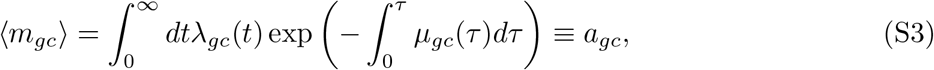

with *λ*_*gc*_(*t*) and *µ*_*gc*_(*t*) the transcription and mRNA decay rates a time *t* in the past of this cell. We denote this expected mRNA counts as the *transcriptional activity a*_*gc*_ of gene *g* in cell *c*. The *actual* number of mRNAs in the cell will be a sample from a Poisson distribution with mean *a*_*gc*_.

To a reasonable approximation, the number of captured mRNA-molecules, *n*_*gc*_, in a scRNA-seq experiment correspond to a random sample of the true mRNA molecule number in the cell *c* [28, 76, 80]. Consequently, the combined effects of the intrinsic biological and technical noise are that these counts *n*_*gc*_ are still Poisson samples of the transcriptional activities *a*_*gc*_ [28].

Because the efficiency of capture varies from cell to cell, and because it can be argued that two cells with proportional transcription activities 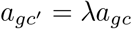 are effectively in the same gene expression state (i.e. they only differ in the absolute size of their mRNA pool), it is more meaningful to describe the gene expression state by *transcription quotients* 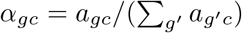. Finally, because absolute gene expression levels vary over 4 − 5 orders of magnitude between genes, it is generally more meaningful to express gene expression levels in logarithmic scale (so that the same fold-change in expression corresponds to the same change, irrespective of the absolute expression level). Therefore, *Sanity* expresses the gene expression state of the cell in terms of *Log Transcription Quotients* or LTQs:

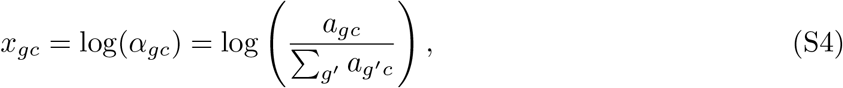

where the *a*_*gc*_ are defined in terms of transcription and mRNA decay rates according to equation (S3).

*Sanity* uses a Bayesian algorithm to estimate the LTQs, *x*_*gc*_, for each gene *g* in each cell *c*. In its output, it reports the means 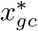 and standard-deviations *ϵ*_*gc*_ of the *posterior* distributions of the LTQs for each gene in each cell. In addition, it also reports an estimate *v*_*g*_ of the true variance in LTQs of gene *g* across the cells in the dataset. Notably, because of the Poisson sampling noise, and because absolute expression levels vary over several orders of magnitude, the resulting sizes of the error-bars *ϵ*_*gc*_ vary enormously across genes and even across cells for the same gene. This variation in measurement accuracy is crucial to take into account in the analysis of scRNA-seq data.

Finally, while *Sanity* provides posterior estimates, *Bonsai* requires as input estimated positions and error-bars of the *likelihood* instead of the posterior. Fortunately, the posterior is proportional to the product of the desired likelihood term and *Sanity*’s prior, such that the likelihood can be well-approximated from the posterior. We will therefore follow the derivation given in [28] and obtain the measurement estimates *µ*_*gc*_ and error-bars *σ*_*gc*_ from the reported posterior estimates 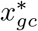 and posterior error-bars *ϵ*_*gc*_ using the identities:

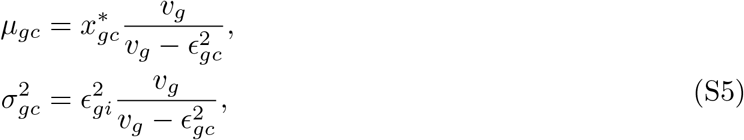

where *v*_*g*_ is *Sanity*’s estimate of the variance for gene *g*.

##### Note on computational implementation

Practically, we can thus use the Equations (S5) to transform the output from Sanity, and then the likelihood term *P* (cell_*i*_|***x***_*gi*_) takes the Gaussian shape in *µ*_*gi*_ and *σ*_*gi*_ as in Equation (S2).

Additionally, to be able to reconstruct this likelihood best, we made a slightly altered version of the Sanity-implementation. Normally, Sanity returns values that are obtained by integrating over the possible values of *v*_*g*_, so that the returned *v*_*g*_ is the expectation value of *v*_*g*_, rather than the maximum posterior estimate. Although this is the most correct option for obtaining posterior estimates of the LTQs, the transformation to reconstruct the likelihood can become inaccurate in some cases. Therefore, we now adapted Sanity to have a mode in which it returns the values for the *v*_*g*_ that maximize the posterior.

#### SI.B.1.3 Selecting features with signal-to-noise over a threshold

In scRNA-seq, many genes with relatively low expression are sampled only very sparsely, so that the error-bars on their expression levels may be much larger than the true variation in their expression levels across cells. Such genes contain very little information about the similarities of gene expression across cells (and thus the tree structure) and we have found that including them leads to worse performance of *Bonsai* on simulated data. That is, they appear to add more noise than they add information about the structure in the data. We therefore decided to only include genes (or more generally, features) that have signal-to-noise over a threshold.

Let 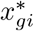 denote the posterior estimate of the LTQ of gene *g* in cell *i, x*_*g*_ the estimate of the mean LTQ over the cells, and 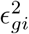 the error-bar on the deviation 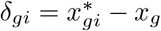. We then define the signal-to-noise ratio, *S*_*g*_, as the average ratio of the posterior estimate of the squared deviation of the’signal’ and the posterior error-bars 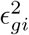 on this signal, i.e.

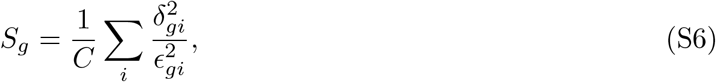

with *C* the number of cells. For scRNA-seq data, *Sanity* reports the means and variances of the posteriors on the gene expression values (see Section SI.B.1.2), so that we can use this output directly to estimate *S*_*g*_.

For other data, we estimate *S*_*g*_ from the input expression estimates *µ*_*gi*_ with error-bars *σ*_*gi*_ as follows. If we assume the true expression values *x*_*gi*_ are drawn from a Gaussian with variance *v*_*g*_ and mean *µ*_*g*_ (which are both unknown), then the likelihood of the estimates *µ*_*gi*_ is given by

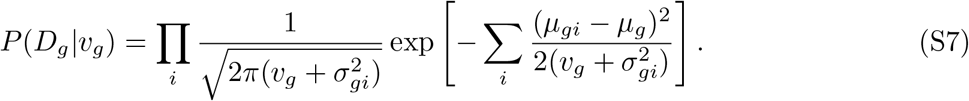

We now first estimate the mean, *µ*_*g*_, by its maximum likelihood value, 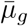, which gives

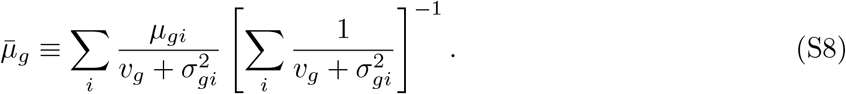

Note that this maximum likelihood estimate, 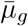, depends on the variance *v*_*g*_ that we still need to be estimate. We therefore substitute the true mean, *µ*_*g*_, by the estimate, 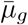, in the likelihood given by Equation (S7), and next estimate the variance *v*_*g*_ by maximizing this likelihood, *P* (*D*_*g*_|*v*_*g*_). The maximal likelihood value of *v*_*g*_ is a solution of the equation

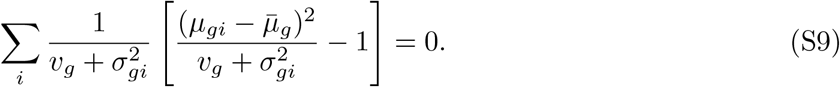

Given the estimated value of the variance *v*_*g*_, the posterior estimate of the’signal’ (*x*_*gi*_ − *x*_*g*_) in cell *i* is given by

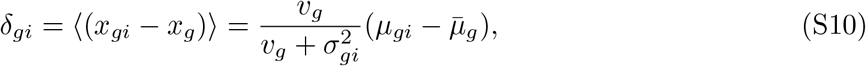

and the error-bar on this estimate is

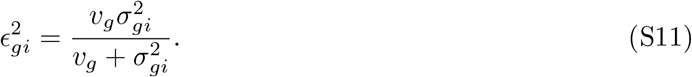

We then finally estimate the signal-to-noise *S*_*g*_ using equation (S6).

By default we select all genes with *S*_*g*_ ≥ 1 to include in the *Bonsai* analysis. However, if desired, users can change the threshold value on *S*_*g*_.

#### SI.B.1.4 The prior probability of movement in gene expression state

The second term in the tree likelihood (Equation (S1) is the prior probability of a movement in gene expression space from a position ***x***_*π*(*i*)_ to the position ***x***_*i*_. This term captures how likely the inferred gene expression changes along the branches of the tree are.

It is important to note that *P* (***x***_*i*_ |***x***_*π*(*i*)_) captures only the *prior* probability distribution on gene expression changes, while by reconstructing the tree on the data we try to infer the *posterior* distribution of gene expression changes. Since we do not want to impose strong assumptions on which gene expression changes are more likely than others, we want to take a very conservative prior. Therefore, we only assume that 1) gene expression changes can be described by a continuous Markov process, and 2) gene expression changes are *a priori* equally likely in all genes relative to the total variation in that gene in the dataset.

The assumption that we can describe gene expression changes by a continuous Markov process means that we expect no discontinuous jumps in gene expression, and that we expect the current gene expression state to be predictive of changes in gene expression. ^2^ In [31, 32], it is shown that, under a mild additional assumption^3^, any continuous Markov process must be a drift-diffusion process, which can be described by the form

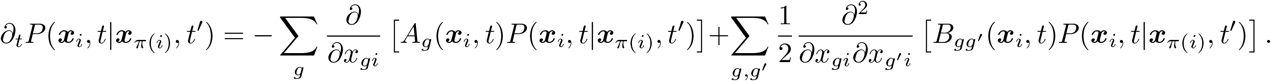

This equation describes the time evolution of the probability distribution of a cell to be at gene expression state ***x***_*i*_ at time *t* while it was at ***x***_*π*(*i*)_ at timepoint *t*^′^. The term *A*_*g*_ describes a deterministic drift that drives the cell’s expression and the term 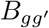describes stochastic diffusion. In our prior process, we will set *A*_*g*_ to zero so as not to assume any deterministic drift; rather, we aim to learn gene expression changes from the data instead of imposing uncertain prior knowledge. Similarly, for the diffusion term 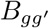, we do not want to assume prior knowledge about any correlation between the genes, so that we will assume independent diffusion across the genes: 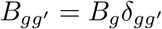. Finally, we set *B*_*g*_ such that the amount of expected diffusion in a gene is proportional to the total variation that is observed in the dataset: *B*_*g*_ = *v*_*g*_. We thus get

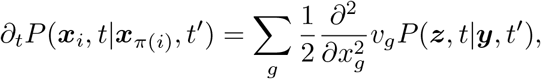

of which the solution is given by a Gaussian

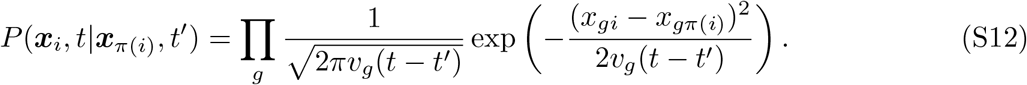

This gives us the likelihood term for movement in gene expression space where the diffusion time along the branch *t*_*i*_ := *t* − *t*^′^.

#### SI.B.1.5 The full tree likelihood expression

Taking together Equations (S1), (S2), and (S12), we get an expression for the likelihood of the tree:

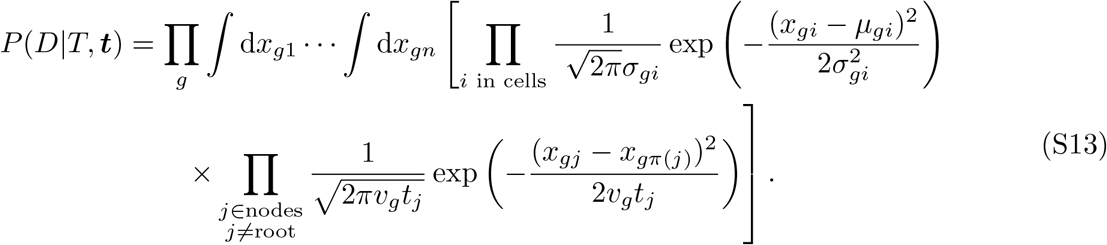

It is important to note here that the set of cells is a subset of the set of nodes. Every cell *i* has an associated node with coordinates *x*_*gi*_ that will show up in the first term, but these will also show up in the second term.

##### Practical note: the likelihood is independent on the choice of root

Because we took a uniform prior over the root position, and because the expression for the likelihood of movement is symmetric in ***x***_*j*_ and ***x***_*π*(*j*)_, we have the following identity

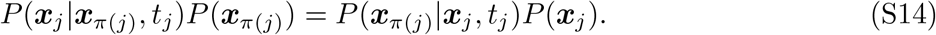

As a result, we can always designate another node as the root without affecting the tree likelihood. We will use this freedom to choose the root node in several derivations and computational implementations.

#### SI.B.1.6 The tree likelihood for other types of data

Although we have focused the discussion above on single-cell RNA-sequencing data, the equations for the tree likelihood generalize to other types of data. Importantly, the assumptions about the data that we have made above should hold, in particular

- We should be able to ‘normalize’ the data such that each cell or sample can be described by coordinates *x*_*gi*_, and the likelihood of the data can be well-described by a multivariate Gaussian distribution in these coordinates with negligible covariance (analogous to Eq. (S2)).
- The prior likelihood of moving from ***x***_*π*(*j*)_ to ***x***_*j*_ in these coordinates can be described by a Markov process in which we do not expect discontinuous jumps.

Once these assumptions are met, the rest of the derivations presented here hold for any data type. *To facilitate a clean presentation, we will in the following borrow scRNA-seq terminology (e*.*g*., *gene, cell, LTQ), and we leave it to the reader to translate these terms to analogs for their preferred data types*.

### SI.B.2 Analytical solutions for the tree likelihood

In this section, we will analytically solve the integrals from Equation (S13). To facilitate this derivation we will introduce several variable transformations, and I will give an overview of those here.

Definition of variables used throughout this SIthroughout this SI

**Tree variables**

- *t*_*i*_: the branch length (or diffusion time) of the branch upstream of node *i*.
- *v*_*g*_: the estimated total variance for gene *g* across the dataset
- *x*_*gk*_: the true LTQ of gene *g* in node *k*. If the node-label *k* matches a cell-label *i*, this cell is associated with that node.

**Observed cell variables**

- *µ*_*gi*_: the mean of the Gaussian distribution describing the likelihood of the experimental data given the log-transcription quotient (LTQ) of gene *g* in cell *i*. This mean is only defined for cell-associated nodes.
- 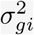: the corresponding variance of the likelihood distribution.
- 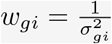: the *precision* for gene *g* at cell *i*, defined as the inverse of the variance.
- 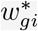: the *precision* for gene *g* at node *i* when corrected for the uncertainty induced by diffusing for a time *t*_*i*_.

**Effective leaf variables**

- 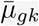: the mean of the Gaussian distribution that describes the likelihood for the entire subtree downstream of node *k*. This mean is thus also defined for nodes not associated with a cell. *Note that these effective leaf variables depend on the definition of the root since this determines which nodes are downstream. If this is not clear from the context this will be indicated by* 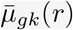, *where r is the designated root*.
- 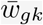: the corresponding precision for the effective leaf. The combination 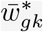 indicates the precision when accounted for the uncertainty induced by diffusion.
- *C*_*r*_(*a*): the set of leaves that are directly downstream of node *a*, conditioned on *r* being designated as the root.

#### SI.B.2.1 Solving the integral for the star-tree

We will first solve the integrals in Equation (S13) in the simplified case of a *star-tree*, which is the simple tree where all cells have an associated node and these ‘cell-nodes’ are all connected to a single root. Denoting the number of cells by |*C*|, the star-tree thus has |*C*| + 1 nodes and |*C*| edges. Later, we will show that we can use the star-tree result to solve the integrals for any tree. Denoting the coordinates of the root-node by *x*_*gr*_, the equations for the star-tree likelihood become

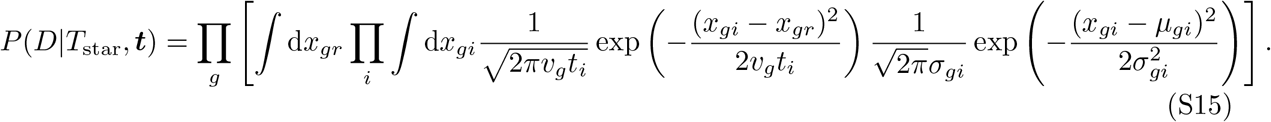

The integral over *x*_*gi*_ can be done by completing the squares in the exponent and recognizing the standard Gaussian shape. This gives

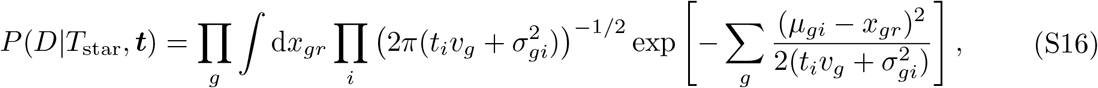

It is useful to here introduce the ‘precision’ of a node, 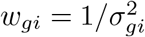, which is defined as the inverse of the variance in the posterior for the node position. We also define 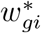 as the precision corrected for the uncertainty induced by diffusion over a time 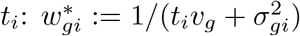. We can then write the likelihood of the star-tree in terms of the distances from all the leaves to a center of mass 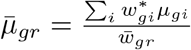 where 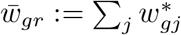. We will derive this per gene first, and only afterwards take the product. Therefore, we drop the subscripts *g* for now:

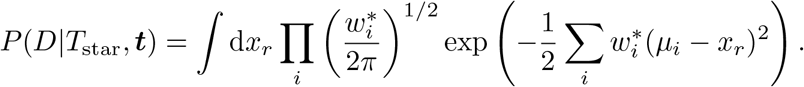

Completing the squares in the exponent and re-writing gives

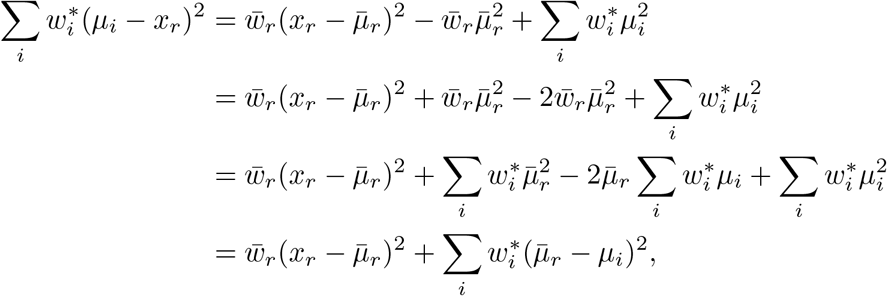

and inserting this gives

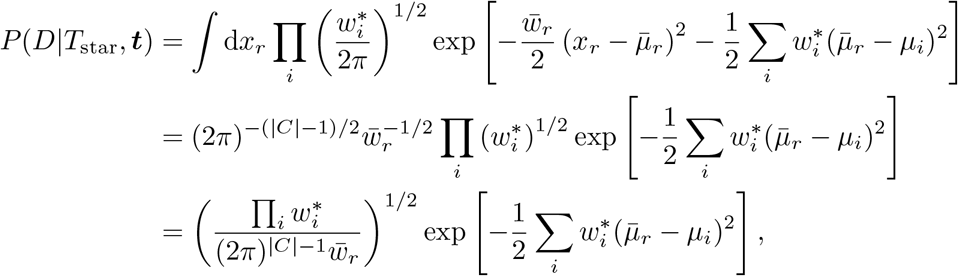

For all genes together this becomes

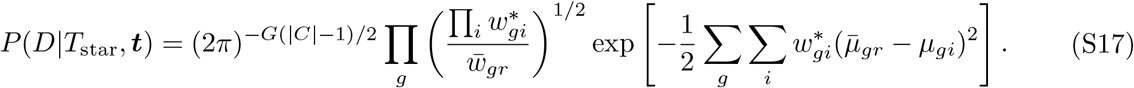

Note that the dependence on the branch lengths *t*_*i*_ enters through:

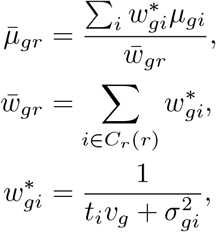

where we use the notation *C*_*r*_(*r*) to indicate the nodes directly downstream of *r* when the root is *r*.

#### SI.B.2.2 Summarizing a sub-tree in an *effective leaf*

We will here show that the contribution to the tree likelihood of a subtree consisting of one ancestor (*t*_*a*_) with several leaves as children, can be summarized by replacing this sub-tree by a single *effective leaf*. This can be used recursively to reduce any tree to a simple star-tree consisting of a root connected to effective leaves that summarize the different sub-trees. This provides an efficient way of calculating the tree likelihood.

For deriving this, we will focus on the set of leaves downstream of an ancestor that we will label *a*. Wihtout loss of generality, we can designate the node upstream of *a* as the root (*r*), because the tree likelihood is independent of the choice of root. In the following, we will denote by *C*_*r*_(*a*) the set of children of ancestor *a* when *r* is considered the root. The sub-tree downstream of *a* we will denote by *T*_*ds*_, and *R* will be the set of all nodes other than *r, a*, and the nodes in *T*_*ds*_. The likelihood contribution of the nodes in *R* only depends on the position of the root *r*, such that we can gather this contribution in a term *P* ({*z* ∈ *R*}|*x*_*gr*_). We get.

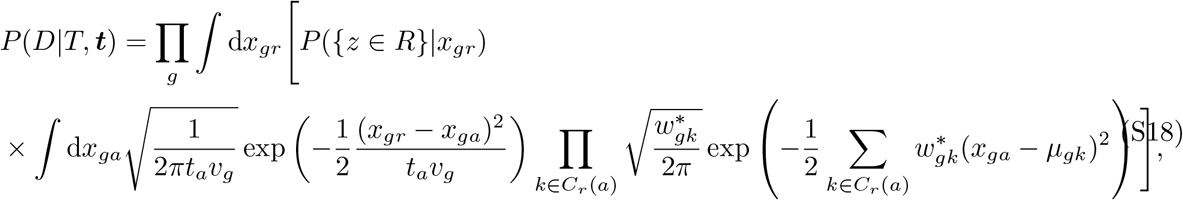

where the integral over the leaf positions *x*_*gk*_ was already performed. We will again do the derivation for one gene at a time, so that we can drop the corresponding subscripts. As before, we can complete the squares in the second exponent. We will denote the effective precision of the effective leaf that summarizes the subtree downstream of *a* as 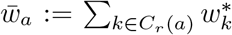, and the effective mean as 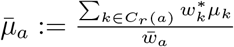, we get

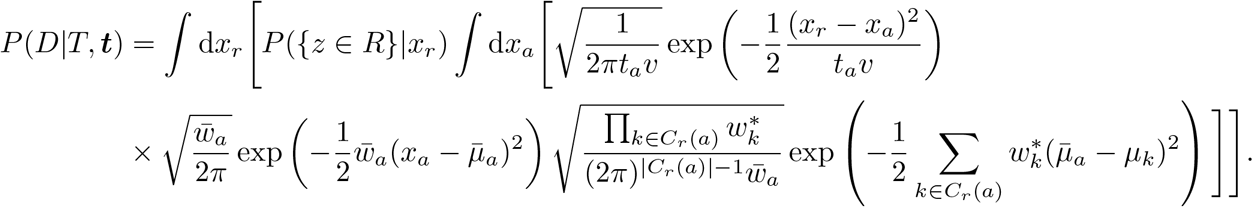

Performing the integral over *x*_*a*_, we get

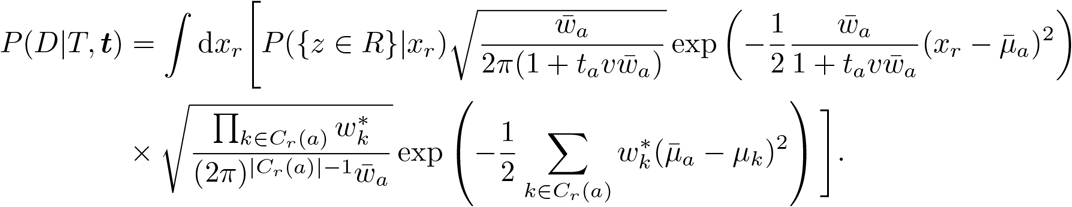

We can see that the second line in this expression only depends on the sub-tree below *a* and not on the root position *x*_*r*_. In fact, it is exactly what one would get for the likelihood of a star-tree built from the leaves in *C*_*r*_(*a*). We can expose this by writing the likelihood in a form that shows that the subtree downstream of *a* is now just summarized by an effective leaf with position 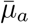 and an effective precision 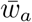. Here, we recognize that this effective precision becomes 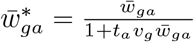 when accounting for the uncertainty induced by diffusion to node *a*. For easy reference, we here also write out all the definitions we used and re-introduce the product over genes:

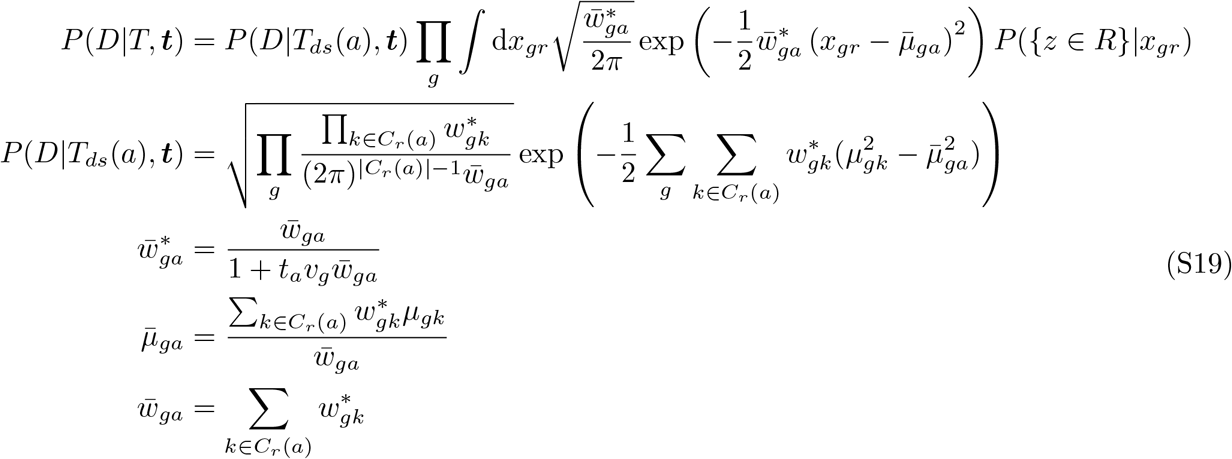

In these expressions, we have denoted by *P* (*D*|*T*_*ds*_(*a*), ***t***) the likelihood that we would get if the sub-tree downstream of *a* would be the full tree.

#### SI.B.2.3 The loglikelihood of a general tree

Equation (S19) shows that we can write the likelihood of a general tree in terms of the likelihood of smaller sub-trees. This means we can now perform all the integrals for a general tree recursively. It is useful to here collect the general expression for the resulting loglikelihood:

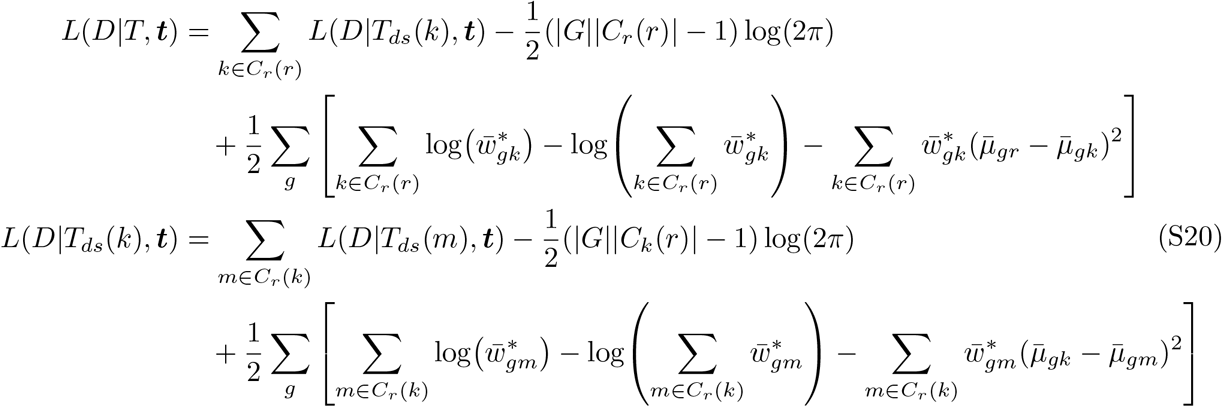

In these equations we have written 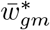, but it should be understood that if *m* is a cell-associated node, this should be replaced by 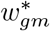.

##### Note on the factors of 2*π*

By doing the accounting correctly, one can see that the number of factors of 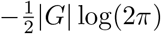 will always be exactly the total number of observed cells minus one. This is thus independent of the tree topology, and we can (and will) therefore leave out these factors in the following derivations and in the computational implementation.

##### Note on the influence of the gene variances *v*_*g*_

Recall that the diffusion rate for different genes in the diffusion prior was scaled to the observed variance in that gene in the dataset, i.e. the variance in the Gaussian prior for movement in gene expression space was *t*_*i*_*v*_*g*_. We can now see how these variances influence the loglikelihood. The *v*_*g*_ enter the expression through the effective precisions 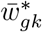 and 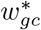. These effective precisions take two different forms:

- for nodes associated with an observed cell we get: 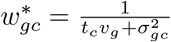
- for internal nodes we get: 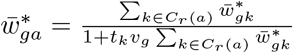, where 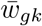 are the effective precisions of the child-nodes.

It is now useful to perform the following transformation of the means and variances for the observed cells, i.e. of the raw input that *Bonsai* uses:

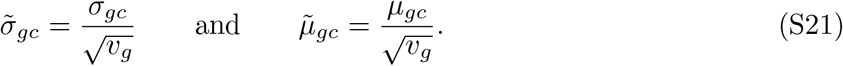

As a consequence, we get ^4^

- for cell-associated nodes: 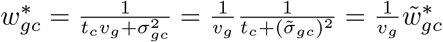
- for internal nodes: 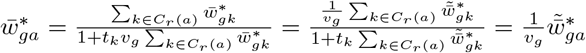

As a consequence, for the loglikelihood we get

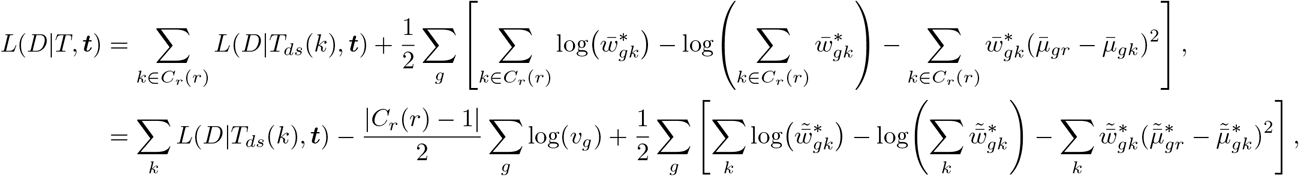

and similarly for the sub-tree likelihoods *L*(*D T*_*ds*_(*k*), ***t***). Just as for the factors of 2*π* we see that the variances will add a term to the loglikelihood that is independent of the tree topology (only scaling with the number of cells). We can thus, without loss of generality, perform this variable transformation and after that discard the *v*_*g*_-terms, which is what we will use in the following.

##### Choosing a new root

We have pointed out before that the tree likelihood is independent of the choice of root. This means that the recursive expression for the loglikelihood (Eq. (S20)) can be written around any internal node. For this, it is important to realize that the definitions of the effective mean and precision of node 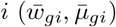 depend on which set of nodes is downstream of this node, which depends on the choice of root. When choosing a new root, we can use that the new means and precisions can be efficiently computed from the old means and precisions.

### SI.B.3 Optimization of the branch lengths

To learn the diffusion times, or branch lengths, that maximize the likelihood of the data, we will optimize the branch lengths, ***t***. For this optimization, it works best to use a quasi-Newton method that uses both the loglikelihood and its derivatives with respect to all branch lengths. For this, it is essential to have a fast way to calculate this loglikelihood and its gradient.

In section SI.B.2.2 we derived that we can summarize the likelihood contribution of any sub-tree by replacing it by an effective leaf. We will now use this to calculate the derivative of the loglikelihood with respect to one of the branch lengths, *t*_*k*_. We start by picking the corresponding edge (let’s say it connects nodes *l* and *k*) and we summarize the sub-trees downstream of *l* as seen from node *k*, and vice versa. Here, we can denote the nodes neighboring *k* without node *l* as *C*_*l*_(*k*), and the converse by *C*_*k*_(*l*). The likelihood (again for one gene) becomes:

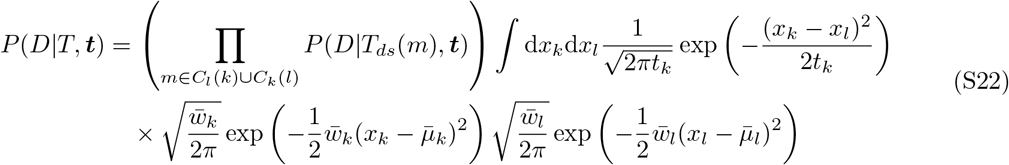

Then performing the integrals, we get

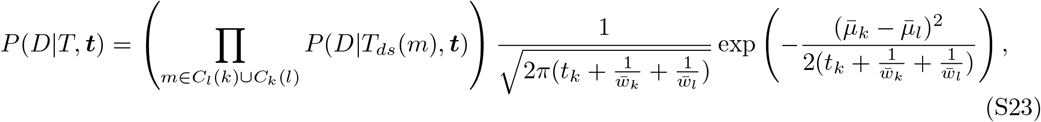

resulting in an alternative expression for the loglikelihood (up to factors of 2*π*):

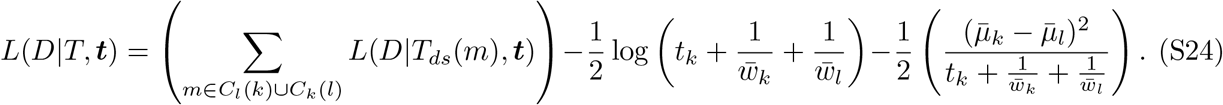

In this form, we can easily find the derivative w.r.t. *t*_*k*_:

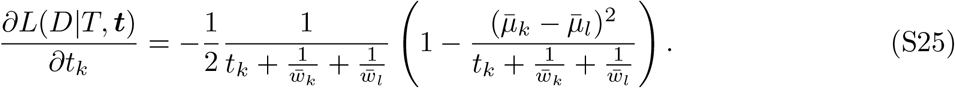

This can be done for all branch lengths in the tree to get the full derivative.

## SI.C Finding the tree with maximum posterior

In Section SI.B, we have derived an expression for the likelihood of the data *given a single tree*. However, we are interested in finding the tree with maximal posterior probability. For this, we should first choose a prior distribution over tree topologies and branch lengths, which we will just take to be uniform. Therefore, we get

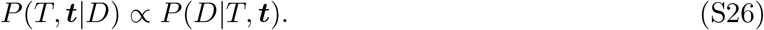

Finding the most likely tree thus boils down to finding a tree that maximizes the likelihood of the data. We have already mentioned that we can maximize the branch lengths for a given topology using the gradient derived above, so the problem remains to find the best tree topology. However, the number of possible tree topologies is extremely large, and it is computationally infeasible to do a brute-force search for the maximal tree. Therefore, we will use the following scheme which is based on tree space search strategies learned from phylogenetics.

1. Start from a star-tree
2. Iteratively add the most likely internal node
3. Further resolve polytomies
4. Optimize branch times
5. Interchange nearest neighbors
6. Optimize branch times a final time

On the following pages, I will explain the separate steps.

### SI.C.1 Start from a star-tree

To start, we create a leaf-node for each cell, and connect these leaves to a single root-node. We thus have a tree with |*C*| + 1 nodes, and |*C*| edges. For this tree, we can calculate the loglikelihood and its gradient with respect to the branch lengths, which we use to do a first optimization of these branch lengths.

### SI.C.2 Iteratively add the most likely internal node

Starting from the star-tree, we use a tree space search strategy loosely based on the popular ‘Neighbor-Joining’ heuristic [34]. For a pair of leaves in the tree, we can propose to add an ancestor between the root and the leaves (see Figure S20**B)**. The new tree would then have |*C*| − 2 leaves connected to the root (*r*), while the other 2 leaves are connected to an ancestor *a*, which is in turn connected to the root. We here search for *the pair of leaves for which adding an ancestor increases the likelihood most*. We will describe below (Section SI.C.2.1) how to calculate this increase in tree likelihood.

After this ancestor is added, we can summarize the newly created subtree in an effective leaf for the ancestor *a*, as described in Section SI.B.2.2 (see Figure S20**C)**). In this way, we have created a star-tree again and we can again search for the best pair of (effective) leaves to merge into a new ancestor. This is repeated until either the root has only 3 children left, or when no pair can be found that increases the tree likelihood.

#### SI.C.2.1 Finding the ancestor that increases the likelihood most

We want to calculate the likelihood increase of merging a pair of leaves *k, l* into an internal node *a* that connects to the root *r*. We thus want to calculate the likelihood for the tree *T*_merge_ where this ancestor is added, and compare it to the likelihood of the tree *T*_orig_ without the added ancestor. In the following, we will again use that we can summarize subtrees in effective leaves (Section SI.B.2.2), and we will denote all leaves other than *k, l* as a set *R*.

##### The likelihood for *T*_orig_

It is convenient to summarize the subtree (*T*_*R*_) comprising the root and all leaves other than *k, l* into an effective leaf. We get a star-tree with three leaves:

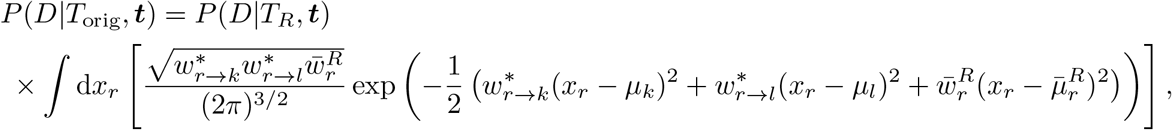

where 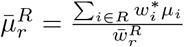 and 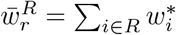. We also need to be explicit about the time-dependence of the precisions, we have 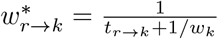, and similar for 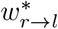. We have here written *t*_*r*→*k*_ to distinguish it from the branch length *t*_*a*→*k*_ that we will introduce later. Using the expression for the star-tree loglikelihood we get

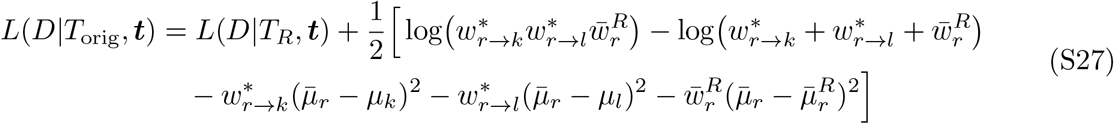

##### The likeilhood for *T*_merge_

When we do merge the leaves *k, l* into an ancestor *a*, it is best to designate the new ancestor as the new root. The likelihood then takes a very similar shape, although the term 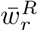 is replaced by 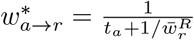 to account for the added diffusion time from the root to the ancestor:

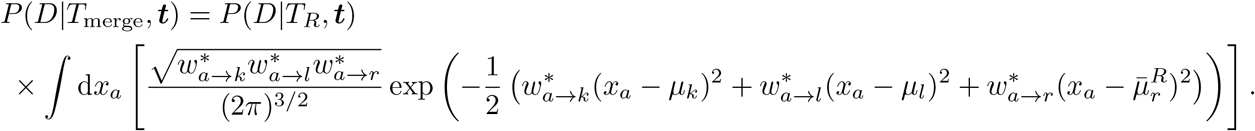

Performing the integral gives

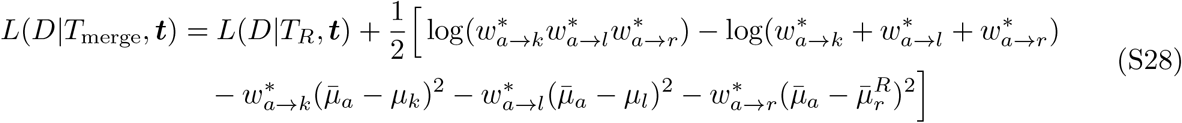

##### The increase in loglikelihood

Combining Equations (S27) and (S28), we can calculate the increase in loglikelihood caused by merging leaves *k, l* into an ancestor:

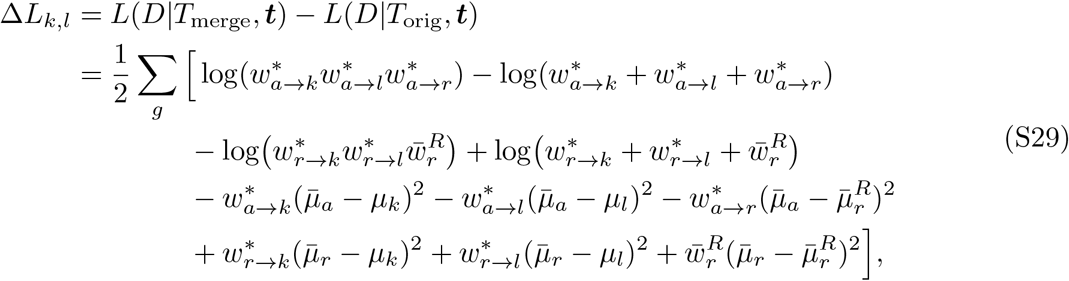

where we thus have the following differences between the two tree topologies:

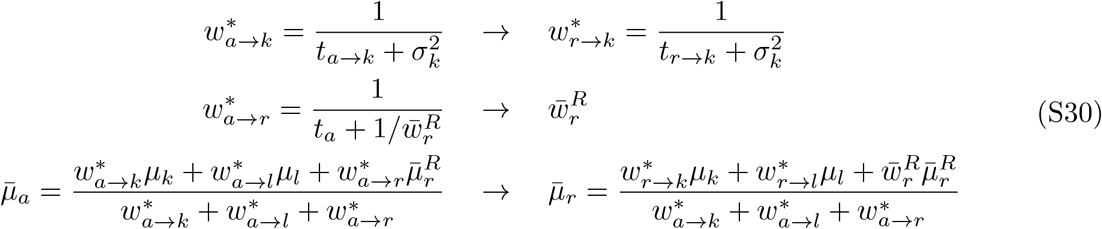

##### Optimizing the branch lengths after the merge

Adding an internal node to the tree could in principle change the optimal branch lengths across the whole tree. However, performing a tree-wide branch length optimization after every proposed merge would quickly become computationally infeasible. Therefore, we assume that the optimal branch lengths are well-approximated by only optimizing the branch lengths in the subtree that is changed, i.e., we only optimize *t*_*a*→*k*_, *t*_*a*→*l*_, *t*_*a*→*r*_ for calculating Δ*L*_*k*,*l*_.

In addition, since we will iteratively add more and more ancestors to the tree, there is a large chance that the branch length *t*_*a*→*r*_ will have to be re-optimized later, either because another ancestor (*a*^′^) is added upstream of *a* so that we optimize a branch length 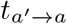, or because *t*_*a*→*r*_ simply becomes sub-optimal because the root changes position when merges are done in different parts of the tree. The same does not hold for the connection between the merged leaves *k* and *l*. As a result, it turns out that our tree-reconstruction yields the highest likelihood trees if we do the following time-optimization scheme:

1. First we optimize the sum *t*_*a*→*k*_ + *t*_*a*→*l*_ by assuming that the ancestor *a* is not attached to the root. This will ensure that the sum of these times (which will thus not change in the following steps in the iteration) is optimized to capture the distance between the leaves *k* and *l*.
2. Then optimize all three times *t*_*a*→*k*_, *t*_*a*→*l*_, *t*_*a*→*r*_ while constraining the above sum.

### SI.C.3 Further resolve polytomies

In the previous step we have iteratively merged pairs of child-nodes of the root into new ancestors, until no pair could further increase the tree likelihood. It turns out, however, that this procedure can result in downstream nodes that have more than 2 child-nodes, so-called polytomies. This for example happens when first an ancestor *a* is added upstream of leaves *k* and *l*, and then another ancestor (*a*^′^) is added upstream of *a* and a leaf *m*, but the optimal times are such that 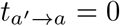. Of course, an edge with zero length doesn’t change the likelihood of the tree, so we can remove the node *a* and directly connect *a*^′^ to *k, l, m*. Since this tree results from an optimization, this configuration is indeed the best given the root-position at that point in the optimization. However, later in the iterative optimization, the root-position may have changed enough to warrant reconsideration of the topology *a* → *k, l, m*. This turns out to be often the case. Therefore, we add a step in the procedure where we go over all nodes that still have more than 2 child-nodes, and we greedily add the ancestors downstream of these nodes that are mostly increasing the likelihood. Note that the necessary calculations are exactly the same as in the original merging procedure (Section SI.C.2), except that now we don’t only consider the children of the root for merging, but also the children in other polytomies.

### SI.C.4 Optimize branch lengths

Since in the iterative procedures above, we only maximized the branch lengths directly connected to the new ancestor, we may have drifted away from the global optimum for all branch lengths. Therefore, we do another round of gradient-based optimization of all branch lengths (see Section SI.B.3).

### SI.C.5 Interchange nearest neighbors

In phylogenetic tree reconstruction, tree moves are often used to search the tree space. These moves change the tree topology after which a new tree likelihood is calculated. A well-known tree move is a so-called Nearest Neighbor Interchange (NNI). In this move, we pick an arbitrary edge in the tree that is not connected to a leaf, let’s say the edge connects nodes *k* and *l*. We then consider the sub-trees connected to the nodes *k* and *l*. In a fully binary tree, i.e., a tree without any polytomies, we would always have 4 sub-trees. Now an NNI-move consists of reconnecting these 4 sub-trees in different ways. For example, if subtrees *T*_*A*_, *T*_*B*_ were downstream of *k* and *T*_*C*_, *T*_*D*_ downstream of *l*, an NNI-move could connect *T*_*A*_, *T*_*C*_ to one node and *T*_*B*_, *T*_*D*_ to another. The nearest-neighbors of these sub-trees have thus been interchanged.

The trees that are reconstructed by *Bonsai* often have polytomies, so we need a way to generalize this NNI-move to having more than 4 sub-trees. We chose to do this by doing the following:

1. We pick an arbitrary edge that connects two internal nodes, say *k* and *l*.
2. We remove the edge and node *k* by taking all sub-trees that were attached to node *k* and attach them to node *l*.
3. We now consider the node *l* as the root of a star-tree. We can now apply the same procedure of iteratively adding ancestors as described in SI.C.2.

It can be seen that this is a generalization of the NNI-move in the case of 4 sub-trees. In that case, we can maximally add one ancestor, and all possibilities are to group the subtrees as (*T*_*A*_, *T*_*B*_), (*T*_*C*_, *T*_*D*_), (*T*_*A*_, *T*_*C*_), (*T*_*B*_, *T*_*D*_) or rather (*T*_*A*_, *T*_*D*_), (*T*_*B*_, *T*_*C*_).

#### Randomly interchanging nearest neighbors

To make sure we are not getting stuck in a local optimum, we do several rounds of random NNI-moves. We follow the steps as outlined above, but now we randomly pick a pair of child-nodes for which we add an ancestor. This random choice is weighted by the tree likelihoods, such that the probability of merging *q, r* is 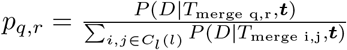.

#### Greedily interchanging nearest neighbors

We end by performing NNI-moves in a greedy fashion. That is, we determine the loglikelihood increase that an NNI-move would bring for every edge in the tree and perform the NNI-move that maximizes this increase. This is repeated until no NNI-move significantly increases the likelihood.

### SI.C.6 Optimize branch lengths a final time

Again, in the NNI-moves, only the connected branch lengths are optimized. We therefore do another round of branch length optimization to end with the tree that is most likely to describe the data.

## SI.D Speeding up the tree reconstruction

Current single-cell datasets can have over 10 million cells, a number that can even complicate running a method that scales quadratically in this number. The above described *Bonsai* implementation needs to compare the loglikelihood increase for merging all pairs of cells at every addition of a new internal node. Since the number of internal nodes that will be added is also proportional to |*C*|, and because the calculation of one loglikelihood increase scales with the number of genes |*G*|, the computational scaling is *O*(|*C*| ^3^ |*G*|). This is clearly problematic. In this Section we show two computational tricks that speed *Bonsai* up to a manageable scaling.

### SI.D.1 Calculating upper bounds for merging likelihoods given a change in the root’s position

In the tree reconstruction procedure, we repeatedly have to calculate the likelihood increase upon merging pairs of leaves into an ancestor. In every iteration, we calculate the likelihood increase for merging many possible pairs, but only one of these pairs leads to the largest likelihood increase, say pair *m, n*. We will thus only merge the maximal pair *m, n* into a new ancestor. Because this merge will influence the position of the root, Equation (S29) shows that this will also affect the loglikelihood increase due to merging all other pairs of leaves. In principle, all calculations should thus be repeated.

However, Equation (S29) also shows that adding an ancestor somewhere else will *only* affect the likelihood increase for pair (*k, l*) through the effective position and precision of the root when accounting for all other leaves, i.e. through 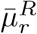, and 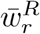. In turn, these depend on the effective position and precision when accounting for all leaves via 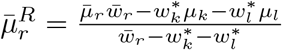 and 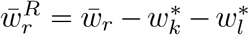. We can therefore try to estimate an upper bound on what Δ*L*_*k*,*l*_ can maximally become given a certain amount of movement in the root position. For this, we will first compute a linear approximation of Δ*L*_*k*,*l*_ as a function of 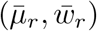. Then, we will take an ellipsoid-region around the current 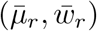 and calculate the maximum for the linear approximation in this region. This, we will use as an upper bound for Δ*L*_*k*,*l*_ as long as the effective root position does not leave the ellipsoid ball. When it does leave the ellipsoid ball, we should recalculate the upper bounds. Note that this is not a strict mathematical upper bound, because the values of Δ*L*_*k*,*l*_ could be underestimated by the linear approximation, but this underestimation will be very small because for small ellipsoid regions this linear approximation will be very good. Even stronger, any underestimation will be completely offset by the much larger overestimation we do by taking the maximum in the ellipsoid. Indeed, for calculating the maximum we will assume (see below) that the root position and precision travelled to exactly the right point at the border of a very high-dimensional ellipsoid. Since this will happen only with probability zero, we will in all real cases overestimate the possible Δ*L*_*k*,*l*_.

#### A linear approximation of Δ*L*_*k*,*l*_

To get the derivative of Δ*L*_*k*,*l*_ with respect to 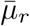 and 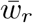, it is convenient to re-write the expression for Δ*L*_*k*,*l*_ using the following identity:

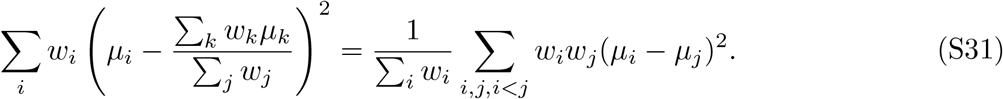

Using this, we get

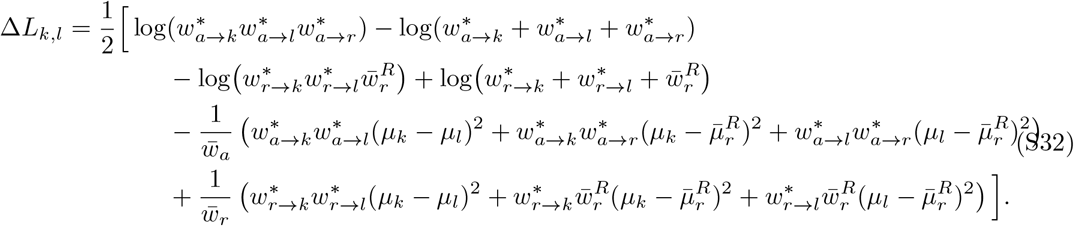

The derivative with respect to 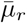 is given by

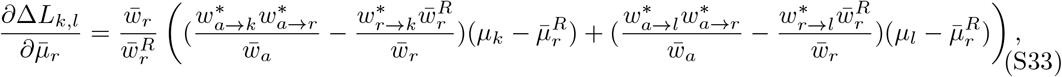

and with respect to 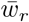 by

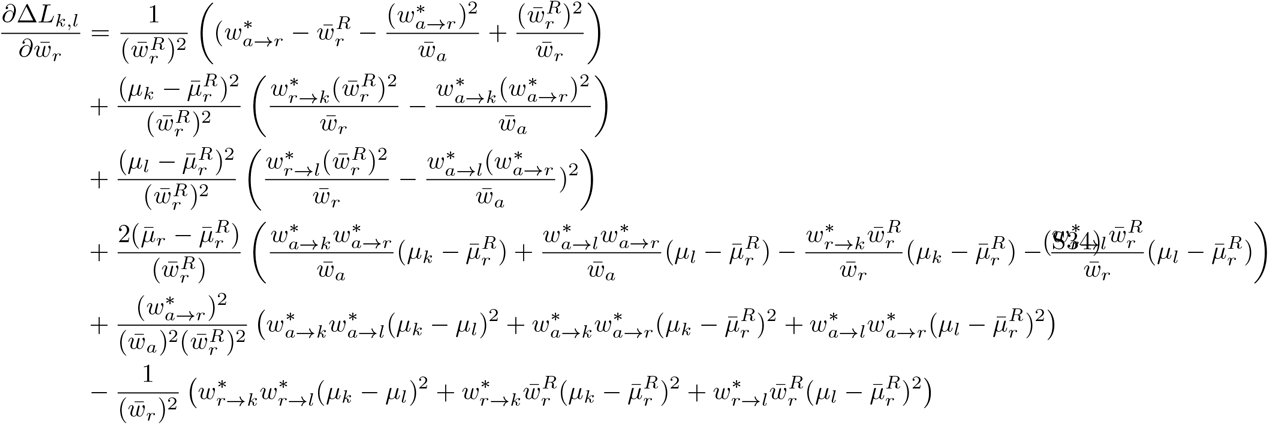

Let us now denote the original position and precision of the root by the usual 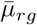 and 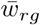, and the movement in these variables by 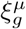 and 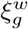. With the above derivatives we then get the linear approximation:

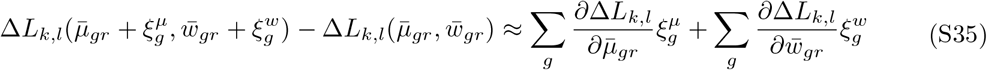

#### Choosing ellipsoid regions around the root position and precision

The above linear approximation allows us to estimate the increase in loglikelihood given a certain root movement 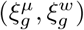. We now want to constrain the allowed movements to a certain ellipsoid around 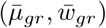 to be able to estimate an upper bound. It is important to note that the validity of the upper bound will not depend on the choice of the ellipsoid. This choice will however determine for how many steps the upper bound will remain valid. We thus want to pick the ellipsoid such that the root position and precision remain within the ellipsoid for as many steps as possible, while still allowing for a tight upper bound.

It is a reasonable choice to assume that the change in root position upon merging two leaves into an ancestral node scales with the standard deviation in the current position of the root and the current number of children of the root (*n*_*c*_), i.e.:

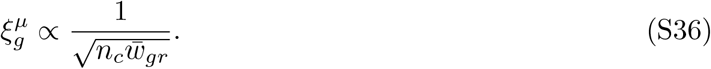

The reasoning behind this is that *x*_*gr*_ is the weighted average of the positions of *n*_*c*_ positions, and we know (when subtracting the mean) that 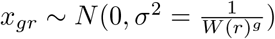. If we make the approximation that the weights are all the same and that the original positions are drawn from some Gaussian, we get that these individual positions must be distributed as 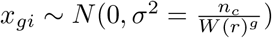. Now, we are taking two of these *x*_*gi*_ out of the average and we add more or less the average of the two. We want to know the distribution of the change:

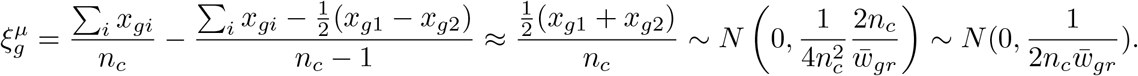

For a typical change in root position, we therefore expect *ξ*^*µ*^ to scale according to (S36). Therefore, we pick the dimensions of the ellipsoid to follow the same scaling.

Similarly, for the change in root precision, a good assumption is that it will scale as

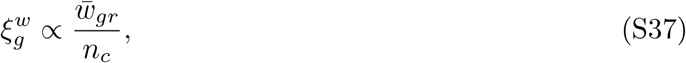

because we know 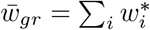. Removing two of these 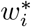 and replacing it by one precision for the ancestor, we can approximate 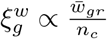.

In addition, we would like the root to remain in the ellipsoid for a certain number of merges (*n*_steps_). We here assume that the root position follows a random walk in each gene-dimension with steps in both the positive and negative direction. This would lead to a scaling of 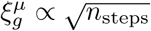. In contrast, the root-precision will only decrease when child-nodes are merged, so there we expect 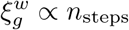. We thus get

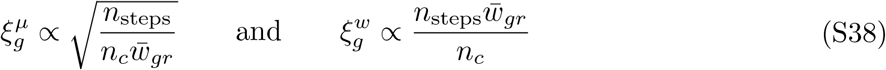

Taking this into account, we choose ellipsoids for the position and precision as

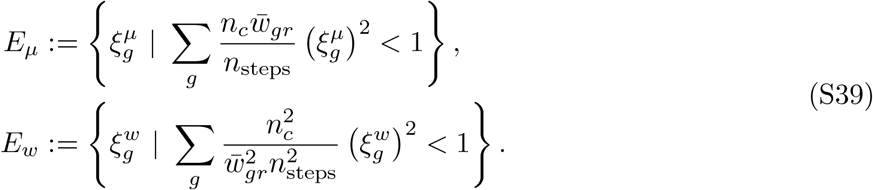

Note that with the *n*_steps_ parameter we can tune the size of the ellipse, and with that, the trade-off between for how many steps the upper bound can be used and how tight the upper bounds are.

#### The final expression for the upper bound

Let us now introduce new variables

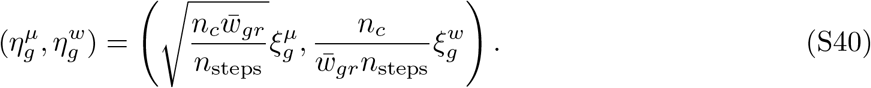

We then get

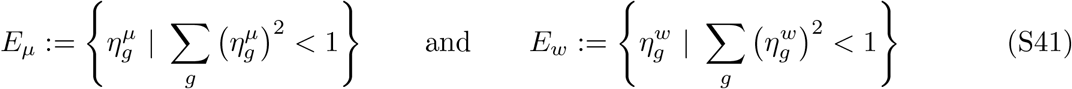

For any 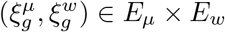 we get that the linear approximation becomes

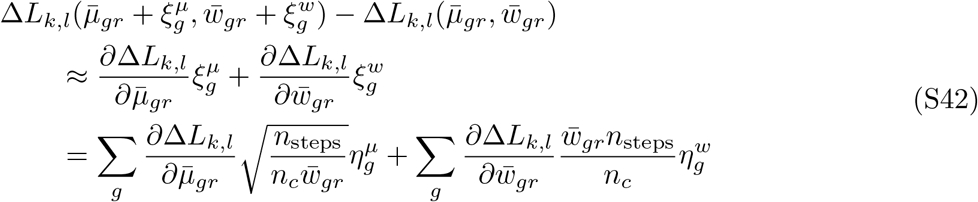

We can now recognize this as a dot product when we define the vectors 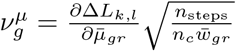 and 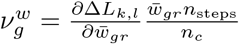. This means we can get an upper bound for the possible loglikelihood increase in the ellipsoid by picking 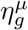 of length 1 and in the direction of 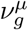, and similarly for 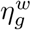. This yields

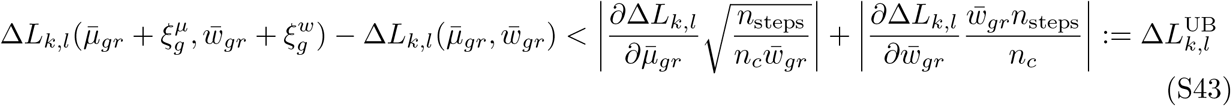

#### Using the upper bound information

To use the upper bound information to speed up the tree reconstruction, we will follow the following steps:

1. We will in the first round of adding an ancestor calculate the maximal loglikelihood increase for all pairs Δ*L*_*k*,*l*_, *and* a corresponding upper bound 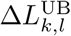 for a given ellipsoid size set by *n*_steps_.
2. To prepare for the next round, we make a list of all pairs, sorted in decreasing order for their upper bound. These upper bounds were thus calculated for the original root-position and -precision: 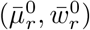.
3. In the next round we calculate the true loglikelihood increase given the new root position and precision: 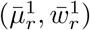 for the first pair in the list. For this pair, we thus estimated the largest upper bound for its loglikelihood increase. We compare its true loglikelihood increase with the upper bound for the second pair in the list. If the loglikelihood increase for the first pair is larger than the upper bound of the second pair, we know that we have found the best pair to merge. We can add the corresponding internal node and move on to the next round. If not, we proceed to calculating the loglikelihood increase for the second pair, and so forth until we have found the best pair.
4. After each addition of an internal node, we need to re-calculate the root position and precision. At that point, we test if these position and precision still fall within the ellipsoid region for which we estimated the upper bounds. If not, we need to calculate new upper bounds in the next round. In addition, for each new internal node, we need to calculate the loglikelihood increase of merging this node to all existing child-nodes. We then also calculate upper bounds for these new pairs and add them to the sorted list.

#### SI.D.1.1 Dynamically setting the ellipsoid size

It is hard to determine the optimal setting of the ellipsoid size (as controlled by the parameter *n*_steps_). Therefore, in the *Bonsai* implementation, we monitor for every round how many pairs we needed to check from the sorted list of upper bounds. If this becomes too large, we decrease the ellipsoid size, as the upper bounds are not sufficiently tight. If the number of checked pairs is small, we increase the ellipsoid size, because we can probably get away with performing fewer calculations of new upper bounds.

### SI.D.2 Limiting the number of candidate-pairs

Although the above described usage of upper bound estimates make sure we do not have to compute |*C*| ^2^ increases in likelihood at every round of adding an internal node, it is still necessary to perform this computation several times, i.e., every time that the root has moved outside of the ellipsoid. It is therefore necessary to further reduce *Bonsai*’s computational footprint to run the largest datasets.

We will here use that, for every round of adding an internal node, we only need to find *the best* pair of nodes to merge into an ancestor, i.e., we don’t need a full ranking of all pairs of nodes. We therefore suggest to limit the search for the best pair by calculating the *k* nearest neighbors for every node based on a simple Euclidean distance in LTQ-space. We have seen in Equation (S32) that the precise loglikelihood increase for merging a certain pair of nodes is a complicated function of the positions and the uncertainties on these positions for the two nodes and the root. However, it is clear that nodes that are close together and far from the root are likely candidates for having a high loglikelihood increase. Correspondingly, a node-pair that is far away from each other is never going to be the best pair to merge in the next round.

In practice, before the first round we calculate the *k* nearest neighbors based on Euclidean distance for every child-node of the root, where *k*-values as low as 5 − 10 have been found to give the optimal tree-reconstruction on simulated datasets. Then we calculate loglikelihood increases only between each node and its nearest neighbors. The best pair of nodes is connected to a new ancestor, and this new ancestor inherits the union of the nearest neighbors of its downstream nodes. The nearest-neighbor relations are re-computed after a certain number of new ancestors has been added (typically after 0.1 |*C*| new ancestors).

It is important to stress that we use the nearest-neighbors only to find the one pair of nodes that is most beneficial to merge, and for that we restrict our search to |*C*| *k* candidates. This is a safe restriction to make, even though it is important to acknowledge that it is generally very hard to capture the full structure of a dataset by calculating a nearest-neighbor graph, and that it is even hard to correctly call nearest neighbors given the typical noisyness of single cell data. In other words, the likelihood that the one best pair in the whole dataset is among the many nearest-neighbors that we calculate is still very close to 1.

### SI.D.3 The effect on *Bonsai*’s computation time

These computational advancements drastically reduce the computational footprint of the *Bonsai* -tree reconstruction. In Figure 3 one can see that *Bonsai* scales sub-quadratically in the number of cells.

## SI.E Creating simulated scRNA-seq datasets

We have tested the performance of *Bonsai* on various simulated datasets. In particular, in Figure 2, and Supplementary Figures S4-S10, we describe the results of *Bonsai* on simulated scRNA-seq data. In general, the creation of such simulated datasets consists of two steps: 1) generating ground truth values, and 2) adding realistic measurement noise to these ground truth values. As explained in Section SI.B.1.2, for scRNA-seq data, the true underlying physical variables are the Log Transcription Quotients (LTQs). Step 1) of simulating the datasets thus consists of generating an LTQ vector for each cell, while step 2) comprises simulating measured mRNA counts given these LTQ values. Whereas the first step depends on the simulator’s choice of what is a reasonable way to mimic a true dataset, there is a principled way of performing the second step. We will therefore first describe the second step, which remains the same across the seven simulated scRNA-seq datasets, and then describe how the LTQs were generated for the different datasets. Note that the simulation scripts can also all be found in the GitHub-repository (https://github.com/dhdegroot/Bonsai-data-representation) in the subfolder paper_figure_scripts_and_notebooks/simulating_datasets.

### SI.E.1 Sampling mRNA-counts given LTQ-values

Remember that the LTQs are defined as

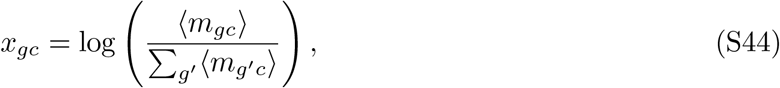

This means that exp 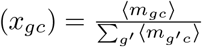. To get a realistic estimate of the sum of expected mRNA-counts that appears in the denominator, we chose to just sample the sum of mRNA-counts per cell from a real dataset, specifically a human pancreatic dataset by Baron et al. [81]. In this dataset, the mean total mRNA-count per cell is 5828, but it varies from 1201 to 34 509. Once we have picked a total count (*N*_*c*_) for each cell and use it as a proxy for 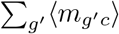 we get *N*_*c*_ exp(*x*_*gc*_) = ⟨*m*_*gc*_⟩, so that now the expected mRNA-number is fixed for each gene and each cell. As mentioned in Section SI.B, and in [28], and as experimentally proven in [76], the true counts are then Poisson samples with these given means. Thus, for every gene-cell pair, we sample a count, *n*_*gc*_, from a Poisson distribution with mean *N*_*c*_ exp{*x*_*gc*_}. These counts *n*_*gc*_ are the data on which we can now test *Bonsai*, and the LTQs *x*_*gc*_ are stored as the corresponding ground truth information about the cells’ gene expression states.

### SI.E.2 Simulating LTQ-values for each cell

We chose to simulate 7 datasets, 6 of which are based on an underlying tree structure, while the other mimics a dataset with 7 strongly separated celltypes. The reason that we based six datasets on a tree structure is that trees provide a way to create non-trivial structure in high-dimensional datasets, while other sampling schemes often result in equidistant data points. The first simulated dataset is based on a binary tree with constant branch lengths, after which we generate more datasets by sampling varying branch lengths, 2) creating an unbalanced tree, 3) enforcing a realistic lower-dimensional covariance structure, or combinations thereof. These simulations can all be reproduced using the script `paper_figure_scripts_and_notebooks/simulating_datasets/simulate_unba lanced_randomtimes_realcovariance.py` that can be found in the *Bonsai-data-representation* GitHub-repository. The basic ingredients for the simulated datasets are:

- All simulated datasets will have 17 499 genes, and 1024 cells.
- For each gene, we pick a random gene from the Baron-dataset [81], and use its corresponding mean log-transcription quotient, here estimated as 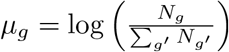.
- In the extensive analysis done in [28], it was noted that the gene variances in the Baron dataset were roughly exponentially distributed with mean 2. For the simulated datasets, we therefore randomly sample a gene variance, *v*_*g*_, from an exponential distribution with mean 2 for each gene.

#### SI.E.2.1 A binary tree with constant branch lengths

To simulate the generation of gene expression states along a binary tree with constant branch lengths, we take the following steps:

1. We assign the zero vector of length 17 499 as the root coordinates: ***y***_root_ = **0**.
2. To generate the coordinates for the two children of the root, we sample a change in gene expression for each gene from a Gaussian distribution with variance *tv*_*g*_, where the branch length *t* is always equal to 1 in this dataset. The two child-nodes of the root are thus given coordinates ***y***_child0_ = ***y***_root_ + ***δ***_0_, and ***y***_child1_ = ***y***_root_ + ***δ***_1_, where the gene expression changes *δ*_0*g*_ are drawn from a Gaussian distribution with variance *v*_*g*_.
3. We repeat the previous step for 10 generations. For example, we generate coordinates for the child-node downstream of child 0 as ***y***_child00_ = ***y***_0_ + ***δ***_00_. As such, we end up with 1024 cells in the last generation. For the simulated dataset, we only take the coordinates of the cells from this last generation.
4. Now that each cell in the dataset is assigned a coordinate vector, we ensure that the means and variances are realistic for gene expression data. For each gene, we center the data by subtracting the mean across cells, after which we rescale the coordinates such that the variance becomes the prescribed variance *v*_*g*_, and add *µ*_*g*_ to match the prescribed mean.
5. Finally, the definition of log-transcription quotients demands that for any LTQ-vector, ***x***, we have ∑_*g*_ exp(*x*_*g*_) = 1. We thus subtract a constant for each cell’s LTQ vector that ensures this.

Given the simulated LTQ-values per cell, we now sample the captured mRNA counts as outlined in Section SI.E.1.

#### SI.E.2.2 Varying branch lengths

For the dataset based on a tree with varying branch lengths, we follow the same steps as for the simple binary tree, except that the gene expression changes are not drawn from a Gaussian distribution with variance 1*v*_*g*_, but with a variance *t*_*i*_*v*_*g*_, where *t*_*i*_ is independently drawn for each new child-node. We draw these branch lengths between 0.5 and 2, and uniformly in log-scale, i.e. log(*t*_*i*_) ∈ Unif[log(0.5), log(2)]

#### SI.E.2.3 An unbalanced tree

To create a dataset based on an unbalanced tree, i.e., a tree with different numbers of leaves on different sides of the tree, we modify the scheme outlined in SI.E.2.1, because we will no longer add two children to every internal node for 10 generations. Instead, we create a list of leaf-nodes, which we initialize with only the root-node. Then, at each step, we take a random leaf-node from the list and make it an internal node by adding two child-nodes downstream of it. These two child-nodes then also replace their parent in the list of leaf-nodes. We repeat this 1023 times, such that 1024 leaves are created.

#### SI.E.2.4 A tree embedded in a lower-dimensional subspace

In two of the datasets, we mimic the covariance structure of a real scRNA-seq dataset. This is to address the possible criticism that the above-created datasets were created by an independent stochastic process in each gene, while in a real dataset, the gene expression changes may occur in a lower-dimensional subspace. This could create a non-trivial covariance structure between the genes, and we want to test with this dataset if *Bonsai* can still reconstruct a faithful representation despite that.

To find a realistic covariance structure between genes, we start by taking the Baron dataset again [81], and we infer the log-transcription quotients (LTQs) for this data using *Sanity* [28]. Since *Sanity* infers a Gaussian approximation of the posterior distribution of the LTQs, we get the most realistic LTQs by sampling from this Gaussian posterior. We therefore sample an LTQ-value for every gene-cell-pair from the posterior distribution, getting realistic LTQs in a matrix *X*_*gc*_. We center *X*_*gc*_ by subtracting the mean LTQ per gene to get 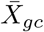, and then perform a singular value decomposition to get matrices *U*_*gk*_, *σ*_*k*_, *V*_*ck*_ that satisfy for all gene-cell-pairs:

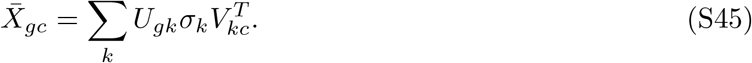

We now want to create an LTQ-matrix 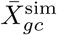 that results from a ground truth tree-structure, and that has a similar covariance structure as 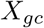. For this, we are using that the columns of *U*_*gk*_ contain the PCA-components in gene-space, and that 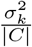 captures the variance along the *k*-th component. In the following, we will keep these two components the same. In contrast, the columns of *V*_*kc*_ contain the coordinates of the cells in the PCA-basis. We will simulate a matrix 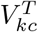 by modeling diffusion along a binary tree. This is thus the same as in Section SI.E.2.1, except that now the diffusion happens in the PCA-basis rather than in the gene-basis. Given the resulting cell-vectors in the PCA basis, we scale each component to have a variance of 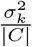, after which we left-multiply with *U*_*gk*_ to rotate the simulated vectors back from the PCA-basis into the gene-basis. The resulting dataset, by construction, has the same covariance structure as the original real dataset.

#### SI.E.2.5 A pseudo-bulk-based dataset

To create a simulated dataset not based on a tree, we start by taking a single-cell dataset of the mouse cortex and hippocampus (Zeisel et al. [82]), of which the cells were annotated into 7 clusters. We create so-called pseudobulked counts by summing all mRNA counts for cells from the same celltype. We thus obtain 7 mRNA count vectors, one for each celltype, on which we run *Sanity* to obtain the corresponding ‘celltype LTQs’. Subsequently, we assign each of the 1024 simulated cells to one of the 7 celltypes in proportion to the abundance of the celltype in the original dataset. The final cell-type abundances range from 34 cells for the rarest to 320 cells for the most abundant celltype. We then create the LTQs for each cell by taking its celltype-LTQ vector and adding a contribution drawn from a Gaussian with variance 0.05. This small amount of intra-celltype variation is to be expected in biological datasets, and ensures that the kNN-test presented in S10 is well-defined. Finally, we again ensure that the LTQs satisfy the constraint ∑_*g*_ exp(*x*_*g*_) = 1, and we sample the mRNA counts as described in SI.E.1.

## SI.F Creating more general simulated datasets

In Section SI.E, we describe how we simulated scRNA-seq datasets including the sampling of mRNA counts. However, we have also used three test datasets in which we directly simulated the input coordinates that were given to *Bonsai*, PCA and UMAP. We will here shortly outline how these data were simulated. The simulation scripts and the necessary commands to run them can also all be found by reading through the Jupyter Notebooks provided in the GitHubrepository (https://github.com/dhdegroot/Bonsai-data-representation) in the subfolder `paper_figure_scripts_and_notebooks`.

### SI.F.1 Noise-free datasets with increasing dimensionality

Using Figure S2, we argued that distances between objects can be captured better and better at increasing dimensionality. For this figure, we simulated a set of datasets with 100 objects in either 2, 10, 100, 1000 or 10 000 dimensions. For each gene, we sample a gene-variance *v*_*g*_ from an exponential distribution with mean 2. Then, for each cell, we draw a cell-variance *t*_*c*_ uniform in log-scale between 0.1 and 10, i.e. log *t* Unif[log(0.1), log(10)], after which we rescale all the *t*_i_ such that 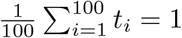. Finally, we just draw the coordinate for object *i* and gene *g* from a Gaussian distribution with mean zero and variance *t*_*i*_*v*_*g*_. These coordinates are stored as the input to *Bonsai*.

### SI.F.2 Noisy datasets with an increasing number of objects

Through Figure S11, we showed that tree-structures can automatically use nearby cells to get more accurate distance estimates in noisy datasets. For this, we will use the 100 points in 100 dimensions from the simulation in SI.F.1 as cluster-centers; these will correspond to the *true positions*. Then, we mimic having noisy readouts from these cluster-centers by creating either 1, 2, 5, 10, or 20 objects around each center, thus creating in total 100, 200, 500, 1000 or 10 000 objects. The *measured coordinates* of these objects are given by the cluster-center-coordinates plus a noise term drawn from a Gaussian with variance 2.5. When *Bonsai* is used to reconstruct a tree and infer the distances, it is given the measured coordinates and the correct size of the measurement noise as input. The inferred object-to-object-distances are compared to the distances between the true positions, i.e., between the cluster-center-coordinates.

To make the task of inferring distances between objects perfectly comparable across the datasets with increasing numbers of objects, we use random seeds. In this way, we make sure that the first object around each cluster-center always has the exact same coordinates in each dataset, that the second object around each cluster-center has the same coordinates in the last 4 datasets, and so on. The distance-comparison presented in Figure S11 is done only on the distances between the first objects around each cluster-center, such that both the *true* coordinates (i.e., the cluster-center-coordinates) and the *measured* coordinates are the same for the ‘test-set’ in the 5 datasets.

## SI.G Tree visualization

Through *Bonsai-scout*, we offer an interactive dashboard with which the *Bonsai* tree can be visualized and explored interactively. We will here shortly discuss some core aspects of the available visualizations.

### SI.G.1 Tree layout algorithms

In *Bonsai-scout*, we offer three layout-types that can be used to plot the tree in two dimensions: the ladderized dendrogram, the equal-angle layout, and the equal-daylight layout. The full algorithms can be found in the *Bonsai* GitHub-repository in the file `bonsai_scout/my_tree_layout.py`.

The *dendrogram layout* may be the most common way to visualize trees and does not require a description. We here choose to ladderize the dendrogram, which means that, from any node, we choose to sort the branches form top-to-bottom based on how many leaves there are below those branches.

The equal-angle and equal-daylight algorithm were proposed by Felsenstein in his excellent book *Inferring phylogenies* on pages 578 to 584 [19]. These are circular layouts that are especially suitable for unrooted trees like the one that *Bonsai* produces. The equal-angle algorithm works well for trees of all sizes, and will never lead to crossing edges. The equal-daylight algorithm can be used to improve the equal-angle tree by more homogeneously distributing the leaves over the circle. Although the equal-daylight algorithm leads to considerably better results for small trees, it can become computationally intense for large trees, and can, in the implementation suggested originally, even lead to crossing edges. We have re-implemented the equal-daylight algorithm to streamline its computation and to avoid crossing edges, but on large trees the benefits are modest. Therefore, we only offer the equal-daylight layout for trees with less than 2000 nodes.

### SI.G.2 The interpretation of tree distances

It is important to note that distances between tree-nodes should always be measured by adding up the branch lengths along the path that connects the nodes on the tree, rather than by measuring the 2-dimensional distance between the positions of the nodes in the figure. This has a number of consequences. First, as long as we keep the tree topology and the branch lengths fixed, we can visualize the tree in different ways without changing the tree’s content. We benefit from this by offering different tree layouts. Second, the tree is independent of the order in which branches are connected to a node. For example, when a node has two downstream nodes, say A, and B, the tree-likelihood is independent of ordering these as AB, or as BA, even though it affects the 2-dimensional visualization. To aid users in gathering intuition for the effects of these meaningless changes, we offer a “Layout tweaking”-tab in *Bonsai-scout*, where one can change the ordering of downstream branches around a selected node.

#### SI.G.2.1 Distances in the dendrogram layout

One of the layouts that is offered in *Bonsai-scout* is the dendrogram layout. In the dendrogram layout, it should be noted that vertical distances are uninformative for the actual distances between tree-nodes. Indeed, in the dendrogram layout, distances should be measured by *only summing horizontal branch lengths* along the path on the tree between the objects.

#### SI.G.2.2 Distances on a disk with hyperbolic geometry

For trees that have large variation in the branch lengths, it can be difficult to visualize the whole tree while still being able to resolve details on a part of the tree. To accommodate this, we project the tree on a hyperbolic disk on which distances near the edge of the disk get more and more compressed. To offer the user some intuition for this, when we visualize a hyperbolic disk, we always include a number of equal-sized squares on the background that appear to get smaller towards the edges of the disk. Branch lengths can thus be visually measured by how many of the equal-sized squares the branch crosses.

Technically, the projection to the hyperbolic disk uses a mapping from the 2-dimensional reals to the 2-dimensional unit ball. The projection is dependent on two parameters, an origin ***x***_0_ = (*x*_0_, *y*_0_) and a zoom *z*. Given the position of any 2-dimensional point (*x, y*), we first subtract the origin, and then scale by the zoom: 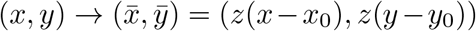. Next, we switch to polar coordinates, (*r, θ*), such that 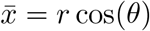, and 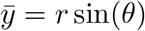. The hyperbolic mapping leaves *θ* unchanged, while it maps *r* to 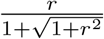. Note that this mapping sends an infinitely large *r* to 1, and the origin (*r* = 0) to itself.

^1^Note that these assumptions still allow for a change in gene expression to be caused by a signal outside of the cell, e.g. caused by the cell’s spatial position, as long as this signal is reflected in the current gene expression.

^2^Note that this assumption still allows for a change in gene expression to be caused by a signal outside of the cell, as long as the change can be predicted from the current gene expression. For instance, a spatial signal may drive changes in gene expression, but as long as the cell’s spatial position is reflected in its current expression then the current state may still be predictive of the changes.

^3^The additional assumption is that the probability distribution of gene expression changes over a very short time interval *δt* has a finite mean and variance that scale proportionally to *δt*.

^4^It is timely to here apologize for the combined usage of a bar and a tilde in this notation.

